# Reorganization of DNA loops by competition between condensin I and a linker histone

**DOI:** 10.1101/2025.04.29.651375

**Authors:** Tetsuya Yamamoto, Keishi Shintomi, Tatsuya Hirano

## Abstract

Condensin-mediated loop extrusion is thought to be one of the primary mechanisms underlying mitotic chromosome assembly. However, how this process is affected by other chromosomal proteins, such as histones, is not well understood. Our previous study showed that in *Xenopus* egg extracts co-depleted of topoisomerase IIα and the histone chaperone Asf1, a highly characteristic chromatin structure called the “sparkler” is assembled. The sparkler is a compact structure assembled on nucleosome-free, entangled DNA in which multiple protrusions radiate from a core. Interestingly, condensin I is concentrated at the tips of the protrusions, whereas the linker histone H1.8 is enriched in the remaining regions of the structure. To understand the biophysical mechanisms underlying sparkler assembly, we construct a model predicting that DNA loops extruded from the entangled DNA undergo phase separation into two domains: loops enriched in condensin I remain as protrusions, whereas those enriched in H1.8 are reeled into the central region. We propose that H1.8 competes with condensin I for DNA binding, thereby reorganizing DNA loops formed by condensin I under this specialized condition.

## INTRODUCTION

Upon entry into mitosis, chromosomal DNA undergoes a series of drastic structural transitions from relatively diffuse interphase chromatin to a discrete set of rod-shaped structures, each composed of duplicated sister chromatids. This process, collectively referred to as mitotic chromosome assembly, is essential for the faithful segregation of genetic information into daughter cells. Over the past two decades, it has been established that condensins, a class of structural maintenance of chromosomes (SMC) protein complexes, play central roles in this process (1). A recent theory, combined with computer simulations, predicted that condensins drive mitotic chromosome assembly by forming successive loops through an active mechanism called loop extrusion (2-6). Indeed, subsequent single-molecule assays supported the loop extrusion theory by demonstrating that budding yeast condensin translocates along DNA and forms DNA loops in an ATP hydrolysis-dependent manner (7,8). Despite these remarkable advances, the mechanism by which condensins extrude loops within crowded DNA environments where they interact with other chromosomal proteins is not well understood.

To understand the mechanisms underlying mitotic chromosome assembly, we have developed and refined a cell-free assay using Xenopus egg extracts, in which mitotic chromosome-like structures are assembled from mouse sperm nuclei (Fig. 1**a**) (9-11). When these nuclei are incubated with egg extracts depleted of the histone chaperone Asf1, individual chromosome-like structures composed of condensin-enriched central axes and fuzzy extended loops – nucleosome-depleted chromosomes, are assembled (Fig. 1**b**). However, when the same nuclei are incubated with egg extracts co-depleted of the histone chaperone Asf1 and topoisomerase IIα (topo IIα), the chromosomal DNA remains entangled and is converted into a highly characteristic structure, called the sparkler, in which multiple protrusions radiate from a DNA-dense core (Fig. 1c). Condensin I is concentrated at the tips of the protrusions, whereas the linker histone H1.8 is concentrated at the remaining regions of the structure. The compaction of DNA is probably driven by multivalent interaction involving H1.8 aberrantly bound to nucleosome-free DNA (10). Such interactions have also been found from other variants of linker histones (12-15). A previous theory sought to explain sparkler assembly as a volume-phase transition of a DNA gel, driven by multivalent interactions between H1.8 and DNA tension generated by condensin I-mediated loop extrusion (15). However, this theory did not address the mechanism of the emergence of protrusions, one of the most prominent features of sparklers.

**Figure 1.**
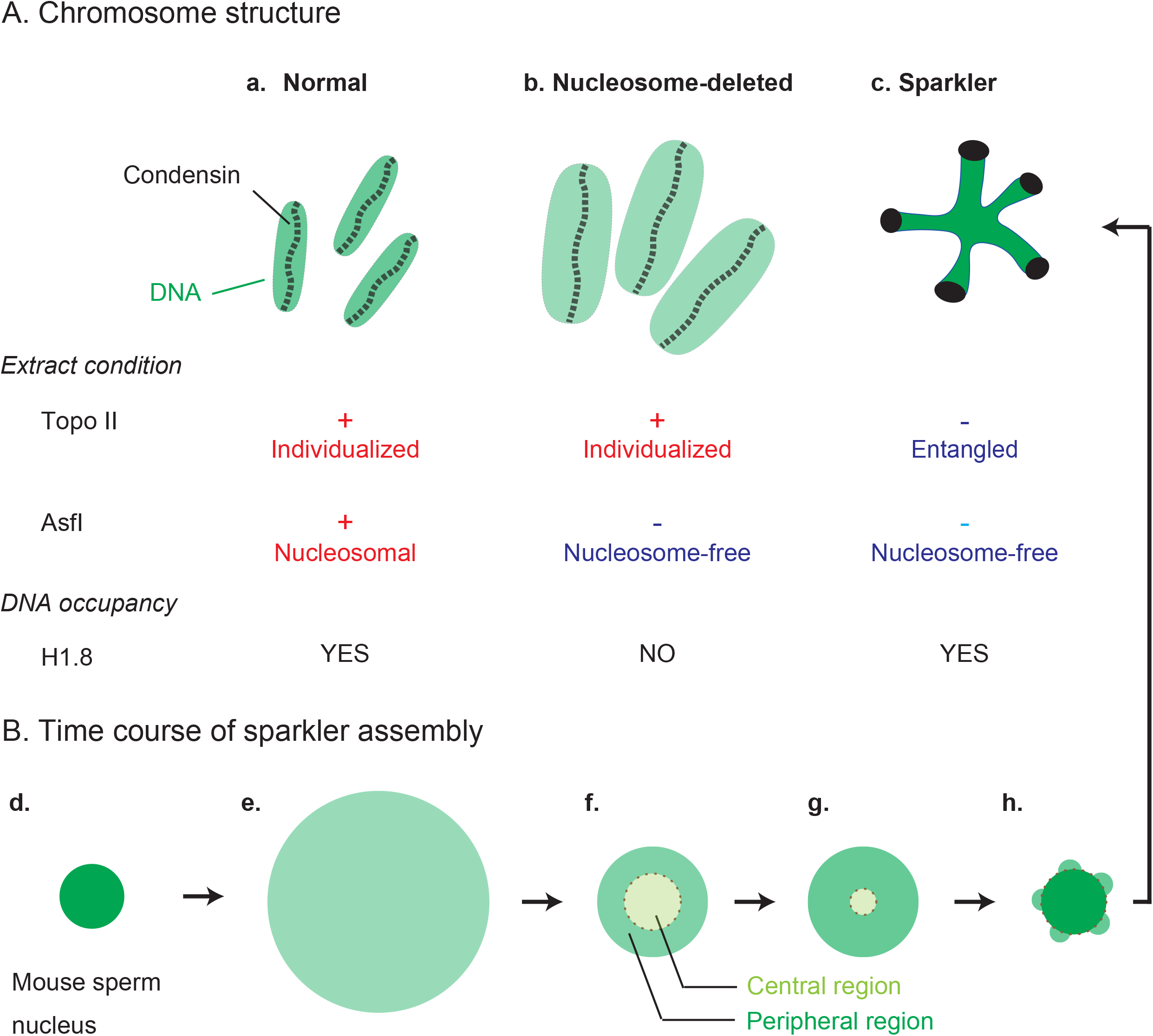
Background information. **(A)** Chromosome/chromatin structures assembled in *Xenopus* egg extracts under three different conditions. In control extracts, a cluster of individual, rod-shaped chromosomes is assembled on nucleosomal DNA (**a**). In extracts depleted of the histone chaperone Asf1, a cluster of individual chromosome-like structures with fuzzy surfaces is formed on nucleosome-free DNA (“nucleosome-depleted” chromosomes; Shintomi et al., 2017) (**b**). In extracts co-depleted of topo IIα and Asf1, no individual chromosomes are observed, resulting in the formation of a highly characteristic chromatin structure (“sparkler”) on nucleosome-free, entangled DNA (Shintomi and Hirano, 2021) (**c**). Although the linker histone H1.8 is undetectable in the nucleosome-depleted chromosomes (**b**), it is abundantly present in the sparkler (**c**). **(B)** Time course of sparkler formation (Shintomi and Hirano, 2021). When mouse sperm nuclei (**d**) are incubated with *Xenopus egg* extracts co-depleted of topo IIα and Asf1, entangled chromosomal DNA strands within the nuclei rapidly expand to form a cloud-like structure (**e**). The cloud-like structure gradually shrinks to form a bipartite architecture composed of central and peripheral regions (**f**). At this stage, the concentrations of DNA and condensin I are higher in the peripheral region than in the central region. As the DNA volume fraction in the peripheral region increases (**g**), apparent symmetry breaking occurs, resulting in the appearance of protrusions in the peripheral region (**h**). Simultaneously, the entire structure becomes more compact. In this final structure of sparklers (**c**), condensin I is concentrated at the tips of the protrusions where DNA is sparse, while H1.8 is enriched in the other regions where DNA is dense.

The time course of sparkler assembly provides further insight (10) (Fig. 1**d-h**). Mouse sperm DNA strands incubated in an extract co-depeted of Asf1 and topo IIα (Fig. 1**d**) swell into a uniform cloud-like structure without dissolving into the external solution over the experimental time scale (Fig. 1**e)**. The entangled DNA strands are subsequently reorganized into a spherical structure, in which both DNA and condensin I are more dilute in the central region than in the peripheral region (Fig. 1**f**). The thickness of the peripheral region increases over time, whereas the size of the central region decreases (Fig. 1**g**). Subsequently, the bulk of the DNA is lost from the peripheral region, leaving multiple DNA domains behind: the spherical symmetry of the system is broken (Fig. 1**h**). The DNA lost from the peripheral region is reeled into the central region, increasing its size and DNA concentration. Subsequently, the entangled DNA strands adopt a star-like structure, with DNA domains remaining in the peripheral region as protrusions (Fig. 1**c**). This series of events implies that the emergence of protrusions is driven by symmetry breaking.

Chromosomal DNA molecules present in the mouse sperm nuclei are entangled and remain entangled when the nuclei are exposed to Xenopus egg extracts depleted of topo IIα. Here, we construct a theoretical model to study the biophysical mechanism of symmetry breaking during sparkler assembly by taking into account DNA entanglement. In polymer physics, entanglement is treated as effective crosslinks; entangled DNA strands act as a polymer gel (16,17). Unlike chemical crosslinks, DNA strands can slide through effective crosslinks (18-21). Our theory treats the central region as an entangled DNA gel, and the peripheral region as a brush of DNA loops extruded by condensin I. Our theory predicts that the radius of the central gel region decreases over time to relieve the DNA tension generated by the loop extrusion process. This in turn increases the DNA concentration in the periphery and promotes H1.8 binding. When the DNA concentration becomes sufficiently high, the peripheral region undergoes phase separation into domains of swollen and condensed DNA loops. Swollen DNA loops enriched in condensin I are stabilized by continuous condensin I-loading and remain in the peripheral region. In contrast, condensed DNA loops enriched in H1.8 resist condensin I loading and are consequently reeled into the central gel region. Therefore, the phase separation leads to the breaking of spherical symmetry in the system. Our theory reveals a linker histone as a regulator of DNA loop stability extruded by condensin I.

## MATERIALS AND METHODS

### Model overview

We treat entangled DNA strands in a solution containing linker histone and condensin (Fig. 2). The DNA strands act as electrically neutral flexible polymers composed of Kuhn units of length *b* because their electric charges are neutralized at physiological salt concentrations (22). The chromosomal DNA strands are extremely long and therefore remain entangled over the experimental timescale (23). We treat DNA entanglement as effective crosslinks, where the DNA strands can slide across crosslinks (18-21). Linker histone and condensin bind to the DNA strands in a mutually exclusive manner (10,24). We treated linker histone bound to DNA as factors that give rise to the multivalent attractive interaction and condensin bound to DNA as the loop extrusion factor.

**Figure 2.**
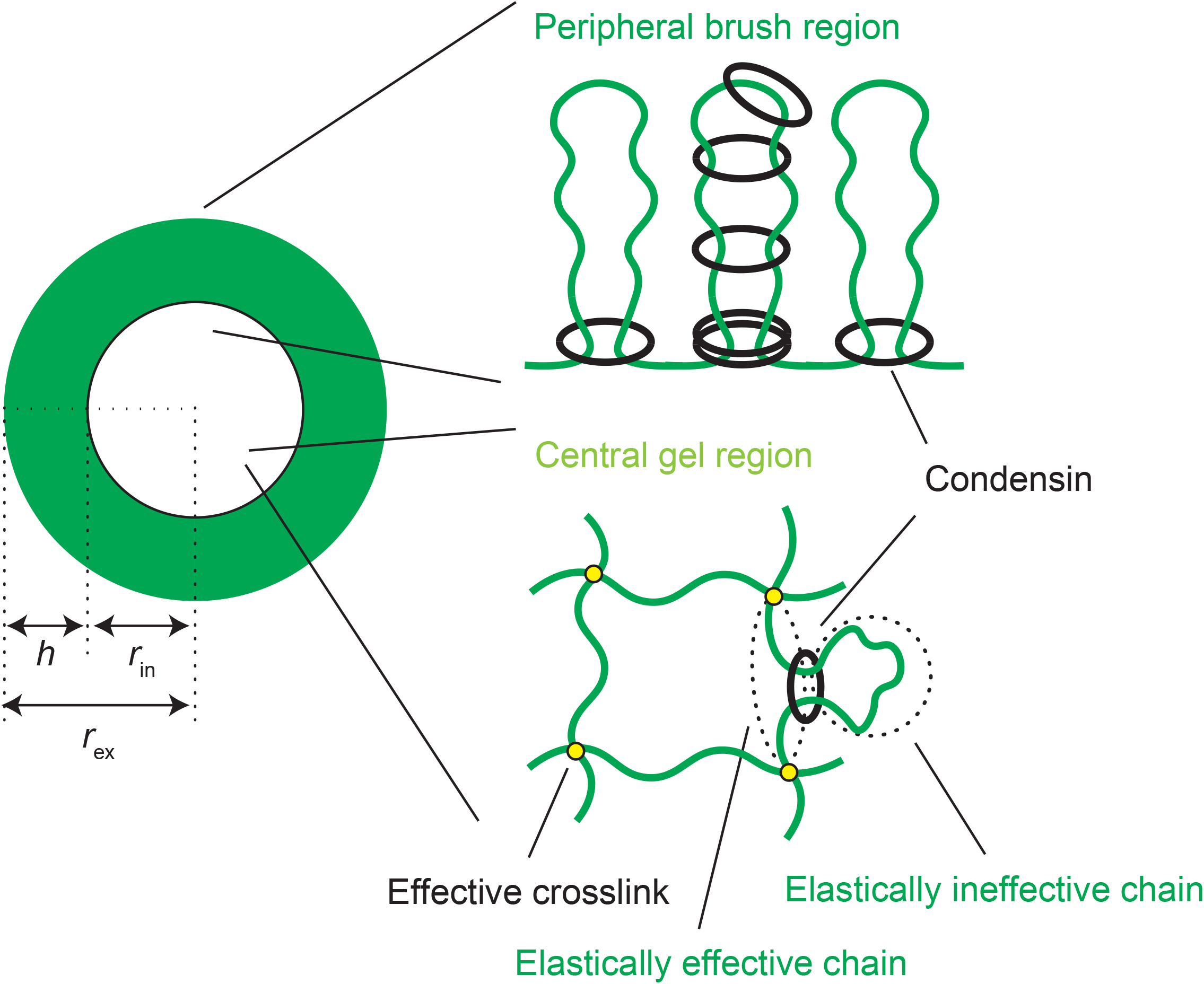
Entangled DNA model. The center is modeled as an entangled DNA gel and the periphery is modeled as a brush of DNA loops extruded by condensin, which also extrudes loops within the central gel. These loops are elastically ineffective, while other parts of the entangled DNA network are elastically effective.

The central region of a sparkler intermediate is treated as a spherical, entangled DNA gel, whereas the peripheral region is treated as a brush of DNA loops extruded from the central region by condensin I. While condensin I also generates loops in the central region, these loops do not respond to deformation of the DNA gel and are therefore considered elastically ineffective (17,25). In contrast, the remaining DNA in the central region responds to gel deformation and is considered elastically effective. The numbers of DNA units in each region were determined by loop extrusion dynamics, while the total number *N* of DNA units in the system is conserved

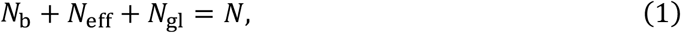

where *N*_b_, *N*_eff_, and *N*_gl_ are the numbers of DNA units in the peripheral, elastically effective chains, and elastically ineffective chains, respectively.

We analyze both cases in which dynamics is limited by loop extrusion and relaxation. Relaxation of polymer gels and brushes is governed by solvent flow (26,27). The structures of the central and peripheral regions are represented by the volume fraction *ϕ* of DNA, the occupancy *α*_h_ by linker histone, and the occupancy *α*_c_ by condensin. The subscripts `c’ and `h’ indicate condensin and linker histone, respectively.

### Free energy

The free energy of the system is given by the sum of the free-energy contributions from the central and peripheral regions,

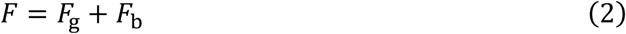

The subscripts `g’ and `b’ indicate the central gel and peripheral brush regions, respectively. The free energy is a functional of the occupancy *α*_c_ by condensin, the occupancy *α*_h_ by linker histone, and the DNA volume fraction *ϕ*. The free energy of the central gel region has the form

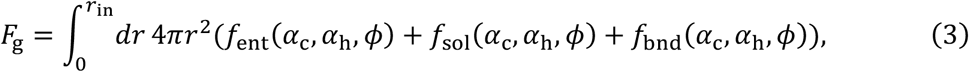

where *f*_ent_(*α*_c_, *α*_h_, *ϕ*) is the elastic free energy due to entanglement, *f*_sol_(*α*_c_, *α*_h_, *ϕ*) is the solution free energy, and *f*_bnd_(*α*_c_, *α*_h_, *ϕ*) is the binding free energy, all per unit volume. *r*_in_ is the radius of the central region. The free energy of the peripheral brush region has the form

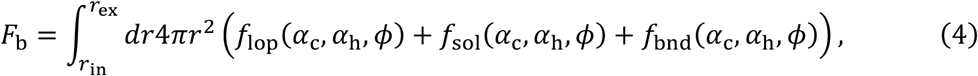

where *f*_lop_(*α*_c_, *α*_h_, *ϕ*) is the elastic free energy per unit volume due to stretching of DNA loops. *r*_ex_ is the radius of the entire DNA structure, including both the central and peripheral regions. The free-energy contributions *F*_g_ and *F*_b_ are designed to take into account the features of condensin and linker histone by extending the Flory-Rehner theory of polymer gels (28) and the Alexander-de Gennes theory of polymer brushes (29,30), respectively, with an extension based on asymptotic analysis of the Daoud-Cotton theory (31) (see also sec. S1 and Fig. S1 in SI).

The elastic free energy *f*_ent_(*α*_c_, *α*_h_, *ϕ*) per unit volume due to entanglement has the form

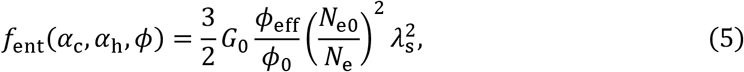

which represents the elastic free energy stored by isotropic deformation of the entangled DNA network by the swelling ratio *λ*_s_ from the reference state (32). The reference state is defined as the most relaxed state (15). Eq. (5) is an extension of the elastic free energy of the affine tube model, the simplest model of entanglement (17,25). The derivation of eq. (5) is provided previously (32). *ϕ*_eff_ is the volume fraction of elastically effective chains and *ϕ*_0_ is the DNA volume fraction in the reference state. *N*_e0_ and *N*_e_ are the numbers of DNA units in a subchain between the effective crosslinks in the reference state and after deformation, respectively. The ratio *N*_e_/*N*_e0_ is equal to the fraction *N*_eff_/*N* of DNA units in the elastically effective chains. *G*_0_ (= *ϕ*_0_*k*_B_*T*/(*b*^3^*N*_e0_)) is the shear modulus in the reference state, where *k*_B_ is the Boltzmann constant and *T* is the absolute temperature. The volume fraction *ϕ*_eff_ of the elastically effective chains has the relationship

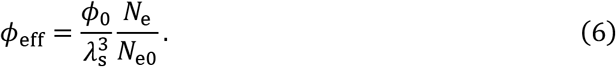

The radius *r*_in_ (= *λ*_s_*r*_R_) is a function of the swelling ratio *λ*_s_, where *r*_R_ is the radius of the entire DNA structure in the reference state.

The solution free energy per unit volume has the form (17,27,33)

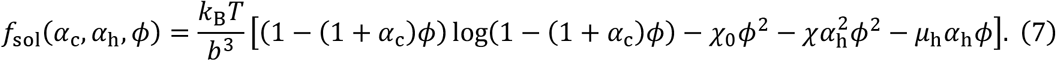

The first term of eq. (7) is the mixing free energy, which represents the random motion of the solvent molecules. The second term of eq. (7) is the DNA-DNA interaction and the parameter *χ*_0_ is the ratio of the magnitude of this interaction to the thermal energy *k*_B_*T*. The third term of eq. (7) is the linker histone-linker histone multivalent interaction and the parameter *χ* is the ratio of the magnitude of this interaction to the thermal energy *k*_B_*T*. The linker histone-DNA interaction is treated separately (eq. (8)). The fourth term of eq. (7) is the contribution of the chemical potential *μ*_h_*k*_B_*T* of linker histone. For simplicity, we assumed that the volumes of DNA units, condensin, and solvents are equal. We neglected the contribution of linker histone to the mixing free energy because their volume is much smaller than the volumes of DNA units and condensin.

The binding free energy per unit volume has the form

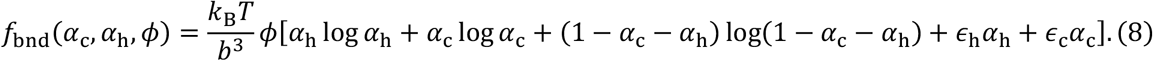

The first, second, and third terms in eq. (8) are the entropic contributions, and the fourth and fifth terms are the energetic contributions with respect to the binding and unbinding of the linker histone and condensin. *k*_B_*Tϵ*_h_ and *k*_B_*Tϵ*_c_ are the binding energy of linker histone and condensin to DNA units, respectively.

The elastic free energy per unit volume in the brush region has the form (29)

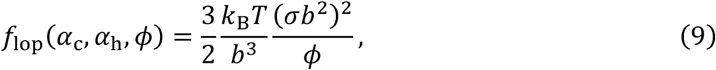

where 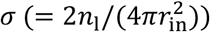 is the number density of DNA loops in the peripheral brush region and *n*_1_ is the number of loops in this region. The number density *σ* depends on the swelling ratio *λ*_s_ in the central region via *r*_in_ ; this represents the connectivity between this and the peripheral regions. The elastic energy in the peripheral region results from the stretching of DNA loops due to the excluded volume interaction between neighboring DNA loops rather than entanglement.

In this paper, we take into account the curvature of the interface between the central and peripheral regions only minimally (Sec. S1 and Fig. S1 in SI). With this treatment, the DNA volume fraction is uniform in both of the central and peripheral regions, which greatly simplifying the model. This treatment is effective for the case in which the brush height *h* is smaller than the radius *r*_in_ and, otherwise, it is only an approximation. The DNA volume fraction *ϕ*_g_ in the central region has the form

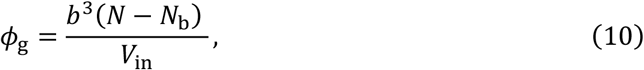

where 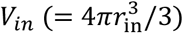 is the volume of the central region. The volume fraction *ϕ*_b_ in the peripheral region has the form

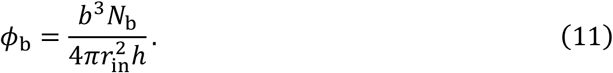

### Chemical potential of linker histone

The minimization of the free energy *F* with respect to the occupancy *α*_h_ leads to the equality of the chemical potential of linker histone,

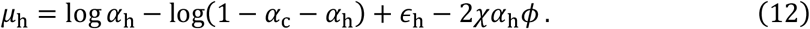

The derivation of eq. (11) is shown in Sec. S2.3 in SI. Eq. (12) is composed of the contributions of entropy (first and second terms), binding energy (third term), and linker histone-linker histone multivalent interaction (fourth term). It can be applied to both of the central and peripheral regions.

### Osmotic pressure

The minimization of the free energy *F* with respect to the DNA volume fraction *ϕ*_g_ leads to the osmotic pressure in the central gel region,

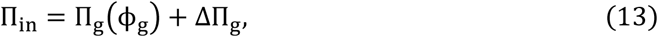

where the second term results from the fact that the free energy of the peripheral region depends on the DNA volume fraction *ϕ*_g_ in the central region via the radius *r*_in_. The derivation of eq. (13) is shown in Sec. S2.6 in SI. The first term is the excess osmotic pressure Π_g_(*ϕ*_g_) in the central region and the second term is the osmotic pressure ΔΠ_g_ that results from the lateral osmotic pressure in the peripheral region. The excess osmotic pressure Π_g_(*ϕ*_g_) has the form

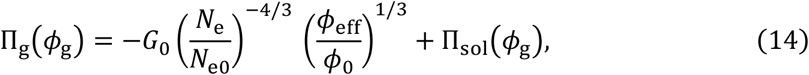

where the first term is the elastic stress of the entangled DNA network and the second term is the osmotic pressure of the solution

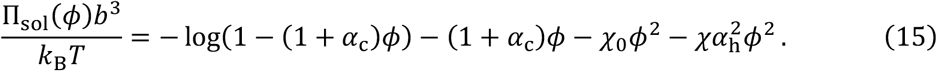

The osmotic pressure contribution ΔΠ_g_ has the form

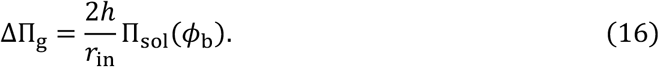

for *h* < *r*_in_/2 . Eq. (16) represents the projection of the lateral osmotic pressure *h*Π_sol_(*ϕ*_b_) in the radial direction due to the curvature 2/*r*_in_ of the interface. For *h* > *r*_in_/2, we use

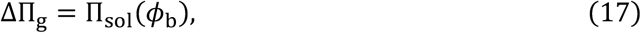

(Sec. S1 and Fig. S1 in SI). The osmotic pressure Π_in_ is balanced by the hydrostatic pressures *p*_in_ of the solvent

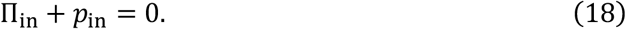

The minimization of the free energy *F* with respect to the DNA volume fraction *ϕ*_b_ in the peripheral region leads to the form

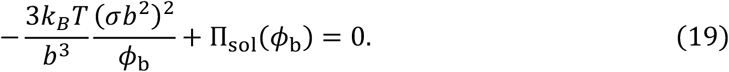

The derivation of eq. (19) is shown in Sec. 2.5 in SI. This equation represents the balance between the elastic stress of the DNA loops (first term) and the osmotic pressures (second term) in the peripheral region. For simplicity, we neglect the contribution of the solvent flow to the conformation of DNA.

### Loop extrusion rate and DNA tension

Single-molecule experiments have shown that the extrusion rate *v* decreases with increasing DNA tension *f*. For simplicity, we assumed the linear dependence of the extrusion rate

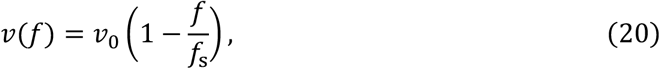

for *f* < *f*_s_ and *v*(*f*) = 0 for *f* > *f*_s_, motivated by the treatment of other molecular motors (34). *f*_s_ is the stall force of condensin and *v*_0_ is the maximum extrusion rate.

Loop extrusion produces tension in the elastically effective chains in the central region because it decreases the number of DNA units in the subchains between effective crosslinks. The relaxation of polymer gels and brushes is governed by solvent flow (26). If DNA translocation by condensin is faster than relaxation, the volumes *V*_in_ and *V*_ex_ of the central and peripheral regions are constant. The DNA tension *f*_ex_, which is overcome by condensin to extrude DNA units towards the peripheral region,, is derived by using the principle of virtual displacement as

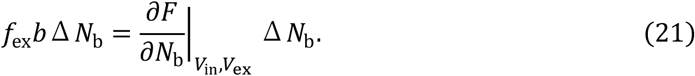

Eq. (21) represents that the work done by extruding Δ*N*_b_ DNA units from the central to the peripheral regions (left side) is equal to the variation of the free energy due to this process with fixed volume (right side). Δ*N*_b_ (= 2*v*Δ*tn*_1_) is the total number of DNA units extruded by *n*_1_ condensin complexes at the interface between central and peripheral regions during the time period Δ*t*. Thus, DNA tension has the form

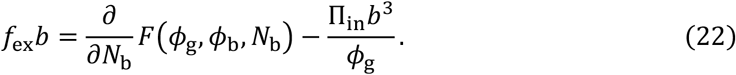

The second term of eq. (22) results from the fact that the DNA volume fraction *ϕ*_g_ depends on *N*_b_.

Similarly, DNA tension, which is overcome by condensin to extrude DNA in the elastically effective part to the elastically ineffective part, is derived as

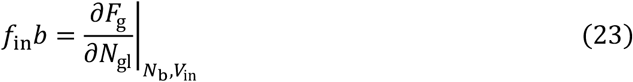

because the number of DNA units in the central region does not change with this process.

The expressions for *f*_ex_ and *f*_in_ are shown in Secs. S2.7 and S2.8 in SI. Extrusion from the elastically effective to ineffective regions does not change the DNA volume fraction *ϕ*_g_ in the center; thus, the DNA tension *f*_in_ results solely from elastic force produced by entanglement. In contrast, extrusion from the central to the peripheral regions decreases the excluded volume interaction in the central region; this slightly reduces the DNA tension *f*_ex_ . Note that *f*_ex_ and *f*_in_ are instantaneous DNA tensions during extrusion steps before the DNA is relaxed.

### Loop extrusion dynamics

The time evolution equation of the number *N*_b1_ (= *N*_b_/*n*_1_) of DNA units in a loop in the periphery has the form:

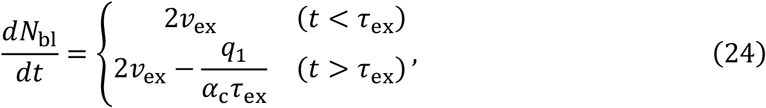

where *v*_ex_(= *v*(*f*_ex_)) is the extrusion rate of the condensin at the interface between the central and peripheral regions, and *q*_1_ is the probability that the number of condensin at the interface is unity.

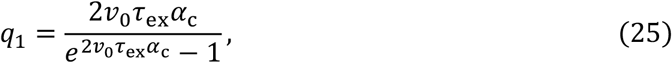

because condensin arrives at the interface with rate 2*v*_0_*α*_c_ due to the loop extrusion and are unloaded with rate 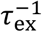 (Sec. S3.1 in SI). The first term of eq. (24) is the extrusion rate of DNA by the condensin at the interface. The second term of eq. (24) for *t* > *τ*_ex_ represents that the 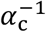 DNA units between condensin complexes are released towards the center by the unloading of all the condensin complexes at a loop anchor with rate *q*_1_/*τ*_ex_. We neglect the loop branching because the simultaneous loading of multiple condensin complexes is a rare event (4).

For simplicity, we considered cases in which each elastically ineffective loop in the central region is produced by at most one condensin. The time evolution equation of the number *N*_gl_ of DNA units in the elastically ineffective chains has the form:

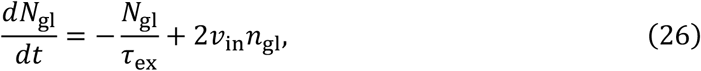

where *v*_in_ (= *v*(*f*_in_)) is the extrusion rate of condensin that form elastically ineffective loops in the central region and *n*_gl_ is the number of such loops. The time evolution of the number of elastically ineffective loops has the form

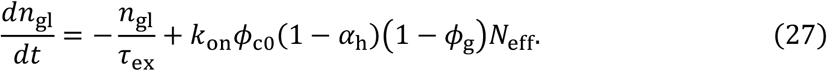

Eq. (27) is an extension of the kinetic equation for binding and unbinding (35). The solution of eqs. (26) and (27) has the form

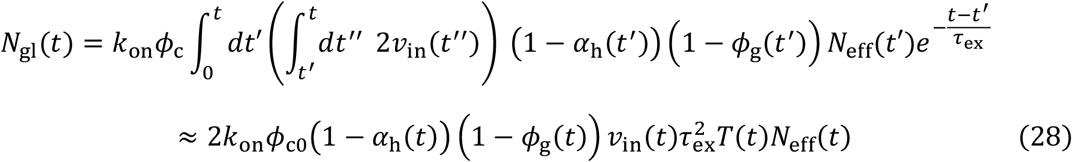

with the time factor

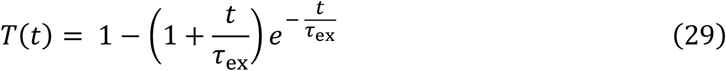

To derive the last form of eq. (28), we assumed that the functions *v*_in_, *α*_h_, *ϕ*_g_, and *N*_eff_, change slower than 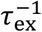 (Sec. S3.2 in SI). This assumption is effective for *t* > *τ* and is only an approximate treatment for *t* < *τ*_ex_.

### Condensin loading/unloading kinetics

Both topological loading and loop extrusion are driven by ATP hydrolysis. The time evolution equation of the occupancy *α*_c_ by condensin in the central gel has the form

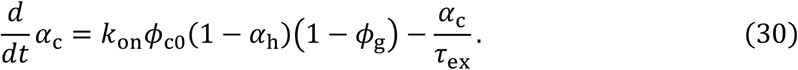

The first term of eq. (30) is the loading rate of a condensin and *k*_on_ is the rate constant that accounts for this process. The second term of eq. (28) is the unloading rate of condensin. *ϕ*_c0_ is the volume fraction of condensin in the external solution. The factors (1 − *α*_h_) and (1 − *ϕ*_g_) in the first term of eq. (30) accounts for the suppression of the loading of a condensin by linker histones and the osmotic pressure, respectively (Secs. S2.2 and S2.4 in SI).

The time evolution of occupancy *α*_c_ in the loop brush region has the form

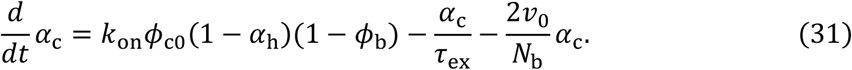

The first term of eq. (31) is the loading rate and the second term is the unloading rate of a condensin. The third term of eq. (31), is the rate at which a condensin is translocated to the interface between the central and peripheral regions via loop extrusion.

To derive eqs. (30) and (31), we assume that condensin complexes loaded at the peripheral region do not penetrate the central region, motivated by the fact that condensin is most abundant at the interface during the assembly of the peripheral region is assembled. We derive the occupancy *α*_c_ by condensin for the steady state, 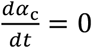.

### Relaxation dynamics

Because the DNA volume fraction is larger in the peripheral region than in the central region during sparkler assembly, we assume that relaxation is governed by solvent flow across the peripheral region. For cases in which the loop extrusion process is faster than the relaxation, the time evolution of the volume of the central region is given by (26,27,36)

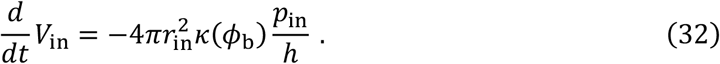

The solvent flow is proportional to the pressure gradient *p*_in_/*h* with the Darcy coefficient *κ*(*ϕ*_b_). The mesh size of the peripheral brush region is given by the average distance *σ*^−1/2^ between the loop. We thus use the Darcy coefficient of the form (26,27,36)

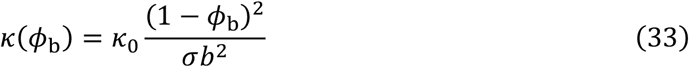

by using a constant *κ*_0_ (≈ *b*^2^/*η*) inversely proportional to the solvent viscosity *η* (36).

### Phase separation in the peripheral region

In general, eq. (19) has two stable solutions, where the volume fraction of one solution (swollen loops) is smaller than that of the other (condensed loops). When the peripheral region is composed of coexisting domains of such loops, the free energy of the peripheral has the form (27,37)

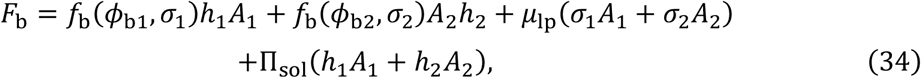

where *f*_b_(*ϕ*_b_, *σ*) (= *f*_lop_(*α*_c_, *α*_h_, *ϕ*_b_, *σ*) + *f*_sol_(*α*_c_, *α*_h_, *ϕ*_b_) + *f*_bnd_(*α*_c_, *α*_h_, *ϕ*_b_)) is the free energy per unit volume in the peripheral region. For simplicity, we omitted the explicit description of the dependence of the free energy *f*_b_ on the occupancies *α*_c_ and *α*_h_. The subscripts 1 and 2 indicate the values of the swollen and condensed loops, respectively. *A*_1_ and *A*_2_ are the areas occupied by the domains of swollen and condensed loops, respectively. The first and second terms in eq. (34) are the free energy contributions from the domains of the swollen and condensed loops, respectively. The third term is the Lagrange multiplayer to ensure that numbers of loops are constant and the fourth term is the contribution of osmotic pressure.

Minimizing the free energy with respect to *σ*_1_, *σ*_2_, *A*_1_, and *A*_2_ leads to the relationships

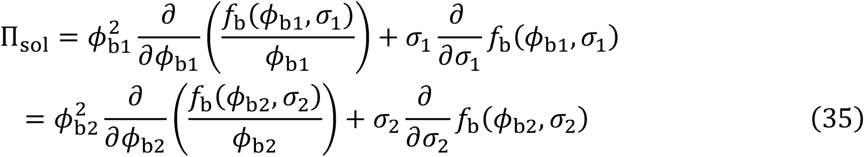

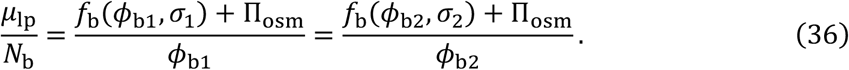

Eqs. (35) and (36) represent the balance of lateral osmotic pressure and chemical potential of the loops between the domains of the swollen and condensed loops (see also eq. (14)). In general, these domains are not at the thermodynamic equilibrium because ATP hydrolysis is involved in the loading and loop extrusion by condensin. Our treatment here is thus only an approximation of phase separation in the peripheral brush region.

### Parameter estimates

As our model considers both loop extrusion and relaxation dynamics in a system composed of a central entangled gel region and a peripheral brush region, the number of parameters involved is rather large. Our model is described by 10 parameters: the rescaled shear modulus 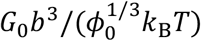, the interaction parameters *χ*_0_ and *χ*_*m*_, the binding free energy, *ϵ*_c_ − *μ*_c_ and *ϵ*_h_ − *μ*_h_, the number of extruded DNA units 2*v*_0_*τ*_ex_, the rescaled stall force *f*_s_*b*/(*k*_B_*T*) of condensin, the occupancy *α*_c0_ (= *k*_on_*τ*_ex_) by condensin for *ϕ* = 0 and *α*_h_ = 0, and combinations of parameters

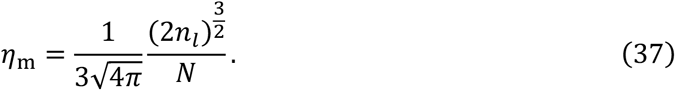

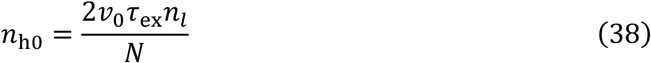

Equation (37) is the ratio of the volume 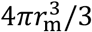, of the central gel region to the total volume *Nb*^3^ of the system when all the solvent in the system is excluded, 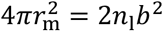, and has been used to describe a system of similar geometry (32). The radius *r*_0_ of the dried DNA strands,

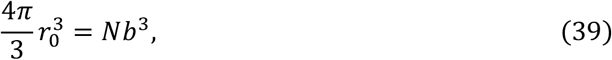

represents the length of the system. Eq. (38) is the ratio of DNA units extruded during time period *τ*_ex_ to the total *N* of DNA units in the system. The parameters used in numerical calculations are listed in Table 1.

**Table 1.** Parameter values.

| Parameter | Physical meaning | Estimate | Used |
| --- | --- | --- | --- |
| $G_0 b^3 / (\phi_0^{\frac{1}{3}} k_B T)$ | Shear modulus | NA | 0.01 |
| $\chi_0$ | DNA-DNA interaction | 0.44 | 0.5 |
| $\chi_m$ | Histone-histone interaction | NA | 2.8 |
| $\epsilon_c - \mu_c$ | DNA-condensin binding energy | NA | 0.0 |
| $\epsilon_h - \mu_h$ | DNA-histone binding energy | NA | 1.9 |
| $2v_0\tau_{ex}$ | Extruded DNA length | 130 | 100 |
| $f_s b / (k_B T)$ | Stall force | 10 | 10 |
| $\eta_m$ | Ratio of bulk and interface | 0.05 | 0.01 |
| $n_{h0}$ | Fraction of DNA units in loops | 0.3 | 0.3 |
| $\alpha_{c0}$ | Condensin occupancy | 0.03-0.3 | 0.3 |

At physiological salt concentrations, the length *b* of each DNA unit is 100 nm (∼ 300 bp) (22). The excluded volume 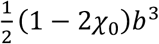 of DNA units is ∼ 0.06 *b*^3^ (38). With this small excluded volume, the correlation is not significant and our mean field treatment is effective (39). The values of the rescaled shear modulus *G*_0_*b*^3^/(*k*_B_*T*), interaction parameter *χ*_m_, and binding energies, *ϵ*_c_ and *ϵ*_h_, are not available. Thus, we set their values to be large enough for linker histone to condense the DNA strands (15).

Single-molecule experiments suggest that yeast condensin extrudes DNA 2*vτ*_ex_ by 40 kbps (≈ 130 DNA units) with a stall force of *f*_s_∼0.1 pN (8). The number of condensin loaded on DNA is estimated as 1 per 1–10 kbps (40,41), which amounts to *α*_c0_. ∼ 0.03– 0.3. The time scale *τ*_s_, with which condensin extrudes DNA units towards the steady state, is estimated as 2*v*_0_*n*_*l*_*τ*_s_ ≈ *N*. By using the experimental time scale *τ*_s_ ∼ 30 min (10) and the extrusion time *τ*_ex_ ∼ 10 min of budding yeast condensin (8), the ratio *n*_h0_ is estimated as 0.3. The number of DNA loops in the peripheral region is *n*_1_ ∼ 3 x 10^4^, because the number of units in mouse sperm DNA (∼ 3 Gbps) is *N* ∼ 1 x 10^7^ and 2*v*_0_*τ*_ex_ ∼ 100 (Eq. (36)). By using these values, the ratio *η*_m_ is estimated to be 0.05 (Eq. (35)). Table 1 Parameter values.

## RESULTS

### Relaxation of the central gel region drives phase separation of the peripheral brush region

We first treat cases in which relaxation caused by solvent flow is faster than loop extrusion, 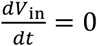. Our model predicts DNA fractions in the central and peripheral regions, DNA volume fraction, condensin occupancy, and linker histone occupancy over time (Fig. S2 in SI). The fraction *N*_b_/*N* of DNA units in the peripheral region increases with time (solid line, Fig. 3**a**). DNA tension is greatly reduced by relaxation and does not significantly decelerate loop extrusion of condensin because of the relatively large stall force of condensin (solid and dotted lines, Fig. 3**a**). Solvent flows out from the central region to relax the DNA tension; thus, the radius *r*_in_ of the central gel region decreases with time (red line, Fig. 3**b**). This decreases the distance between DNA loops in the peripheral region. The DNA volume fraction *ϕ*_g_ in the peripheral region thus increases over time (dark green line, Fig. 3**c**).

**Figure 3.**
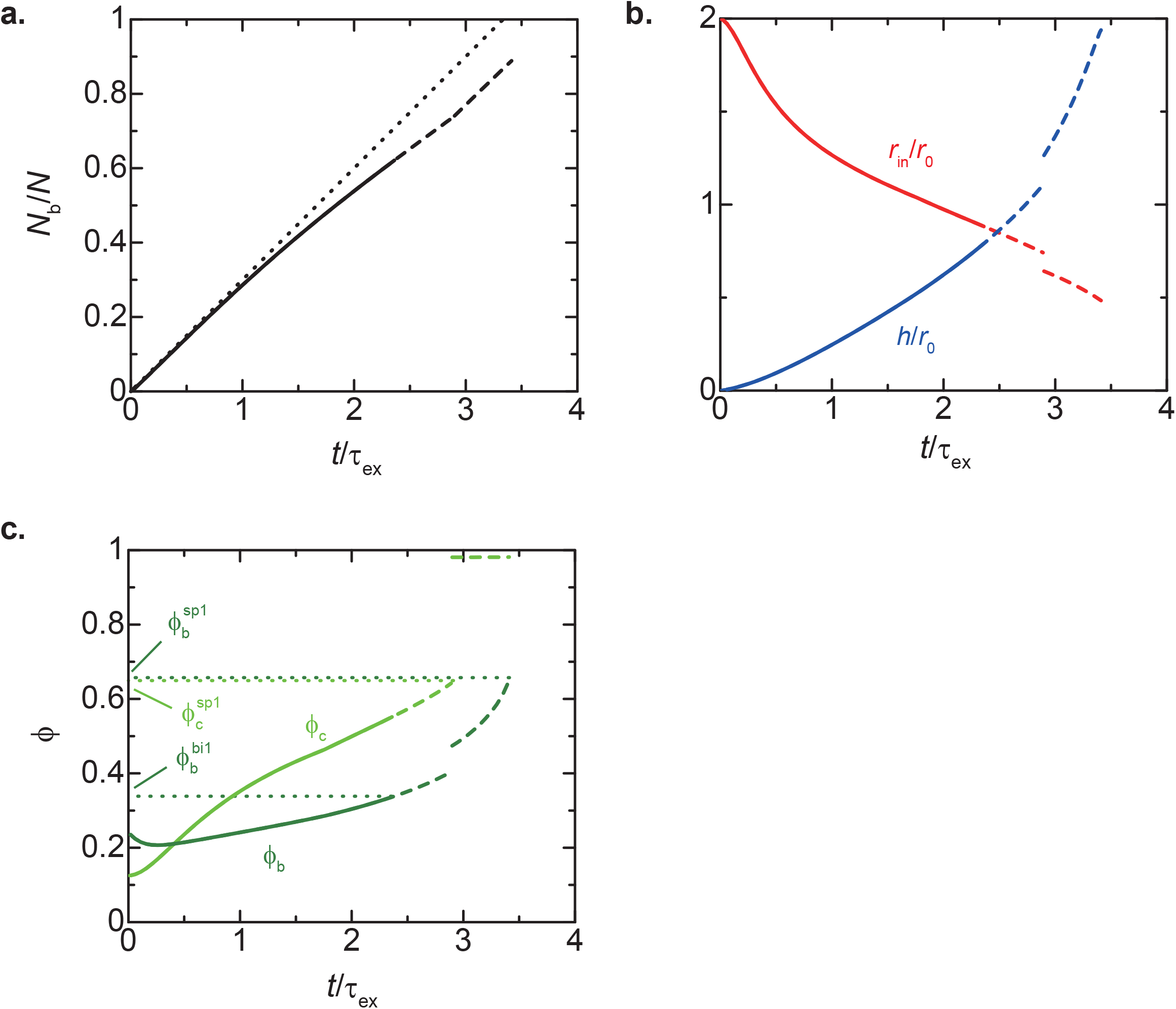
Loop extrusion dynamics with local equilibrium. **a**. Fraction of DNA units *n*_b_ in the peripheral brush region as a function of time *t*. The dotted line represents the case of ideal loop extrusion without deceleration by DNA tension and condensin unloading. **b**. The radius *r*_in_ of the central gel region (red) and the height *h* of the peripheral brush region (blue) are shown as a function of time *t*. These quantities are rescaled by the length scale *r*_0_ (Eq. (37)). **c**. DNA volume fractions in the center (light green) and periphery (deep green) are shown as a function of time 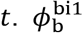and 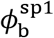 are the values of the DNA volume fraction at the binodal and spinodal lines of the periphery, respectively. The uniform swollen state is stable for 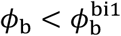 (solid lines). The two-phase coexistent state is stable, while the uniform swollen state is quasi-stable, for 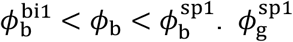 is the DNA volume fraction at the spinodal line of the central gel. Values of parameters used for these calculations are summarized in Table 1.

DNA loops produced in the early stages are enriched in condensin and are relatively swollen (magenta line, Fig. 4**a**). The probability of the binding of linker histones to the DNA loops increases with the DNA volume fraction in the peripheral region. When the DNA volume fraction in the peripheral region becomes sufficiently large, DNA loops are enriched with linker histone and condense owing to multivalent interactions. Osmotic pressure and mutual exclusion between condensin and linker histone act to suppress the loading of condensin onto condensed DNA loops rich in linker histone. Our theory predicts that the peripheral region undergoes phase separation into two domains: swollen DNA loops mainly occupied by condensin and condensed DNA loops mainly occupied by linker histone at a DNA volume fraction larger than the binodal value 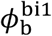. This phase separation is so-called multivalent bridging-induced phase separation (42). Because the uniform swollen state of the peripheral region is still metastable for 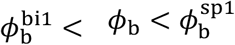, the phase separation probably does not occur immediately at 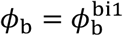, but after a lag. During this lag, loop extrusion continues to increase the DNA volume fraction *ϕ*_b_. The DNA volume fraction *ϕ*_b_ at which phase separation occurs is determined by the nucleation kinetics of domains of condensed DNA loops.

**Figure 4.**
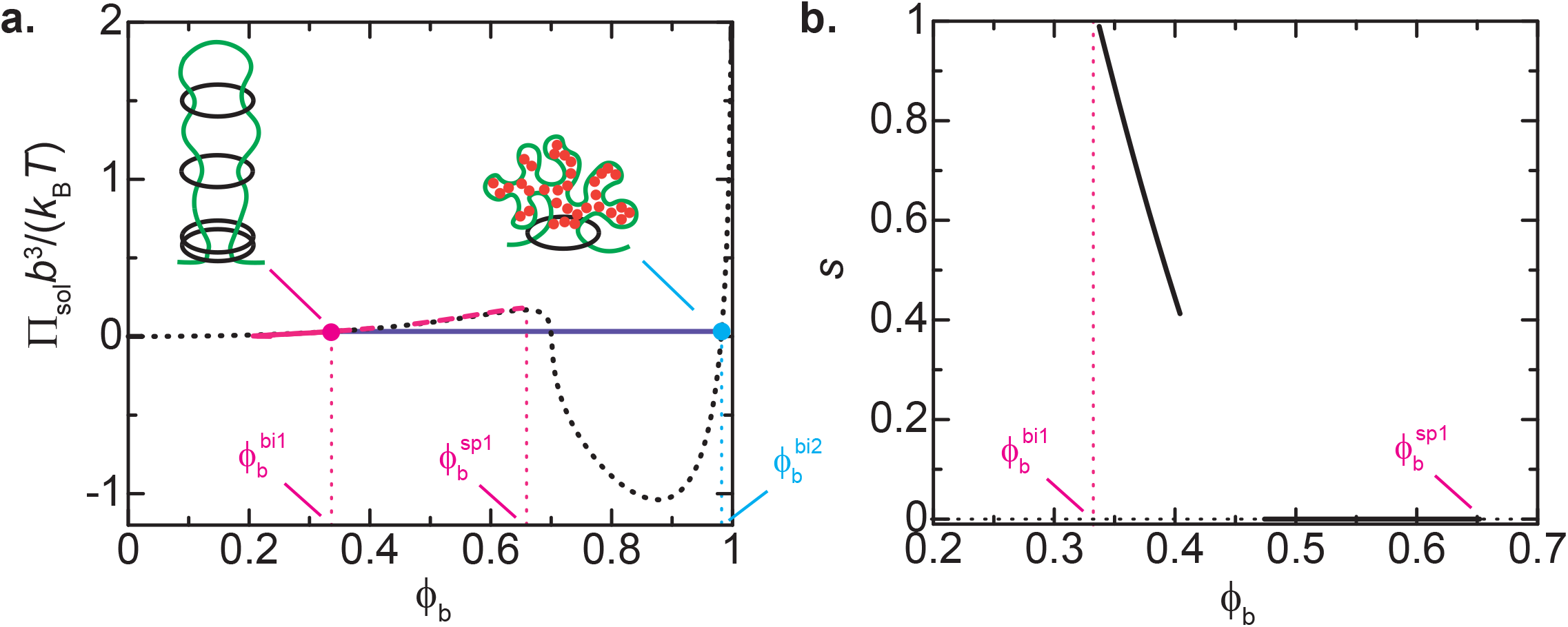
Peripheral brush region phase separation. **a**. Lateral osmotic pressure Π_sol_(*ϕ*_b_) in the brush region as a function of the DNA volume fraction *ϕ*_b_ in the periphery. Magenta line, osmotic pressure during loop extrusion (Fig. 3); dotted line, osmotic pressure with fixed DNA fraction, *n*_b_ = 0.62 and *t*/τ_ex_ = 2.36, at 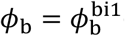(Fig. 3**c**); purple, binodal line. 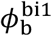 and 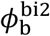 are the DNA volume fractions at the binodal line. **b**. Fraction *s* of the domains of swollen loops rich in condensin complexes as a function of DNA volume fraction *ϕ*_b_ at which phase separation occurs. Values of parameters used for these calculations are summarized in Table 1.

The fraction *s* of swollen domains depends on the DNA volume fraction *ϕ*_b_ in the peripheral region at which phase separation occurs (Fig. 4**b**). The fraction *s* is determined by the condition

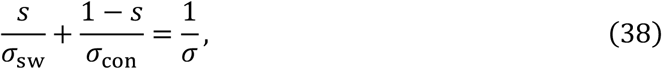

where 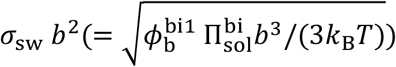 and 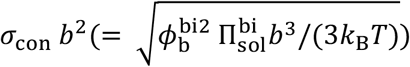 are the area densities of swollen and condensed DNA loops, respectively, with 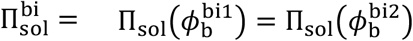 . Eq. (38) is applicable to cases in which the free energy cost arising from local shear deformation of the interface due to the inequality of area per loop, *σ*_sw_ ≠ *σ*_con_, is negligible.

### Condensed loops enriched in H1.8 are reeled into the central gel region, triggering symmetry breaking

We analyze the dynamics of DNA loops after phase separation. Steady loading of condensin is necessary to form a stable DNA loop. Swollen DNA loops, mainly occupied by condensin, are thus stable (magenta line, Fig. 5**a**). In contrast, the number of DNA units in the condensed DNA loops mainly occupied by linker histone rapidly decrease with time because loading of condensin is suppressed by the linker histone (cyan line, Fig. 5**b**). This results in breaking the spherical symmetry of the system. Therefore, our theory predicts that symmetry breaking during the assembly of the sparkler results from the phase separation in the peripheral region.

**Figure 5.**
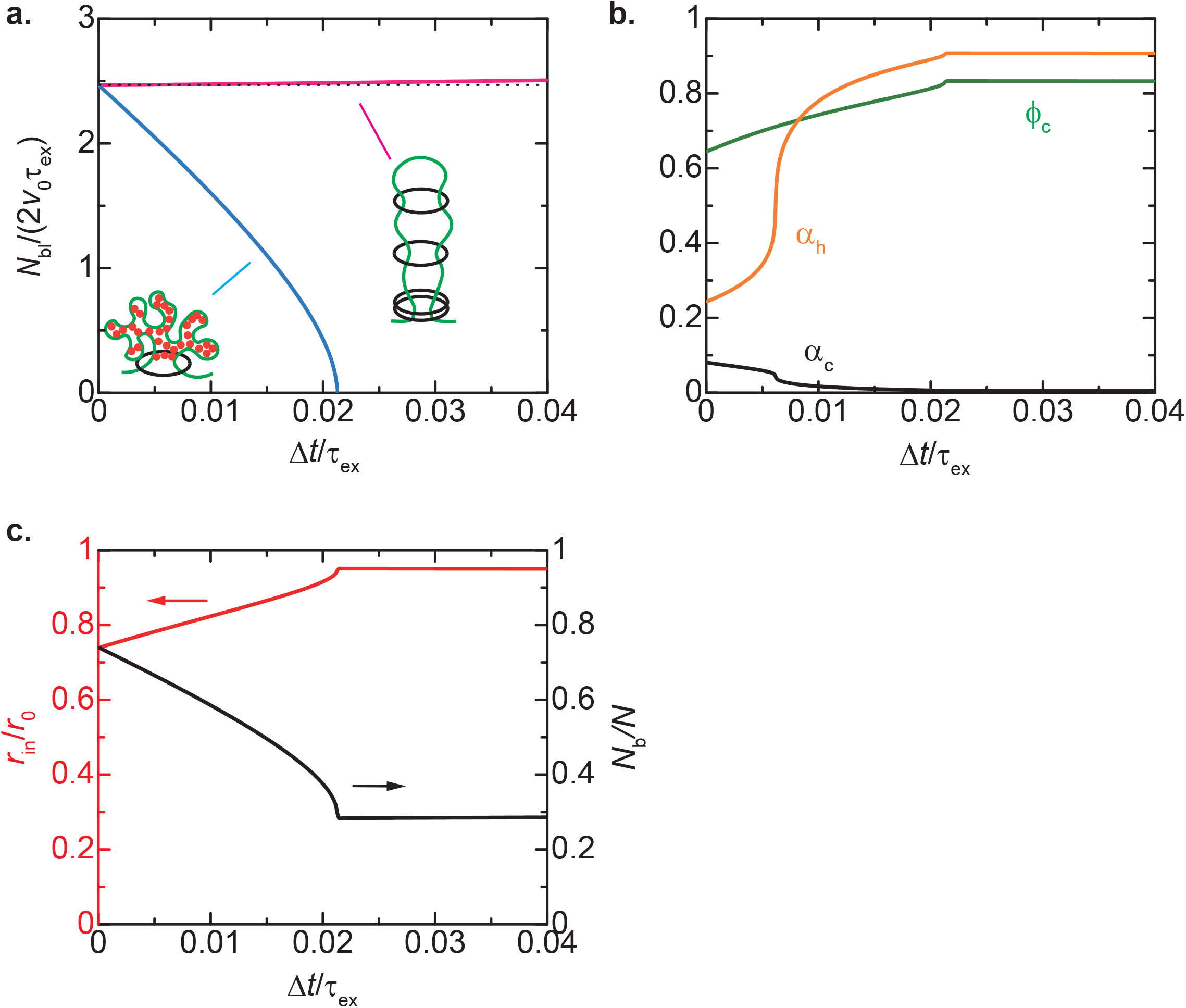
Time evolution after peripheral brush region phase separation. **a**. Numbers of units in swollen loops rich in condensin complexes (magenta) and condensed loops rich in linker histones (cyan) as a function of time Δ*t* after phase separation. **b**. DNA volume fraction *ϕ*_g_ (green), condensin occupancy *α*_c_ (black), and linker histone occupancy *α*_h_ (orange) in the central gel region as a function of time Δ*t*. **c**. Radius *r*_in_ of the central gel and fraction *n*_b_ of DNA units in the peripheral brush as a function of time Δ*t*. These are cases in which the phase separation of the peripheral brush region occurs at *n*_h_=0.74 and *t*/τ_ex_ = 2.9, corresponding to *s* = 0.4. Values of parameters used for these calculations are summarized in Table 1.

We address the dynamics of the system after phase separation using the affine approximation, in which 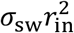 and 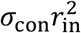 are constant, and by neglecting the solvent exchange between the central gel region and the external solution. These approximations are effective for cases in which the decrease in DNA units in the condensed loops is faster than the solvent flow from the central region to the external solution (Fig. 5). With these approximations, the DNA volume fraction *ϕ*_g_ in the central gel region increases over time because the DNA in the condensed loops at the peripheral region are reeled into the central region, (green line, Fig. 5**b**), in turn attracting linker histone to the central gel region (orange line, Fig. 5**b**). The occupancy *α*_h_ by linker histone increases continuously when the binding energy *ϵ*_h_ − *μ*_h_ is smaller than a critical value, 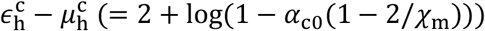. The linker histone occupancy *α*_h_ can jump at a value of DNA volume fraction *ϕ*_g_ for 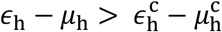,(Fig. S3 in SI). The radius *r*_in_ of the central gel region increases with time because the number of DNA units in the central gel region increase. Linker histone bound to DNA suppress the production of new loops due to mutual exclusion between linker histone and condensin.

The features predicted by our theory are consistent with those of experiments (10).

### The fast relaxation approximation can be effective when both the central and peripheral regions are in a swollen state

When loop extrusion is faster than relaxation, condensin extrudes DNA units to the peripheral region with fixed volume of the central region in the short time scales (Fig. S4 in SI). Because DNA tension is not relieved in these time scales, it greatly decelerates loop extrusion. The DNA volume fraction in the peripheral region is approximately constant after initial swelling because the volume of the central region is constant (green line, Fig. S4**d** in SI). Relaxation occurs only after loop extrusion is stopped by DNA tension, 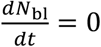 (Fig. S5 in SI). Elastic stress arising from DNA tension is balanced by the hydrostatic pressure *p*_in_ of the solvent in the central gel region, which creates the gradient of hydrostatic pressure across the peripheral region. This gradient drives solvent flow from the central region towards the external solution to relax the DNA tension. Thus, the radius *r*_in_ of the central gel region decreases over time (red line, Fig. 6**a**). The DNA volume fraction *ϕ*_b_ in the peripheral region increases during relaxation and the occupancy by linker histone increases over time (Fig. 6**b**). Our theory predicts that the peripheral brush region undergoes phase separation into swollen DNA loops mainly occupied by condensin and condensed DNA loops mainly occupied by linker histone at DNA volume fractions in the window 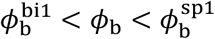, similarly to the case of fast relaxation (Fig. 6**b**). DNA in condensed loops is rapidly reeled into the central gel region. Therefore, our theory predicts symmetry breaking also for the case of slow relaxation.

**Figure 6.**
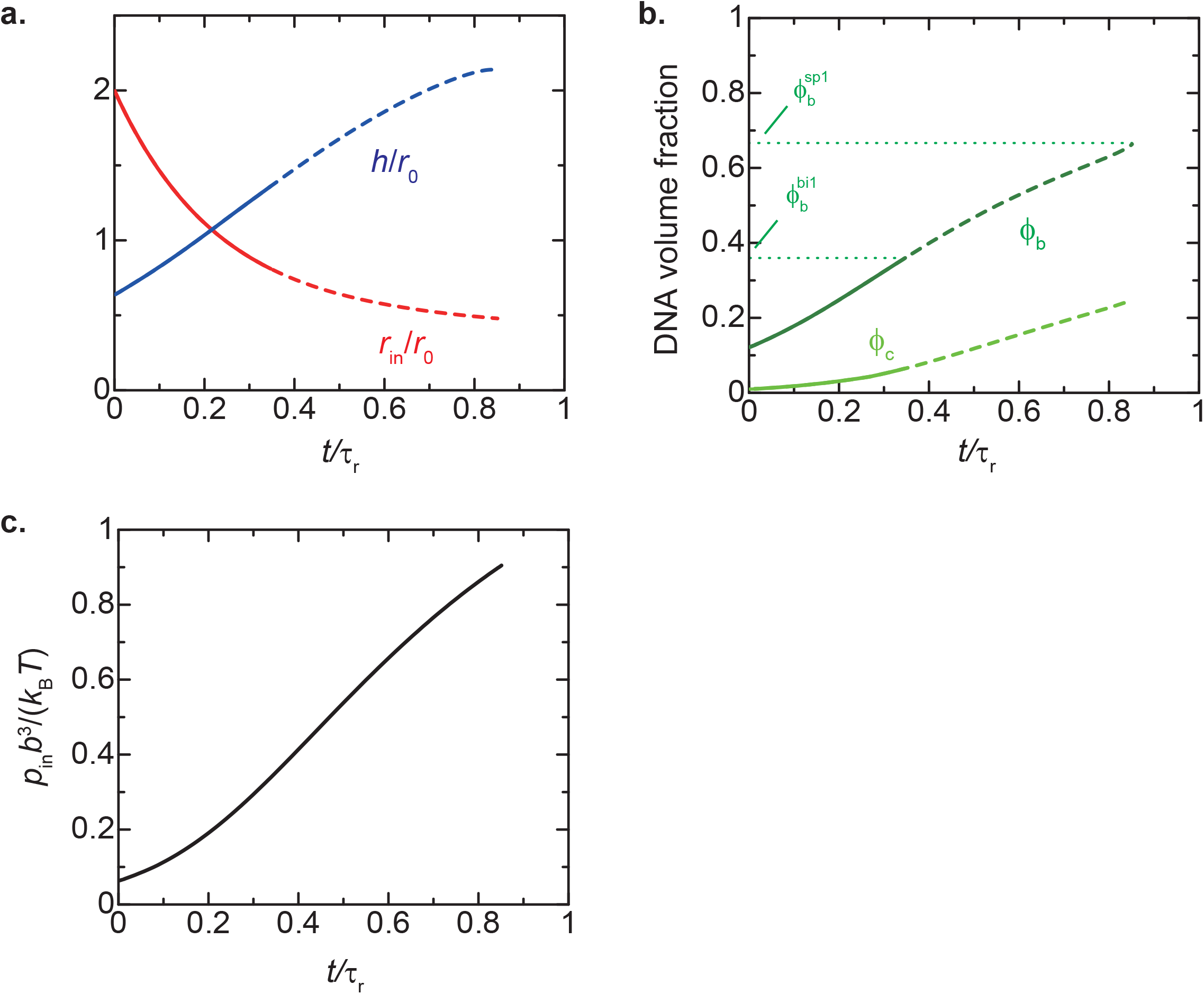
Dynamics limited by relaxation. **a**. Radius *r*_in_ of the central gel region (red) and the thickness *h* of the peripheral brush region (blue) as a function of time *t*, rescaled by the length scale *r*_0_, (Eq. (37)). **b**. DNA volume fractions in the center (light green) and periphery (deep green) as a function of time *t*. **c**. Hydrostatic pressure *p*_in_ as a function of time *t*, rescaled by the relaxation time *τ*_r_, (Eq. (39)). 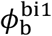 Is the DNA volume fraction at the binodal line. The most stable states are the uniform brush of swollen DNA loops for 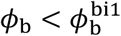 (solid line) and the coexistence of the swollen and condensed DNA loops for 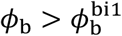 (broken line). Values of parameters used for these calculations are summarized in Table 1.

In contrast to many passive systems, the hydrostatic pressure *p*_in_ increases with time during relaxation because loop extrusion by condensin reinforces DNA tension (Figs. 6**c** and S5**a**). Nevertheless, solvent flow decreases over time because the thickness *h* of the peripheral region increases (blue line, Fig. 6**a**).

Our theory predicts that the relaxation time has the form

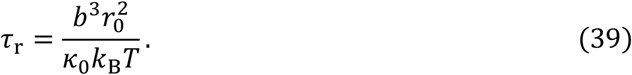

The Darcy constant *κ*_0_ (≈ *b*^2^/*η*) is estimated as 5 x 10^-13^ [m^2^/Pa s] by using the viscosity of Xenopus egg extracts, *η* ≈ 20 mPa s (43). The relaxation time *τ*_r_ ∼ 1.5 min is thus ∼ 10-fold shorter than that of the loop extrusion *τ*_ex_ ≈ 10 min; the relaxation is rather rapid. The relaxation time *τ*_ex_ can be longer than the time scale, with which the DNA in condensed loops is reeled to the central region after the phase separation (cyan line, Fig. 5**a**), consistent with our assumption in the previous section.

Relaxation time increases with increasing thickness and DNA volume fraction in the peripheral brush region. With fast relaxation, these quantities increase with time (blue line, Fig. 3**b** and dark green line, Fig. 3**c**). One may think that the relaxation per unit step of loop extrusion can eventually become slower than that of the time scale of the unit extrusion step at longer time scales. By using eq. (30), the relaxation time Δ*τ*_r_ per unit step of loop extrusion is derived as

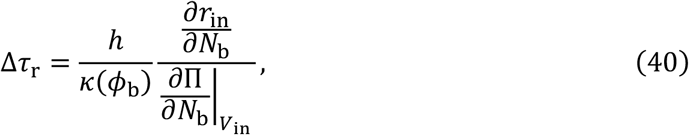

Where 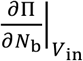 represents the partial derivative of Π with respect to *N*_b_ with the constraint of constant volume *V*_in_. The derivation of eq. (40) is shown in Sec. S3 in SI. Eq. (40) suggests that the relaxation time Δ*τ*_r_ per unit step can be much longer than the step time 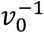 of loop extrusion when all DNA loops in the peripheral region are condensed and *ϕ*_b_ ≈ 1 . Solvent flow within the central gel region can also be prohibitively slow when the central gel region is condensed, *ϕ*_g_ ≈ 1. Thus, the volume *V*_in_ of the central gel region is approximately constant if either or both the central and peripheral regions are condensed. Indeed, it is well known that the volume-phase transition of a polymer gel is a very slow process for the same reason (27,44-46). With the parameter values shown in Table 1, the relaxation time Δ*τ*_r_ (< 0.01 *τ*_r_) per unit step is estimated as ∼ 10-fold less than the time between the steps of loop extrusion, 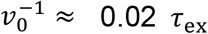 before the phase separation (Fig. 7). However, for cases in which the shear modulus *G*_0_*b*^3^/(*k*_B_*T*) is small, the relaxation time Δ*τ*_r_ per unit extrusion step can be larger than the time 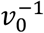 per unit extrusion step at longer time scales (Sec. S5.10 in SI).

**Figure 7.**
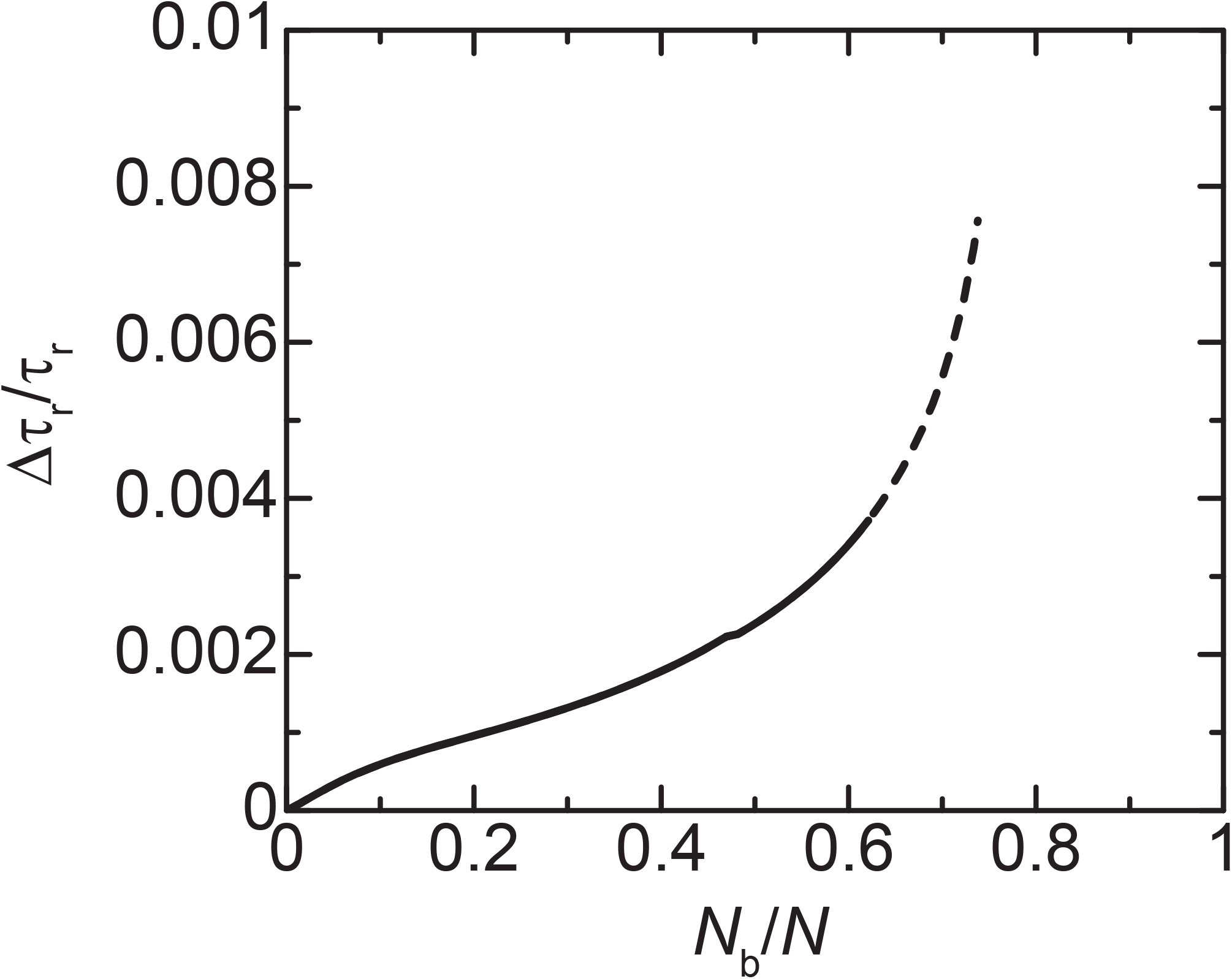
Relaxation time per unit step of loop extrusion (Δ*τ*_r_)*τ*_r_ as a function of the fraction of DNA units in the peripheral brush region, rescaled by the relaxation time *τ*_r_, (Eq. (39)). Values of parameters used for these calculations are summarized in Table 1.

## DISCUSSION

The sparkler is a highly characteristic chromatin structure assembled on nucleosome-free, entangled DNA in *Xenopus* egg extracts. In the current study, we have developed a theory that predicts the mechanism of symmetry breaking observed during sparkler assembly. According to this theory, the size of the central gel region decreases to relieve the DNA tension produced by condensin I-mediated loop extrusion occurring at the central-peripheral interface (Fig. 8**a**). This leads to a higher DNA volume fraction in the peripheral region, thereby increasing the likelihood of linker histone H1.8 binding to this region. When the DNA volume fraction in the peripheral region becomes sufficiently high, competition between condensin I and H1.8 drives phase separation of the peripheral region into two domains: swollen loops enriched in condensin I and condensed loops enriched in H1.8 (Fig. 8**b**). Experimentally, protrusions with condensin I enriched at their tips are irregularly positioned in the final structure of a sparkler (10), suggesting a hallmark of phase separation. DNA loops enriched in condensin I are relatively stable, whereas those enriched in H1.8 are rapidly reeled into the central gel region, as H1.8 competes with condensin I and suppresses its loading onto these loops, thereby breaking the spherical symmetry of the system (Fig. 8**b**). Thus, our theory predicts that H1.8 regulates the stability of DNA loops extruded by condensin I and promotes their reorganization. Other variants of linker histones likely exhibit similar functions, provided that they compete with condensin I and that their multivalent interactions are sufficiently strong. H1.8 also suppresses the formation of new DNA loops in the central gel region following reorganization. In summary, the series of events observed during sparkler formation is driven by competition between condensin I and H1.8, as well as DNA tension generated through loop extrusion. Importantly, these two key factors exert their greatest effects on the entangled, nucleosome-free DNA substrates used in the experiments (10).

**Figure 8.**
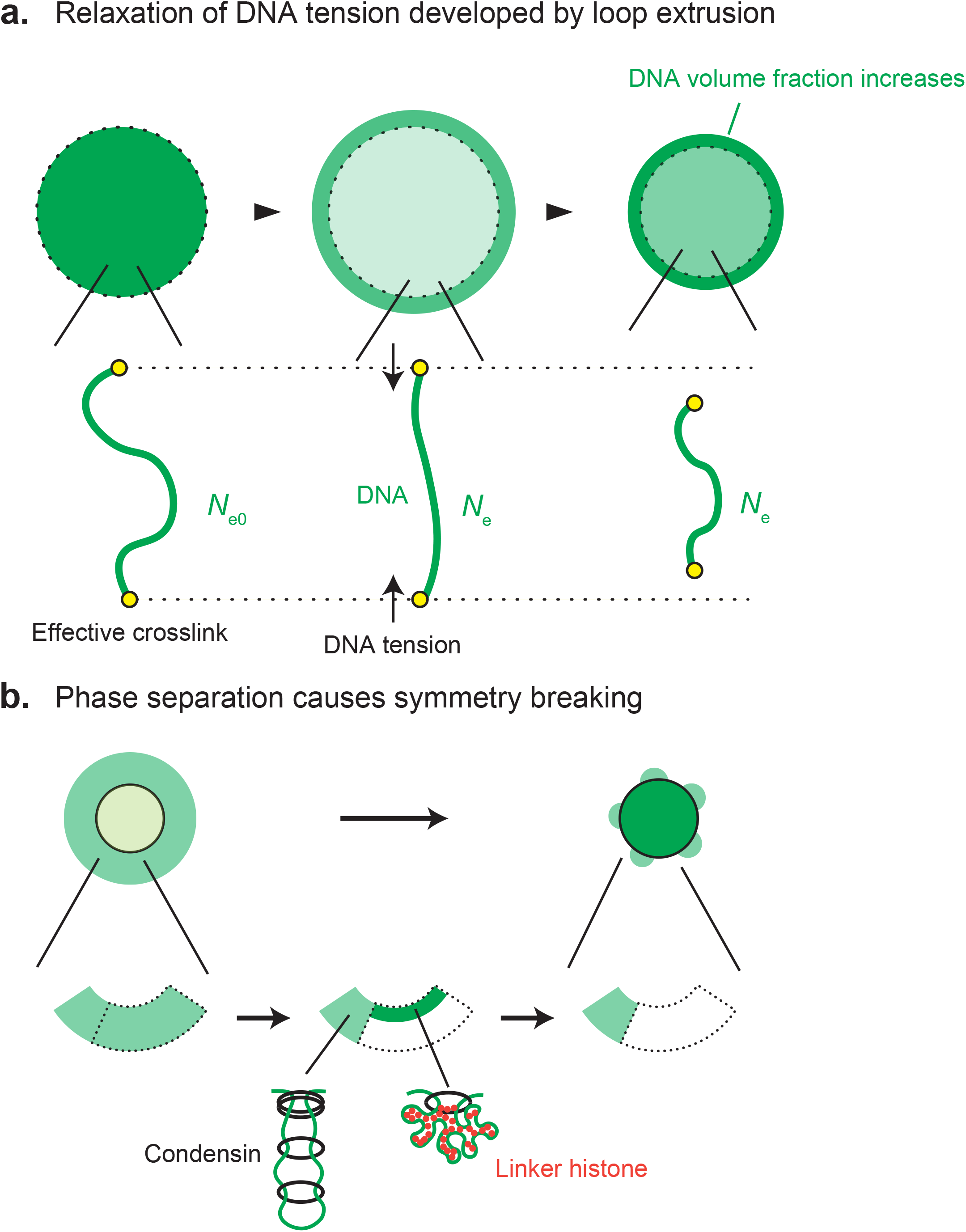
A proposed mechanism of symmetry breaking during sparkler formation. **a**.DNA tension is generated in the central region by condensin I-mediated loop extrusion occurring in the peripheral region (middle). Solvent flow from the center to the surrounding solution reduces the central volume, thereby alleviating DNA tension (right). This in turn increases the DNA volume fraction in the peripheral region. **b**. As the DNA volume fraction in the peripheral region increases, H1.8 is attracted to this region. The peripheral region then undergoes phase separation into two domains: swollen loops rich in condensin I and condensed loops rich in the linker histone H1.8 (middle). The DNA in the condensed loops is rapidly reeled into the central region (right).

Following symmetry breaking, what mechanism drives the subsequent morphological changes into the unique sparkler structure containing protrusions? One possibility is that attractive multivalent interactions between H1.8 and DNA generate surface tension at the interface between the central gel region and the surrounding solution, whereas the swollen loop domains are not subject to such surface tension. Although surface tension acting on a uniform, closed surface stabilizes spherical shapes to minimize surface area, surface tension acting on a non-uniform surface can produce more complex shapes. One of such examples can be observed in soap films suspended in frames, which form catenoids (47). Multicomponent lipid membranes also form heterogeneous surfaces and exhibit budding, although the primary driving force in this case is heterogeneity in bending rigidity (48,49). Heterogeneous surface tension arising from phase separation likely plays a key role in the appearance of protrusions. The size of the swollen loop domains is likely determined by a combination of factors: condensin I-mediated loop extrusion at the interface, loop branching resulting from the simultaneous loading of multiple condensin I complexes, and mechanical tension at the three-phase contact line, where the swollen brush phase, condensed gel phase, and surrounding solution converge. Our current theory lays a foundation for future studies aimed at better understanding how protrusions form.

The current study represents the first attempt to provide a theoretical framework for the formation of a recently described peculiar chromatin structure. As discussed below, however, we have made several assumptions to simplify our current model of symmetry breaking. First, we neglected the effects of the curvature of the central-peripheral interface on the DNA volume fraction distribution in the peripheral region, as these effects were well characterized (31). Second, we did not incorporate the relaxation dynamics of DNA arising from loop extrusion, although one can also take it into account using recent theoretical frameworks (50-52). Third, we assumed a linear relationship between the loop extrusion rate and DNA tension, although one could also use a fitted rate-tension curve derived from experimental data (8) or extend the two-state model of molecular motors (53). Fourth, we assumed that condensin I loaded onto the peripheral brush region does not penetrate the central gel region. This assumption was based on the observation that, over relatively short time scales during which the entangled DNA strands remain spherical, condensin I is most concentrated at the interface between the central and peripheral regions (10). We speculate that this is because DNA loops associated with condensin I are repelled by the osmotic pressure exerted by the central region. Fifth, we used free-energy minimization, which is valid at the thermodynamic equilibrium, to predict phase separation in the peripheral region. This leads to equalities in the chemical potential and osmotic pressure between the two coexisting domains, as described in Eqs. (33) and (34), respectively. Sixth, we neglected the branching of DNA loops in the peripheral brush region because it is a rare event that becomes significant only over longer timescales (5). Finally, we assumed that DNA loops in the peripheral region consist of an equal number of DNA units. Although incorporating these factors would make the theory more quantitative, we expect that the underlying physics would remain unchanged.

Although the number of parameters involved in our model may appear large, most of them are, in principle, experimentally measurable. We derive the asymptotic analytical expressions in Sec. S5 in SI and discuss the dependence of the concentration *ϕ*_c0_ of condensin complexes and the shear modulus *G*_0_*b*^3^/(*ϕ*_0_*k*_B_*T*) in Sec. S6 in SI. Briefly, when the condensin occupancy *α*_c0_ is too small, the loops in the peripheral region are unstable or reach the steady state before phase separation occurs, see Fig. S7. The ratio *ϕ*_g_/*ϕ*_b_ of the DNA volume fractions decreases with decreasing the shear modulus 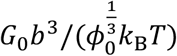, see Fig. S8. Experimentally, the DNA volume fraction *ϕ*_g_ in the central region is smaller than that in the peripheral region *ϕ*_b_, implying that the shear modulus of the entangled DNA strands is rather small. The asymptotic analytical expressions in Sec. S5 capture these results of numerical calculations.

It would be worthwhile to compare our current model of sparkler formation with our previous model of bean formation (32). The bean, a compact, round-shaped structure, is formed in *Xenopus* egg extracts in which topo IIα is depleted and endogenous condensins are replaced with a condensin I mutant defective in loop extrusion (11). The bean consists of a central core, where DNA and the mutant condensin I are concentrated, and a peripheral halo, where DNA is sparsely distributed. Notably, core histones are enriched in the peripheral halo, suggesting a competition between condensin I and nucleosomes. Like beans, the spherical precursors of sparklers prior to symmetry breaking also exhibit bipartite organization, although the spatial distributions of histones and condensin I are inverted between the two structures. A key difference between the two structures lies in their underlying assembly mechanisms: the halo of beans is formed through the free energy of nucleosome assembly (32), whereas the precursor of sparklers is assembled by condensin I-mediated active loop extrusion (this study). Despite this difference, competition between histones and condensin I likely serves as a unifying principle in both cases.

One could argue that our present theory on sparkler assembly and our previous theory on bean formation (32) are not directly relevant to our understanding of mitotic chromosome assembly because the two structures are assembled only under specialized conditions in *Xenopus* egg extracts. In thermodynamics, a phase diagram defines the conditions under which different states of a system emerge and serves as a powerful tool for understanding the properties of systems composed of many molecules. From a broader perspective, mitotic chromosomes, along with structures such as sparklers and beans, can be viewed as parts of a phase diagram of chromatin organization defined by the activities of essential factors. Further exploration of such a phase diagram is likely to elucidate how condensins assemble mitotic chromosomes through their interactions with other essential factors in a crowded environment.

## Supporting information

Supplementary Figures and Discussion

## Acknowledgement

This work was supported by JSPS KAKENHI Grant Numbers JP20H05934 (T. Y.), JP24K06969 (T. Y.), JP23K23815 (K. S.), JP20H05938 (T. H.). T. Y. thanks John Marko (Northwestern Univ.) and Anton Goloborodko (IMBA) for fruitful discussion.

## Data availability

The Mathematica file ‘‘antennamodelVer155_doublelayer(brushcorrection).nb’’ used for the numerical calculations is available in figshare with the identifier (10.6084/m9.figshare.28900841).

