## Supplementary Figures and Discussion for "Reorganization of DNA loops by competition between condensin I and a linker histone"

### Supplementary Information: Structures of entangled DNA strands assembled by condensin complexes and linker histones

### S1 Scaling theory of polymer brush

#### S1.1 Planer brush

We here review the scaling theory of a polymer brush, following the pioneering work by de Gennes.[1] We here treat polymers end-grafted to a planer surface with the grafting density $\sigma$. Each polymer is composed of $N$ (repeat) units of Kuhn length $b$. The units in a polymer are correlated because of their connectivity. However, when the polymer concentration is large enough, the correlation is effective only at the length scales smaller than the correlation length $\xi$ and is screened at the larger length scales because of the interaction between polymers. The essence of de Gennes' theory is that the correlation length of polymers in a brush is given by the average distance $\sigma^{-1/3}$ between grafting point, $\xi = \sigma^{-1/2}$. Because of this property, a polymer in a brush is considered as a string of blobs, where the size of each blob is $\xi$, and the blobs are filled the space $0 < z < h$ above the grafting point, where $h$ is the brush height. The number $g$ of monomers in each blob is estimated by the relationship,

$$\xi = bg^{\nu}, \tag{S1}$$

where $\nu$ is the inverse of the fractal dimension of a polymer with $\nu = 1/2$ in a $\theta$-solvent and $\nu = 3/5$ in an athermal solvent. Because the blobs are space filling, the brush height $h$ has the form

$$h = \xi \frac{N}{g} = bN(\sigma b^2)^{(1-\nu)/(2\nu)} \tag{S2}$$

because the number of blobs is $N/g$ per polymer and each blob has the size $\xi$. The osmotic pressure in a brush has the form

$$\Pi_{\rm in} = \frac{k_{\rm B}T}{\xi^3} = k_{\rm B}T\sigma^{3/2}. \tag{S3}$$

#### S1.2 Spherical brush

A polymer brush prepared on a spherical grafting surface of radius $r_{\rm in}$ is treated by using the Daoud-Cotton theory.[2] As the previous section, we treat a polymer brush composed of $n_{\rm p}$ polymers, each composed of $N$ units of length $b$. This theory takes into account the fact that the correlation length $\xi(r)$ increases with the distance $r$ from the center of the spherical surface. The fact that the spherical area at the distance $r$ from the center is filled with the blobs of $n_{\rm p}$ polymers leads to the relationship

$$4\pi r^2 = n_{\rm p}\xi^2(r). \tag{S4}$$

Because the blobs fill the space between $r_{\rm in} < r < r_{\rm ex}$, the conservation of the number of units has the form

$$n_{\rm p}N = \int_{r_{\rm in}}^{r_{\rm ex}} \frac{4\pi r^2 dr}{\xi^3(r)} g(r), \tag{S5}$$

where $g(r)$ is the number of units in a blob of size $\xi(r)$ and is derived by using eqs. (S1) and (S4). By using eq. (S5), the external radius $r_{\rm ex}$ is derived as

$$r_{\rm ex} = r_{\rm in}\left(1 + \frac{1}{\nu}\frac{h}{r_{\rm in}}\right)^{\nu} \tag{S6}$$

with

$$h = bN\left(\frac{n_{\rm p}b^2}{4\pi r_{\rm in}^2}\right)^{(1-\nu)/(2\nu)}. \tag{S7}$$

Eq. (S7) returns to eq. (S2) by using the grafting density,

$$\sigma = n_{\rm p}/(4\pi r_{\rm in}^2). \tag{S8}$$

The free energy of the polymer brush has the form

$$\begin{aligned} F &= k_{\mathrm{B}}T \int_{r_{\mathrm{in}}}^{r_{\mathrm{ex}}} \frac{4\pi r^2 dr}{\xi^3(r)} \\ &= k_{\mathrm{B}}T n_{\mathrm{p}} \left(\frac{n_{\mathrm{p}}}{4\pi}\right)^{1/2} \nu \log\left(1 + \frac{1}{\nu}\frac{h}{r_{\mathrm{in}}}\right) \end{aligned} \quad \text{(S9)}$$

because the free energy of each blob is estimated as $k_{\mathrm{B}}T$. The osmotic pressure $\Delta\Pi$ applied to the spherical surface has the form

$$\begin{aligned} \Delta\Pi &= -\frac{1}{4\pi r_{\mathrm{in}}^2}\frac{\partial F}{\partial r_{\mathrm{in}}} \\ &= \Pi_{\mathrm{in}} \frac{\frac{1}{\nu}\frac{h}{r_{\mathrm{in}}}}{1 + \frac{1}{\nu}\frac{h}{r_{\mathrm{in}}}}. \end{aligned} \quad \text{(S10)}$$

with

$$\Pi_{\mathrm{in}} = \left(\frac{n_{\mathrm{p}}}{4\pi r_{\mathrm{in}}^2}\right)^{3/2}. \quad \text{(S11)}$$

Eq. (S11) returns to eq. (S3) by using eq. (S8).

For $h < r_{\mathrm{in}}$, eq. (S10) has an asymptotic form

$$\Delta\Pi_{\mathrm{in}} \approx \Pi_{\mathrm{in}} \frac{h}{r_{\mathrm{in}}}. \quad \text{(S12)}$$

For $h > r_{\mathrm{in}}$, eq. (S10) has an asymptotic form

$$\Delta\Pi_{\mathrm{in}} \approx \Pi_{\mathrm{in}}. \quad \text{(S13)}$$

### S2 Osmotic pressure, chemical potential, and DNA tension

#### S2.1 Free energy

We here treat entangled DNA strands composed of $N$ DNA units of length $b$ in a solution of condensin complexes and linker histones. The loop extrusion activity of condensin complexes assembles a brush of DNA loops at the periphery of the central region, which forms an entangled DNA gel. It also produces elastically ineffective loops in the central gel region. The number of DNA units in the system is conserved,

$$N_{\mathrm{eff}} + N_{\mathrm{gl}} + N_{\mathrm{b}} = N, \tag{S14}$$

where $N_{\mathrm{eff}}$ and $N_{\mathrm{gl}}$ are the numbers of DNA units in the elastically effective and ineffective parts of the central gel region, respectively, and $N_{\mathrm{b}}$ is the number of DNA unit in the peripheral brush region. The partition of DNA units into these regions is determined by the loop extrusion dynamics. The structure of the entangled DNA strands is represtend by the DNA volume fraction $\phi$, the occupancy $\alpha_{\mathrm{c}}$ of condensin complexes, the occupancy $\alpha_{\mathrm{h}}$ of linker histones. The DNA volume fraction $\phi_{\mathrm{g}}$ at the central gel region has the form

$$\phi_{\mathrm{g}} = \frac{b^3(N - N_{\mathrm{b}})}{V_{\mathrm{in}}} = \frac{(1 - n_{\mathrm{b}})\phi_0}{\lambda_{\mathrm{s}}^3}, \tag{S15}$$

where $V_{\mathrm{in}}$ $(= 4\pi r_{\mathrm{in}}^3/3)$ is the volume of the central gel region and $r_{\mathrm{in}}$ is the radius of the central gel region. $\lambda_{\mathrm{s}}$ is the swelling ratio. $n_{\mathrm{b}}(= N_{\mathrm{b}}/N)$ is the fraction of DNA units in the peripheral region. The DNA volume fraction $\phi_{\mathrm{b}}$ in the peripheral brush region has the form

$$\phi_{\mathrm{b}} = \frac{b^3 N_{\mathrm{b}}}{4\pi r_{\mathrm{in}}^2 h}, \tag{S16}$$

where $h$ is the thickness of the peripheral brush region.

The free energy of the system thus has the form

$$F = F_{\mathrm{g}} + F_{\mathrm{b}}, \tag{S17}$$

where $F_{\mathrm{g}}$ is the free energy of the central gel region and $F_{\mathrm{b}}$ is the peripheral brush region. The free energy $F_{\mathrm{g}}$ of the entangled DNA gel has the form

$$F_{\mathrm{g}} = \int_0^{r_{\mathrm{in}}} 4\pi r^2 dr\, (f_{\mathrm{ent}}(\phi) + f_{\mathrm{sol}}(\alpha_{\mathrm{c}}, \alpha_{\mathrm{h}}, \phi) + f_{\mathrm{bnd}}(\alpha_{\mathrm{c}}, \alpha_{\mathrm{h}}, \phi) + f_{\mathrm{c}}(\phi_{\mathrm{c}})), \tag{S18}$$

where $f_{\mathrm{ent}}(\phi)$ is the elastic free energy per unit volume due to the entanglement, $f_{\mathrm{sol}}(\alpha_{\mathrm{c}}, \alpha_{\mathrm{h}}, \phi)$ is the solution free energy per unit volume, $f_{\mathrm{bnd}}(\alpha_{\mathrm{c}}, \alpha_{\mathrm{h}}, \phi)$ is the binding free energy per unit volume, and $f_{\mathrm{c}}(\phi_{\mathrm{c}})$ is the free energy of condensin complexes that are not bound to DNA ($\phi_{\mathrm{c}}$ is the volume fraction of such condensin complexes). The free energy $F_{\mathrm{pln}}$ of the planar brush region has the form

$$F_{\mathrm{b}} = \int_{r_{\mathrm{in}}}^{r_{\mathrm{ex}}} 4\pi r^2 dr\, (f_{\mathrm{lop}}(\phi) + f_{\mathrm{sol}}(\alpha_{\mathrm{c}}, \alpha_{\mathrm{h}}, \phi) + f_{\mathrm{bnd}}(\alpha_{\mathrm{c}}, \alpha_{\mathrm{h}}, \phi) + f_{\mathrm{c}}(\phi_{\mathrm{c}})), \tag{S19}$$

where $f_{\mathrm{lop}}(\phi)$ is the elastic free energy per unit volume of DNA loops in the peripheral brush region.

The elastic free energy due to the entanglement has the form

$$f_{\mathrm{ent}}(\phi) = \frac{3}{2} G_0 \frac{\phi_{\mathrm{eff}}}{\phi_0} \left( \frac{N_{\mathrm{e0}}}{N_{\mathrm{e}}} \right)^2 \lambda_{\mathrm{s}}^2. \tag{S20}$$

Eq. (S20) represents the free energy by swelling an entangled gel from the reference state, when the DNA strands are entangled, by the swelling ratio $\lambda_{\mathrm{s}}$. $G_0$ is the shear modulus and $\phi_0$ is the DNA volume fraction, both at the reference state. The subscript 0 indicates the quantitites at the reference state. $N_{\mathrm{e}}$ is the number of DNA units between effective crosslinks due to the entanglement and $N_{\mathrm{e0}}$ is the value of $N_{\mathrm{e}}$ at the reference state. The fact that the number of effective crosslinks is constant leads to the relationship

$$\frac{N}{N_{\rm e0}} = \frac{N_{\rm eff}}{N_{\rm e}}, \tag{S21}$$

where $N_{\rm eff}$ is the number of DNA units in the elastically effective part of the central gel region. $\phi_{\rm eff}$ is the volume fraction of elastically effective chains and has the form

$$\phi_{\rm eff} = \frac{b^3 N_{\rm eff}}{V_{\rm in}} = n_{\rm e}\frac{\phi_0}{\lambda_{\rm s}}, \tag{S22}$$

where $n_{\rm e}$ $(= N_{\rm e}/N_{\rm e0})$ is the fraction of DNA units in the elastically effective part of the central gel region.

The solution free energy per unit volume has the form

$$\begin{aligned} f_{\rm sol}(\alpha_{\rm c}, \alpha_{\rm h}, \phi) \quad = \quad & \frac{k_{\rm B}T}{b^3}\left[(1-(1+\alpha_{\rm c})\phi-\phi_{\rm c})\log(1-(1+\alpha_{\rm c})\phi-\phi_{\rm c}) - \chi_0\phi^2 - \chi_{\rm m}\alpha_{\rm h}^2\phi^2 \right. \\ & \left. -\mu_{\rm h}\alpha_{\rm h}\phi - \mu_{\rm c}\alpha_{\rm c}\phi\right]. \end{aligned} \tag{S23}$$

Eq. (S23) is composed of the mixing free energy of solvent (the first term), the free energy due to the DNA-DNA interaction (the second term), the free energy due to the linker histone-linker histone interaction (the third term), and the chemical potentials of linker histones and condensin complexes (the fourth and fifth terms). $\chi_0$ is the interaction parameter that accounts for the DNA-DNA interaction. $\chi_{\rm m}$ is the interaction parameter that accounts for the linker histone-linker histone interaction. $\mu_{\rm h}$ and $\mu_{\rm c}$ are the chemical potentials of linker histones and condensin complexes. For simplicity, we assume that the volumes of a DNA unit, solvent, and a condensin complex are equal and neglected the volume of linker histones in the derivation of the mixing free energy.

The binding free energy per unit volume has the form

$$f_{\mathrm{bnd}}(\alpha_{\mathrm{c}}, \alpha_{\mathrm{h}}, \phi) = \frac{k_{\mathrm{B}}T}{b^3}\phi\left[\alpha_{\mathrm{c}}\log\alpha_{\mathrm{c}} + \alpha_{\mathrm{h}}\log\alpha_{\mathrm{h}} + (1-\alpha_{\mathrm{c}}-\alpha_{\mathrm{h}})\log(1-\alpha_{\mathrm{h}}-\alpha_{\mathrm{c}}) + \epsilon_{\mathrm{h}}\alpha_{\mathrm{h}} + \epsilon_{\mathrm{c}}\alpha_{\mathrm{c}}\right]. \tag{S24}$$

Eq. (S24) is composed of the entropic free energy due to the occupancy of linker histones and condensin complexes (the first, second, and third terms), the free energy due to the binding of linker histones and DNA units (the fourth term), the free energy due to the binding of condensin complexes and DNA units (the fifth term).

The free energy per unit volume of freely diffusing condensin has the form

$$f_{\mathrm{c}}(\phi_{\mathrm{c}}) = \frac{k_{\mathrm{B}}T}{b^3}\left[\phi_{\mathrm{c}}\log\phi_{\mathrm{c}} - \mu_{\mathrm{c}}\phi_{\mathrm{c}}\right]. \tag{S25}$$

The elastic free energy per unit volume of DNA loops in the peripheral region has the form

$$f_{\mathrm{lop}}(\phi) = \frac{3}{2}\frac{k_{\mathrm{B}}T}{b^3}\frac{(\sigma b^2)^2}{\phi}, \tag{S26}$$

where $\sigma$ $(= 2n_{\mathrm{l}}/(4\pi r_{\mathrm{in}}^2))$ is the surface density of DNA loops and depends on the swelling ratio $\lambda_{\mathrm{s}}$ of the central gel region. The surface density $\sigma$ has the relationship

$$\sigma = \sigma_0\left(\frac{\phi_{\mathrm{g}}}{1-n_{\mathrm{b}}}\right)^{2/3} \tag{S27}$$

with the DNA volume fraction $\phi_{\mathrm{g}}$ in the central gel region, where $\sigma_0$ $(= \eta_{\mathrm{m}}^{2/3}/b^2)$ is the surface density for the case of dried gel, $\phi_{\mathrm{g}}/(1-n_{\mathrm{b}}) = 1$, with

$$\eta_{\mathrm{m}} = \frac{1}{3\sqrt{4\pi\frac{(2n_{\mathrm{l}})^{3/2}}{N}}}. \tag{S28}$$

#### S2.2 Volume fraction of freely diffusing condensin

At the minimum of the free energy, the variation of the free energy with respect to $\phi_c$ is zero

$$\frac{\delta F}{\delta \phi_c} = \frac{k_B T}{b^3} [\log \phi_c + 1 - \mu_c - \log(1 - (1 + \alpha_c)\phi - \phi_c) - 1] = 0, \tag{S29}$$

where we note that the free energy depends on the volume fraction $\phi_c$ only via eqs. (S23) and (S25). This condition leads to the form

$$\mu_c = \log \phi_c - \log(1 - (1 + \alpha_c)\phi - \phi_c). \tag{S30}$$

Eq. (S30) defines the local chemical potential of condensin complexes. For cases in which the diffusion of condensin complexes is fast, the chemical potential $\mu_c$ of condensin complexes is uniform. In such cases, the volume fraction $\phi_c$ is derived as

$$\phi_c = (1 - (1 + \alpha_c)\phi)\phi_{c0} \tag{S31}$$

with

$$\phi_{c0} = \frac{e^{\mu_c}}{1 + e^{\mu_c}} \tag{S32}$$

Eq. (S31) represents the fact that the osmotic pressure arising from DNA suppresses condensin complexes from penetrating the peripheral and central regions.

Substituting eq. (S31) into the first term of eq. (S23) leads to the form

$$\begin{aligned} & (1 - (1 + \alpha_c)\phi - \phi_c) \log(1 - (1 + \alpha_c)\phi - \phi_c) \\ = \; & (1 - (1 + \alpha_c)\phi)(1 - \phi_{c0})(\log(1 - (1 + \alpha_c)\phi) + \log(1 - \phi_{c0})) \\ \simeq \; & (1 - (1 + \alpha_c)\phi) \log(1 - (1 + \alpha_c)\phi), \end{aligned} \tag{S33}$$

where we used $\phi_{c0} \ll 1$ to derive the last form. We thus use

$$\begin{aligned} f_{\mathrm{sol}}(\alpha_{\mathrm{c}}, \alpha_{\mathrm{h}}, \phi) &= \frac{k_{\mathrm{B}}T}{b^3}\left[(1-(1+\alpha_{\mathrm{c}})\phi)\log(1-(1+\alpha_{\mathrm{c}})\phi) - \chi_0\phi^2 - \chi_{\mathrm{m}}\alpha_{\mathrm{h}}^2\phi^2 \right. \\ &\quad \left. -\mu_{\mathrm{h}}\alpha_{\mathrm{h}}\phi - \mu_{\mathrm{c}}\alpha_{\mathrm{c}}\phi\right]. \end{aligned} \tag{S34}$$

in the following discussion and in the main article.

#### S2.3 Chemical potential of linker histones

At the minimum of the free energy, the variation of the free energy with respect to the occupancy $\alpha_{\mathrm{h}}$ is zero,

$$\begin{aligned} \frac{\delta F}{\delta \alpha_{\mathrm{h}}} &= \frac{k_{\mathrm{B}}T}{b^3}\phi\left[\log\alpha_{\mathrm{h}} + 1 - \log(1-\alpha_{\mathrm{c}}-\alpha_{\mathrm{h}})\log(1-\alpha_{\mathrm{c}}-\alpha_{\mathrm{h}}) - 1 \right. \\ &\quad \left. +\epsilon_{\mathrm{h}} - \mu_{\mathrm{h}} - 2\chi_{\mathrm{m}}\alpha_{\mathrm{h}}\phi\right] = 0. \end{aligned} \tag{S35}$$

This leads to the relationship

$$\mu_{\mathrm{h}} = \log\alpha_{\mathrm{h}} - \log(1-\alpha_{\mathrm{c}}-\alpha_{\mathrm{h}}) + \epsilon_{\mathrm{h}} - 2\chi_{\mathrm{m}}\alpha_{\mathrm{h}}\phi. \tag{S36}$$

Eq. (S36) represents the fact that the occupancy $\alpha_{\mathrm{h}}$ is determined by the equality of the local chemical potential $\mu_{\mathrm{h}}$ of linker histones. Eq. (S36) is effective both in the central and peripheral regions. For cases in which the diffusion of linker histone is fast enough, the chemical potential $\mu_{\mathrm{h}}$ is uniform.

#### S2.4 Chemical potential of condensin complexes

The ATP hydrolysis is involved in the loading of condensin complexes to DNA and thus it cannot be treated by using the equality of chemical potential. However, it is instructive to calculate the variation of the free energy with respect to the occupancy $\alpha_{\rm c}$,

$$\frac{\delta F}{\delta \alpha_{\rm c}} = \frac{k_{\rm B}T}{b^3}\phi \left[\log \alpha_{\rm c} - \log(1-\alpha_{\rm c}-\alpha_{\rm h}) + \epsilon_{\rm c} - \log(1-(1+\alpha_{\rm c})\phi) - 1 - \mu_{\rm c}\right]. \quad \text{(S37)}$$

The equilibrium of the loading and unloading of condensin complexes, $\frac{\delta F}{\delta \alpha_{\rm c}} = 0$, leads to the form

$$\mu_{\rm c} + \log(1-\alpha_{\rm c}-\alpha_{\rm h}) + \log(1-(1+\alpha_{\rm c})\phi) = \epsilon_{\rm c} - 1 + \log \alpha_{\rm c} \quad \text{(S38)}$$

Putting both sides of eq. (S38) at the shoulder of exponential leads to

$$\mathrm{e}^{\mu_{\rm c}}(1-\alpha_{\rm c}-\alpha_{\rm h})(1-(1+\alpha_{\rm c})\phi) = \mathrm{e}^{\epsilon_{\rm c}-1}\alpha_{\rm c}. \quad \text{(S39)}$$

Eq. (S39) is indeed the steady state solution of the kinetic equation

$$\frac{d\alpha_{\rm c}}{dt} = k_0\phi_{\rm c0}(1-\alpha_{\rm c}-\alpha_{\rm h})(1-(1+\alpha_{\rm c})\phi) - k_0\mathrm{e}^{\epsilon_{\rm c}-1}\alpha_{\rm c} \quad \text{(S40)}$$

where we used eq. (S32) with $\phi_{\rm c0} \ll 1$ to derive this equation. $k_0$ is the rate constant that accounts for the binding and unbinding of condensin complexes. The factors $(1-\alpha_{\rm c}-\alpha_{\rm h})$ and $(1-(1+\alpha_{\rm c})\phi)$ result from the mutual exclusion of condensin complexes and linker histones and from the osmotic pressure, respectively, see also eq. (S31). These contributions are also involved even when we take into account the active nature of the loading of condensin complexes. For case of $\alpha_{\rm c} \ll 1$, eq. (S40) has an approximate form

$$\frac{d\alpha_{\rm c}}{dt} = k_0\phi_{\rm c0}(1-\alpha_{\rm h})(1-\phi) - k_0\mathrm{e}^{\epsilon_{\rm c}-1}\alpha_{\rm c}. \quad \text{(S41)}$$

#### S2.5 Osmotic pressure in peripheral brush region

We assume that the DNA volume fraction $\phi_{\rm b}$ is uniform in the peripheral brush region. At the minimum of the free energy, the first partial derivative of the free energy with respect to the DNA volume fraction $\phi$ is zero

$$\frac{\partial}{\partial \phi} F = \frac{N_{\rm b}}{\phi_{\rm b}^2}\left[-3\frac{k_{\rm B}T}{b^3}\frac{(\sigma b^2)^2}{\phi_{\rm b}} + \phi_{\rm b}^2 \frac{\partial}{\partial \phi_{\rm b}}\left(\frac{f_{\rm b}(\alpha_{\rm c},\alpha_{\rm h},\phi_{\rm b})}{\phi_{\rm b}}\right)\right] = 0, \tag{S42}$$

where

$$f_{\rm b}(\alpha_{\rm c},\alpha_{\rm h},\phi_{\rm b}) = f_{\rm lop}(\phi_{\rm b}) + f_{\rm sol}(\alpha_{\rm c},\alpha_{\rm h},\phi_{\rm b}) + f_{\rm bnd}(\alpha_{\rm c},\alpha_{\rm h},\phi_{\rm b}) \tag{S43}$$

is the free energy of the brush. This leads to the balance of osmotic pressure

$$-\frac{3k_{\rm B}T}{b^3}\frac{(\sigma b^2)^2}{\phi_{\rm b}} + \Pi_{\rm sol}(\phi_{\rm b}) = 0 \tag{S44}$$

with

$$\frac{\Pi_{\rm sol}(\phi) b^3}{k_{\rm B}T} = -\log(1-(1+\alpha_{\rm c})\phi) - (1+\alpha_{\rm c})\phi - \chi_0\phi^2 - \chi_{\rm m}\alpha_{\rm h}^2\phi^2. \tag{S45}$$

#### S2.6 Osmotic pressure in central gel region

We assume that the DNA volume fraction $\phi_{\rm g}$ is uniform in the central gel region. The osmotic pressure $\Pi_{\rm in}$ of the central gel region is derived by using the thermodynamic relationship

$$\begin{aligned}\Pi_{\rm in} &= -\frac{\partial F}{\partial V_{\rm in}} \\ &= \phi_{\rm g}^2 \frac{\partial}{\partial \phi_{\rm g}}\left(\frac{F}{b^3(N-N_{\rm b})}\right),\end{aligned} \tag{S46}$$

where we used eq. (S15) to derive the last equation.

The derivative with respect to the free energy of the central gel region is calculated as

$$\begin{aligned}
\Pi_{\rm g} &= \phi_{\rm g}^2 \frac{\partial}{\partial \phi_{\rm g}} \left( \frac{F_{\rm g}}{b^3(N-N_{\rm b})} \right) \\
&= \phi_{\rm g}^2 \frac{\partial}{\partial \phi_{\rm g}} \left( \frac{f_{\rm g}}{\phi_{\rm g}} \right) \\
&= \phi_{\rm g}^2 \frac{\partial}{\partial \phi_{\rm g}} \left[ \frac{3}{2} \frac{G_0}{\phi_0^{1/3}} n_{\rm e}^{-1} \frac{\phi_{\rm g}^{-2/3}}{(1-n_{\rm b})^{1/3}} + \frac{f_{\rm sol}(\alpha_{\rm c}, \alpha_{\rm h}, \phi_{\rm g}) + f_{\rm bnd}(\alpha_{\rm c}, \alpha_{\rm h}, \phi_{\rm g})}{\phi_{\rm g}} \right] \\
&= -\frac{G_0}{\phi_0^{1/3}} n_{\rm e}^{-1} \frac{\phi_{\rm g}^{1/3}}{(1-n_{\rm b})^{1/3}} + \Pi_{\rm sol}(\phi_{\rm g}), \qquad \text{(S47)}
\end{aligned}$$

where

$$f_{\rm g}(\alpha_{\rm c}, \alpha_{\rm h}, \phi_{\rm g}) = f_{\rm ent}(\phi_{\rm g}) + f_{\rm sol}(\alpha_{\rm c}, \alpha_{\rm h}, \phi_{\rm g}) + f_{\rm bnd}(\alpha_{\rm c}, \alpha_{\rm h}, \phi_{\rm g}) \qquad \text{(S48)}$$

is the free energy per unit volume in the central gel region. Eq. (S47) is an extension of the osmotic pressure of polymer gel by taking into account the fact that polymers can slide through effective crosslinks due to the entanglement. The derivative with respect to the free energy of the peripheral brush region is calculated as

$$\begin{aligned}
\Delta\Pi_{\rm g} &= \phi_{\rm g}^2 \frac{\partial}{\partial \phi_{\rm g}} \left( \frac{F_{\rm b}}{b^3(N-N_{\rm b})} \right) \\
&= \frac{2}{3} \frac{k_{\rm B}T}{b^3} \frac{N_{\rm b}}{N-N_{\rm b}} \frac{\phi_{\rm g}}{\phi_{\rm b}} \frac{3(\sigma b^2)^2}{\phi_{\rm b}} \\
&= \frac{2h}{r_{\rm in}} \Pi_{\rm sol}(\phi_{\rm b}). \qquad \text{(S49)}
\end{aligned}$$

To derive the last form of eq. (S49), we used

$$\frac{h}{r_{\rm in}} = \frac{1}{3} \frac{N_{\rm b}}{N-N_{\rm b}} \frac{\phi_{\rm g}}{\phi_{\rm b}}, \qquad \text{(S50)}$$

which is derived by using eqs. (S15) and (S16). Eq. (S49) has a similar form to eq. (S12), implying that eq. (S49) is effective only for $h < r_{\rm in}/2$. We use

$$\Delta\Pi_{\rm g} = \Pi_{\rm sol}(\phi_{\rm b}) \tag{S51}$$

for $h > r_{\rm in}/2$.

The osmotic pressure $\Pi_{\rm in}$ in the central gel region thus has the form

$$\begin{aligned} \Pi_{\rm in} &= \Pi_{\rm g} + \Delta\Pi_{\rm g} \\ &= -\frac{G_0}{\phi_0^{1/3}} n_{\rm e}^{-1} \frac{\phi_{\rm g}^{1/3}}{(1-n_{\rm b})^{1/3}} + \Pi_{\rm sol}(\phi_{\rm g}) + \Delta\Pi_{\rm g}. \end{aligned} \tag{S52}$$

#### S2.7 DNA tension for extruding DNA from elastically effective to ineffective regions

The DNA tension that counteracts the extrusion from the elastically effective to ineffective regions is derived by the principle of virtual displacement,

$$f_{\rm in} b \Delta N_{\rm gl} = \Delta F, \tag{S53}$$

where $\Delta N_{\rm gl}$ is the number of DNA units extruded by all the condensin complexes in the central gel region. In the time scale of one step of loop extrusion, the solvent does not go out from the central gel region and thus the volume in the central gel region is constant. The number of DNA units in the central gel region does not change with this process. The DNA tension thus has the form

$$\begin{aligned} f_{\rm in} b &= \left.\frac{\partial F}{\partial N_{\rm gl}}\right|_{N_{\rm b}, V_{\rm in}} \\ &= -\left.\frac{\partial}{\partial n_{\rm e}}\left(\frac{F}{N}\right)\right|_{N_{\rm b}, V_{\rm in}} \\ &= \frac{3}{2}\frac{G_0 b^3}{\phi_0^{1/3}}\frac{1}{n_{\rm e}^2}\frac{\phi_{\rm g}^{-2/3}}{(1-n_{\rm b})^{-2/3}}. \end{aligned} \tag{S54}$$

The second form of eq. (S54) is derived by using eqs. (S14) and (S21).

To understand the physical meaning of eq. (S54), it is instructive to think of the case in which DNA is a 'melt' in the reference state, where

$$G_0 = \frac{\phi_0}{b^3 N_{e0}} k_B T. \tag{S55}$$

in this case, eq. (S54) is rewritten as

$$f_{in} b = \frac{1}{2} k \Delta x^2 \tag{S56}$$

with

$$k = \frac{3 k_B T}{b^2} \tag{S57}$$

$$\Delta x = \frac{\lambda_s b N_{e0}^{1/2}}{N_e}. \tag{S58}$$

Flexible polymers are often treated as beads connected by springs (the spring-bead model). $k$ is the spring constant of a spring and $\Delta x$ is the length of the spring. Eq. (S56) is thus the elastic energy per spring.

#### S2.8 DNA tension for extruding DNA from central to peripheral regions

The DNA tension that counteracts the extrusion from the central to peripheral regions is derived by using the principle of virtual displacement,

$$f_{ex} b \Delta N_b = \Delta F. \tag{S59}$$

In the time scale of one step of loop extrusion, the solvent does not go out from the central gel region and thus the volume in the central gel region is constant. The DNA tension thus has the form

$$\begin{aligned} f_{\mathrm{ex}} b &= \left.\frac{\partial F}{\partial N_{\mathrm{b}}}\right|_{V_{\mathrm{in}},V_{\mathrm{ex}}} \\ &= -(\mu_{\mathrm{ex}}^{\mathrm{g}} - \mu_{\mathrm{ex}}^{\mathrm{b}}). \end{aligned} \tag{S60}$$

The chemical potential $\mu_{\mathrm{ex}}^{\mathrm{g}}$ of DNA units in the central gel region has the form

$$\begin{aligned} \mu_{\mathrm{ex}}^{\mathrm{g}} &= \left.\frac{\partial F_{\mathrm{g}}}{\partial N_{\mathrm{eff}}}\right|_{V_{\mathrm{in}},V_{\mathrm{ex}}} \\ &= -\left.\frac{\partial}{\partial n_{\mathrm{b}}}\left((1-n_{\mathrm{b}})\frac{f_{\mathrm{g}}(\phi_{\mathrm{g}},n_{\mathrm{b}})b^3}{\phi_{\mathrm{g}}}\right)\right|_{V_{\mathrm{in}},V_{\mathrm{ex}}} \\ &= \frac{G_0 b^3}{\phi_0^{1/3}}\frac{(1-n_{\mathrm{b}})^{-1/3}}{\phi_{\mathrm{g}}^{2/3}}\frac{1}{n_{\mathrm{e}}} + \frac{3}{2}\frac{G_0 b^3}{\phi_0^{1/3}}\frac{(1-n_{\mathrm{b}})^{2/3}}{\phi_{\mathrm{g}}^{2/3}}\frac{1}{n_{\mathrm{e}}^2} \\ &\quad + \frac{f_{\mathrm{sol}}(\alpha_{\mathrm{c}},\alpha_{\mathrm{h}},\phi_{\mathrm{g}})b^3 + f_{\mathrm{bnd}}(\alpha_{\mathrm{c}},\alpha_{\mathrm{h}},\phi_{\mathrm{g}})b^3}{\phi_{\mathrm{g}}} + \frac{\Pi_{\mathrm{g}}(\phi_{\mathrm{g}})b^3}{\phi_{\mathrm{g}}} \\ &= -\frac{1}{2}\frac{G_0 b^3}{\phi_0^{1/3}}\frac{(1-n_{\mathrm{b}})^{-1/3}}{\phi_{\mathrm{g}}^{2/3}}\frac{1}{n_{\mathrm{e}}} + \frac{3}{2}\frac{G_0 b^3}{\phi_0^{1/3}}\frac{(1-n_{\mathrm{b}})^{2/3}}{\phi_{\mathrm{g}}^{2/3}}\frac{1}{n_{\mathrm{e}}^2} \\ &\quad + \frac{f_{\mathrm{g}}(\alpha_{\mathrm{c}},\alpha_{\mathrm{h}},\phi_{\mathrm{g}})b^3}{\phi_{\mathrm{g}}} + \frac{\Pi_{\mathrm{g}}(\phi_{\mathrm{g}})b^3}{\phi_{\mathrm{g}}}. \end{aligned} \tag{S61}$$

The second form of eq. (S61) is derived by using eqs. (S14) and (S21). The chemical potential $\mu_{\mathrm{ex}}^{\mathrm{b}}$ of DNA units in the peripheral brush region has the form

$$\begin{aligned} \mu_{\mathrm{ex}}^{\mathrm{b}} &= \left.\frac{\partial F_{\mathrm{b}}}{\partial N_{\mathrm{b}}}\right|_{V_{\mathrm{in}},V_{\mathrm{ex}}} \\ &= \left.\frac{\partial}{\partial n_{\mathrm{b}}}\left(\frac{f_{\mathrm{b}}(\phi_{\mathrm{g}},\phi_{\mathrm{b}},n_{\mathrm{b}})b^3 n_{\mathrm{b}}}{\phi_{\mathrm{b}}}\right)\right|_{V_{\mathrm{in}},V_{\mathrm{ex}}} \\ &= \frac{f_{\mathrm{b}}(\phi_{\mathrm{g}},\phi_{\mathrm{b}},n_{\mathrm{b}})b^3}{\phi_{\mathrm{b}}} - \frac{\Delta\Pi_{\mathrm{g}}}{\phi_{\mathrm{g}}}, \end{aligned} \tag{S62}$$

where we used eqs. (S44) and (S49) to derive the last form of eq. (S62).

### S3 Loop extrusion dynamics

#### S3.1 DNA extrusion from central to peripheral regions

The number $N_{\rm bl}$ of DNA units in a loop at the peripheral region increases due to the loop extrusion by condensin complexes at the interface between the central and peripheral regions. The number $N_{\rm bl}$ of DNA units decreases when all the condensin complexes at the interface are unloaded. The probability $q_n$ with respect to the number $n$ of condensin complexes at the interface follows the time evolution equation

$$\frac{d}{dt}q_n(t) = -J_{n+1}(t) + J_n(t) \tag{S63}$$

with

$$J_n(t) = 2v_0\alpha_{\rm c}q_{n-1} - \frac{n}{\tau_{\rm ex}}q_n. \tag{S64}$$

The first term of eq. (S64) represents the fact that there are $\alpha_{\rm c}N_{\rm bl}$ condensin complexes in a loop and these complexes are transported towards the interface with the rate $2v_0/N_{\rm bl}$ and the second term of this equation represents the unloading among $n$ condensin complexes at the interface.

The probability $q_n^{\rm s}$ in the steady state, $J_n(t) = 0$, is derived as

$$q_n^{\rm s} = \frac{2v_0\tau_{\rm ex}\alpha_{\rm c}}{n}q_{n-1}^{\rm s} \tag{S65}$$

$$= \frac{(2v_0\tau_{\rm ex}\alpha_{\rm c})^{n-1}}{n!}q_1^{\rm s}. \tag{S66}$$

For cases in which there are at least one condensin complex at the interface, the probability is normalized as

$$\sum_{n=1}^{\infty} q_n^{\rm s} = 1. \tag{S67}$$

By using eq. (S67), the probability $q_1^{\rm s}$ that there is only one condensin complex at the interface is derived as

$$q_1^{\rm s} = \frac{2v_0\tau_{\rm ex}\alpha_{\rm c}}{{\rm e}^{2v_0\tau_{\rm ex}\alpha_{\rm c}} - 1}. \tag{S68}$$

The rate with which all the condenxin complexes are unloaded from the interface is $\frac{q_1^{\rm s}}{\tau_{\rm ex}}$. Because the average number of DNA units between two bound condensin complexes is $\alpha_{\rm c}^{-1}$, the number of DNA units decreases from the DNA loop per unit time is derived as

$$\alpha_{\rm c}^{-1}\frac{q_1^{\rm s}}{\tau_{\rm ex}} = \frac{2v_0}{{\rm e}^{2v_0\tau_{\rm ex}\alpha_{\rm c}} - 1}. \tag{S69}$$

The reeling of DNA, eq. (S69), is significant for $t > \tau_{\rm ex}$, when condensin complexes start to unload. The time evolution of the number $N_{\rm bl}$ of the DNA units in a loop at the peripheral region has the from

$$\frac{dN_{\rm bl}}{dt} = 2v_{\rm ex} \tag{S70}$$

for $t < \tau_{\rm ex}$ and

$$\frac{dN_{\rm bl}}{dt} = 2v_{\rm ex} - \frac{2v_0}{{\rm e}^{2v_0\tau_{\rm ex}\alpha_{\rm c}} - 1} \tag{S71}$$

for $t > \tau_{\rm ex}$.

#### S3.2 DNA extrusion from elastically effective to ineffective regions

The time evolution equation of the number $N_{\mathrm{gl}}$ of DNA units in the elastically ineffective loops at the central gel region has the form

$$\frac{dN_{\mathrm{gl}}}{dt} = -\frac{N_{\mathrm{gl}}}{\tau_{\mathrm{ex}}} + 2v_{\mathrm{in}}n_{\mathrm{gl}}. \tag{S72}$$

The first and second terms of eq. (S72) are the changes of the number of DNA units in elastically ineffective region due to the unloading of condensin complexes and the loop extrusion, respectively. $v_{\mathrm{in}}$ and $n_{\mathrm{gl}}$ are the extrusion rate and number of condensin complexes that perform the extrusion from the elastically effective to ineffective regions. The time evolution equation of the number $n_{\mathrm{gl}}$ of condensin complexes has the form

$$\frac{dn_{\mathrm{gl}}}{dt} = -\frac{n_{\mathrm{gl}}}{\tau_{\mathrm{ex}}} + k_{\mathrm{on}}\phi_{\mathrm{c0}}(1-\alpha_{\mathrm{h}})(1-\phi_{\mathrm{g}})N_{\mathrm{eff}}. \tag{S73}$$

The first and second terms of eqs. (S73) are the unloading and loading rate of condensin complexes to the elastically effective region, respectively. $k_{\mathrm{on}}$ is the rate constant that accounts for the loading of condensin complexes. The mutual extrusion between condensin complexes and linker histones as well as the contribution of the osmotic pressure are takein into account via the factors $1-\alpha_{\mathrm{h}}$ and $1-\phi_{\mathrm{g}}$ in the second term of this equation, see also eq. (S41).

The solution of eq. (S73) has the form

$$n_{\mathrm{gl}}(t) = \int_0^t dt' \, \frac{\alpha_{\mathrm{c}}^{\mathrm{s}}(t')}{\tau_{\mathrm{ex}}} N_{\mathrm{eff}}(t')\mathrm{e}^{-(t-t')/\tau_{\mathrm{ex}}} \tag{S74}$$

with

$$\alpha_{\mathrm{c}}^{\mathrm{s}}(t) = k_{\mathrm{on}}\tau_{\mathrm{ex}}(1-\alpha_{\mathrm{h}}(t))(1-\phi_{\mathrm{g}}(t)). \tag{S75}$$

The solution of eq. (S72) has the form

$$
\begin{aligned}
N_{\mathrm{gl}}(t) &= \int_0^t dt''\, 2v_{\mathrm{in}}(t'')n_{\mathrm{gl}}(t'')\mathrm{e}^{-(t-t'')/\tau_{\mathrm{ex}}} \\
&= \int_0^t dt'' \int_0^{t''} dt'\, 2v_{\mathrm{in}}(t'')\frac{\alpha_{\mathrm{c}}^{\mathrm{s}}(t')}{\tau_{\mathrm{ex}}}N_{\mathrm{eff}}(t')\mathrm{e}^{-(t-t')/\tau_{\mathrm{ex}}} \\
&= \int_0^t dt' \left(\int_{t'}^t dt''\, 2v_{\mathrm{in}}(t'')\right)\frac{\alpha_{\mathrm{c}}^{\mathrm{s}}(t')}{\tau_{\mathrm{ex}}}N_{\mathrm{eff}}(t')\mathrm{e}^{-(t-t')/\tau_{\mathrm{ex}}}. \qquad \text{(S76)}
\end{aligned}
$$

The integrand in the last form of eq. (S76) suggests that among $\frac{\alpha_{\mathrm{c}}^{\mathrm{s}}(t')}{\tau_{\mathrm{ex}}}N_{\mathrm{eff}}(t')\Delta t'$ condensin complexes loaded to DNA at time $t'$, only the fraction of $\mathrm{e}^{-(t-t')/\tau_{\mathrm{ex}}}$ remains at the DNA and that these condensin complexes extrude $\int_{t'}^t dt''\, 2v_{\mathrm{in}}(t'')$ DNA units.

For cases in which $v_{\mathrm{in}}(t)$, $\alpha_{\mathrm{c}}^{\mathrm{s}}(t)$, and $N_{\mathrm{eff}}(t)$ change only slowly with time, eq. (S76) has an approximate form

$$
N_{\mathrm{gl}}(t) \approx 2v_{\mathrm{in}}(t)\tau_{\mathrm{ex}}\alpha_{\mathrm{c}}^{\mathrm{s}}(t)N_{\mathrm{eff}}(t)T(t) \qquad \text{(S77)}
$$

with the time factor

$$
\begin{aligned}
T(t) &= \int_0^t \frac{dt'}{\tau_{\mathrm{ex}}}\int_{t'}^t \frac{dt''}{\tau_{\mathrm{ex}}}\,\mathrm{e}^{-(t-t')/\tau_{\mathrm{ex}}} \\
&= 1-\left(1+\frac{t}{\tau_{\mathrm{ex}}}\right)\mathrm{e}^{-t/\tau_{\mathrm{ex}}}. \qquad \text{(S78)}
\end{aligned}
$$

For simplicity, we assume the linear relationship between the extrusion rate $v_{\mathrm{in}}$ and the DNA tension $f_{\mathrm{in}}$,

$$
v_{\mathrm{in}} = v_0\left(1-\frac{f_{\mathrm{in}}}{f_{\mathrm{s}}}\right), \qquad \text{(S79)}
$$

where $f_{\mathrm{s}}$ is the stall force of condensin complex. Substituting eq. (S54) into eq. (S79) leads to the form

$$v_{\rm in} = v_0 \left(1 - \zeta \frac{(1-n_{\rm b})^2}{n_{\rm e}^2}\right) \tag{S80}$$

with

$$\zeta = \frac{3}{2} \frac{G_0 b^3}{\phi_0^{1/3} f_{\rm s}} \frac{\phi_{\rm g}^{-2/3}}{(1-n_{\rm b})^{4/3}}. \tag{S81}$$

By substituting eq. (S80) into eq. (S14), one derives the relationship

$$\frac{1-n_{\rm b}}{n_{\rm e}} = 1 + 2v_0\tau_{\rm ex}\alpha_{\rm c}^{\rm s}(t)T(t)\left(1 - \zeta \frac{(1-n_{\rm b})^2}{n_{\rm e}^2}\right). \tag{S82}$$

The fraction $n_{\rm e}$ of DNA units in the elastically effective region is thus derived as

$$\frac{n_{\rm e}}{1-n_{\rm b}} = \frac{1 + \sqrt{1 + 8v_0\tau_{\rm ex}\alpha_{\rm c}^{\rm s}(t)T(t)\zeta(1 + 2v_0\tau_{\rm ex}\alpha_{\rm c}^{\rm s}(t)T(t))}}{2(1 + 2v_0\tau_{\rm ex}\alpha_{\rm c}^{\rm s}(t)T(t))} \tag{S83}$$

by solving eq. (S82) with respect to $n_{\rm e}$.

For the case of small DNA tension, $\zeta \to 0$, the fraction $n_{\rm e}$ has an asymptotic form

$$n_{\rm e} \to \frac{1-n_{\rm b}}{1 + 2v_0\tau_{\rm ex}\alpha_{\rm c}^{\rm s}(t)T(t)}. \tag{S84}$$

For the case of large DNA tension, $\zeta \to 1$, the fraction $n_{\rm e}$ has an asymptotic form

$$n_{\rm e} \to 1 - n_{\rm b}. \tag{S85}$$

The extrusion rate is zero in the latter limit, see eq. (S80).

#### S4 Relaxation time after one step of loop extrusion

The time evolution of the radius $r_{\rm in}$ of the central gel region has the form

$$\frac{dr_{\rm in}}{dt} = -\kappa(\phi_{\rm b})\frac{p_{\rm in}}{h} \tag{S86}$$

with

$$\kappa(\phi_{\rm b}) = \kappa_0\frac{(1-\phi_{\rm b})^2}{\sigma b^2}, \tag{S87}$$

see eq. (30) in the main article. We treat cases in which condensin complexes extrude $\Delta N_{\rm b}$ DNA units in one step of loop extrusion process. The hydrostatic pressure increases at the moment of this process has the form

$$\Delta p_{\rm in} = -\left.\frac{\partial \Pi_{\rm in}}{\partial N_{\rm b}}\right|_{V_{\rm in}}\Delta N_{\rm b}, \tag{S88}$$

where $|_{V_{\rm in}}$ in the left side represents the constraint of constant volume $v_{\rm in}$ of the central region. The hydrostatic pressure $\Delta p_{\rm in}$ drives solvent flow towards the external solution and the radius $r_{\rm in}$ changes by

$$\Delta r_{\rm in} = \frac{\partial r_{\rm in}}{\partial N_{\rm b}}\Delta N_{\rm b}, \tag{S89}$$

where the derivative in the left side assumes the local equilibrium. This change happens during the relaxation time $\Delta\tau_{\rm r}$. Eq. (S86) suggests the relationship,

$$\frac{\Delta r_{\rm in}}{\Delta \tau_{\rm r}} = -\kappa(\phi_{\rm b})\frac{\Delta p_{\rm in}}{h}. \tag{S90}$$

By substituting eqs. (S88) and (S89) into (S90), the relaxation time $\Delta\tau_{\mathrm{r}}$ per step is derived as

$$\begin{aligned}\Delta\tau_{\mathrm{r}} &= -\frac{1}{\kappa(\phi_{\mathrm{b}})}\frac{\Delta r_{\mathrm{in}}}{\Delta p_{\mathrm{in}}}h \\ &= \frac{1}{\kappa(\phi_{\mathrm{b}})}\frac{\frac{\partial r_{\mathrm{in}}}{\partial N_{\mathrm{b}}}}{\left.\frac{\partial \Pi_{\mathrm{in}}}{\partial N_{\mathrm{b}}}\right|_{V_{\mathrm{in}}}}h. \end{aligned} \tag{S91}$$

By using the relationships

$$r_{\mathrm{in}} = r_0\left(\frac{1-n_{\mathrm{b}}}{\phi_{\mathrm{g}}}\right)^{1/3} \tag{S92}$$

$$\tau_{\mathrm{r}} = \frac{r_0^2 b^3}{\kappa_0 k_{\mathrm{B}} T}, \tag{S93}$$

eq. (S91) is also rewritten as

$$\frac{\Delta\tau_{\mathrm{r}}}{\tau_{\mathrm{r}}} = \frac{\sigma b^2}{(1-\phi_{\mathrm{b}})^2}\frac{\frac{\partial}{\partial N_{\mathrm{b}}}\left(\frac{1-n_{\mathrm{b}}}{\phi_{\mathrm{g}}}\right)^{1/3}}{\left.\frac{\partial}{\partial N_{\mathrm{b}}}\left(\frac{\Pi_{\mathrm{in}} b^3}{k_{\mathrm{B}} T}\right)\right|_{V_{\mathrm{in}}}}\frac{h}{r_0}. \tag{S94}$$

### S5 Asymptotic analysis

#### S5.1 Swollen phase in peripheral region

In the steady state, the occupancy of condensin complexes has the form

$$\alpha_{\mathrm{cb}} = \frac{\alpha_{\mathrm{c0}}(1-\alpha_{\mathrm{hb}})(1-\phi_{\mathrm{b}})}{1+\frac{2v_0\tau_{\mathrm{ex}}}{N_{\mathrm{bl}}}}, \tag{S95}$$

see eq. (29) in the main article. We use the subscript "cb" and "hb" to indicate the quantities of condensin complexes and linker histones in the peripheral region, respectively

. Substituting eq. (S95) into eq. (S36) leads to the form

$$\log \alpha_{\mathrm{hb}} - \log(1-\alpha_{\mathrm{hb}}) - \log\left(1 - \frac{\alpha_{\mathrm{c0}}(1-\phi_{\mathrm{b}})}{1+\frac{2v_0\tau_{\mathrm{ex}}}{N_{\mathrm{bl}}}}\right) - 2\chi_{\mathrm{m}}\phi_{\mathrm{b}}\alpha_{\mathrm{hb}} + \epsilon_{\mathrm{h}} - \mu_{\mathrm{h}} = 0. \tag{S96}$$

The occupancies of condensin complexes and linker histones are derived by solving eqs. (S95) and (S96) as functions of the DNA volume fraction $\phi_{\mathrm{b}}$. The DNA volume fraction $\phi_{\mathrm{b}}$ is derived by using eq. (S44).

For the case of the swollen state, $\alpha_{\mathrm{hb}} \ll 1$, eq. (S96) has an approximate form

$$\log \alpha_{\mathrm{hb}} - \log\left(1 - \frac{\alpha_{\mathrm{c0}}(1-\phi_{\mathrm{b}})}{1+\frac{2v_0\tau_{\mathrm{ex}}}{N_{\mathrm{bl}}}}\right) + \epsilon_{\mathrm{h}} - \mu_{\mathrm{h}} = 0, \tag{S97}$$

where we omitted higher order terms with respect to $\alpha_{\mathrm{hb}}$. This leads to the form

$$\alpha_{\mathrm{hb}} = \left(1 - \frac{\alpha_{\mathrm{c0}}(1-\phi_{\mathrm{b}})}{1+\frac{2v_0\tau_{\mathrm{ex}}}{N_{\mathrm{bl}}}}\right) \mathrm{e}^{-(\epsilon_{\mathrm{h}}-\mu_{\mathrm{h}})}. \tag{S98}$$

Similarly, eq. (S95) is simplied as

$$\alpha_{\mathrm{cb}} = \frac{\alpha_{\mathrm{c0}}(1-\phi_{\mathrm{b}})}{1+\frac{2v_0\tau_{\mathrm{ex}}}{N_{\mathrm{bl}}}}(1-\mathrm{e}^{-(\epsilon_{\mathrm{h}}-\mu_{\mathrm{h}})}). \tag{S99}$$

By using $\alpha_{\mathrm{h}} \ll 1$, eq. (S45) is expanded in a power series of $(1+\alpha_{\mathrm{c}})\phi$ as

$$\begin{aligned} \frac{\Pi_{\mathrm{sol}}(\phi)b^3}{k_{\mathrm{B}}T} &= -\log(1-(1+\alpha_{\mathrm{c}})\phi) - (1+\alpha_{\mathrm{c}})\phi - \chi_0\phi^2 \\ &= \frac{1}{2}((1+\alpha_{\mathrm{c}})^2 - 2\chi_0)\phi^2 + \frac{1}{3}(1+\alpha_{\mathrm{c}})^3\phi^3 + \cdots . \end{aligned} \tag{S100}$$

The DNA volume fraction is derived by solving $\frac{\Pi_{\mathrm{sol}}(\phi)b^3}{k_{\mathrm{B}}T} = 0$ respect to $\phi$. Similarly, eqs. (S19) and (S62) are expanded as

$$\begin{aligned} \frac{f_{\mathrm{b}}(\alpha_{\mathrm{c}},\alpha_{\mathrm{h}},\phi_{\mathrm{b}})}{\phi_{\mathrm{b}}k_{\mathrm{B}}T} &= \frac{3}{2}\frac{(\sigma b^2)^2}{\phi_{\mathrm{b}}^2} + \frac{1}{2}((1+\alpha_{\mathrm{c}})^2 - 2\chi_0)\phi_{\mathrm{b}} + \frac{1}{3}(1+\alpha_{\mathrm{c}})^3\phi_{\mathrm{b}}^2 - 1 \\ &\quad +\alpha_{\mathrm{c}}\log\alpha_{\mathrm{c}} + (1-\alpha_{\mathrm{c}})\log(1-\alpha_{\mathrm{c}}) + (\epsilon_{\mathrm{c}} - \mu_{\mathrm{c}} - 1)\alpha_{\mathrm{c}} \end{aligned} \tag{S101}$$

In the good solvent condition, $(1+\alpha_{\rm cb})^2-2\chi_0>0$, the DNA volume fraction an approximate form

$$\phi_{\rm b} \;=\; \left(\frac{6(\sigma b^2)^2}{(1+\alpha_{\rm cb}(0))^2-2\chi_0}\right)^{1/3}, \tag{S102}$$

where $\alpha_{\rm cb}(0)$ is the value of condensin occupancy with $\phi_{\rm b}=0$, see eq. (S99). In the $\theta$-solvent condition, $(1+\alpha_{\rm cb})^2-2\chi_0\approx 0$, the DNA volume fraction has an approximate form

$$\phi_{\rm b} \;=\; \left(9(\sigma b^2)^2\right)^{1/4}, \tag{S103}$$

where we use the fact that the first order term $\alpha_{\rm c}\phi^3$ with respect to $\alpha_{\rm c}$ in the second term of eq. (S100) is eliminated by the $\phi$ dependent factor of the first order term $\alpha_{\rm c}\phi^2$ with respect to $\alpha_{\rm c}$ in the first term of eq. (S100). For both cases of the solvents, the DNA volume fraction $\phi_{\rm b}$ in the peripheral region is determined by the surface density $\sigma$ of the DNA loops. .

The lateral osmotic pressure is derived as

$$\frac{\Pi_{\rm sol}(\phi_{\rm b})b^3}{k_{\rm B}T} \;=\; \frac{1}{2}\left(6(\sigma b^2)^2\right)^{2/3}\left((1+\alpha_{\rm cb}(0))^2-2\chi_0\right)^{1/3} \tag{S104}$$

in the good solvent and

$$\frac{\Pi_{\rm sol}(\phi_{\rm b})b^3}{k_{\rm B}T} \;=\; \frac{1}{3}\left(9(\sigma b^2)^2\right)^{3/4} \tag{S105}$$

in the $\theta$-solvents. Eqs. (S104) and (S105) are derived by substituting eqs. (S102) and (S103) into eq. (S100), respectively.

#### S5.2 Condensed phase and phase separation in peripheral region

In the condensed phase, the occupancy of linker histones and DNA volume fraction are represented as

$$\alpha_{\mathrm{hb}}^{\mathrm{c}} = 1 - \delta\alpha_{\mathrm{hb}}^{\mathrm{c}} \tag{S106}$$

$$\phi_{\mathrm{b}}^{\mathrm{c}} = 1 - \delta\phi_{\mathrm{b}}^{\mathrm{c}}, \tag{S107}$$

with $\delta\alpha_{\mathrm{hb}} \ll 1$ and $\delta\phi_{\mathrm{b}} \ll 1$. The occupancy of condensin complexes is represented as

$$\alpha_{\mathrm{cb}}^{\mathrm{c}} = \frac{\alpha_{\mathrm{c0}}\delta\alpha_{\mathrm{hb}}^{\mathrm{c}}\delta\phi_{\mathrm{b}}^{\mathrm{c}}}{1+\frac{2v_0\tau_{\mathrm{ex}}}{N_{\mathrm{bl}}}} \approx 0 \tag{S108}$$

by substituting eqs. (S110) and (S117) into eq. (S95) and thus is as small as the second order term with respect to $\delta\alpha_{\mathrm{hb}}^{\mathrm{c}}$ and $\delta\phi_{\mathrm{b}}^{\mathrm{c}}$. By substituting eqs. (S110) - (S108) into eq. (S96) and expanding it in a power series of $\delta\alpha_{\mathrm{hb}}^{\mathrm{c}}$ and $\delta\phi_{\mathrm{b}}^{\mathrm{c}}$, we derive the form

$$-\log\delta\alpha_{\mathrm{hb}}^{\mathrm{c}} - 2\chi_{\mathrm{m}} + \epsilon_{\mathrm{h}} - \mu_{\mathrm{h}} = 0. \tag{S109}$$

By solving eq. (S109) with respect to $\delta\alpha_{\mathrm{hb}}^{\mathrm{c}}$, we derive the occupancy of linker histone in the form

$$\alpha_{\mathrm{hb}}^{\mathrm{c}} = 1 - \mathrm{e}^{-2\chi_{\mathrm{m}}+\epsilon_{\mathrm{h}}-\mu_{\mathrm{h}}}. \tag{S110}$$

The binodal line is derived by the equalities of the lateral osmotic pressure and chemical potential

$$\Pi_{\mathrm{sol}}(\phi_{\mathrm{b}}^{\mathrm{s}}) = \Pi_{\mathrm{sol}}(\phi_{\mathrm{b}}^{\mathrm{c}}) \tag{S111}$$

$$\frac{f_{\mathrm{b}}(\alpha_{\mathrm{cb}}^{\mathrm{s}},\alpha_{\mathrm{hb}}^{\mathrm{s}},\phi_{\mathrm{b}}^{\mathrm{s}})+\Pi_{\mathrm{sol}}(\phi_{\mathrm{b}}^{\mathrm{s}})}{\phi_{\mathrm{b}}^{\mathrm{s}}} = \frac{f_{\mathrm{b}}(\alpha_{\mathrm{cb}}^{\mathrm{c}},\alpha_{\mathrm{hb}}^{\mathrm{c}},\phi_{\mathrm{b}}^{\mathrm{c}})+\Pi_{\mathrm{sol}}(\phi_{\mathrm{b}}^{\mathrm{c}})}{\phi_{\mathrm{b}}^{\mathrm{c}}} \tag{S112}$$

between the two coexisting states, where the superscrits, s and c, indicate the quantitites of the swollen and condensed states, respectively. The surface density $\sigma$ in $f_{\rm b}(\alpha_{\rm cb}, \alpha_{\rm hb}, \phi_{\rm b})$ is derived by using

$$(\sigma b^2)^2 = \frac{1}{3}\phi_{\rm b}\frac{\Pi_{\rm sol}(\phi_{\rm b})b^3}{k_{\rm B}T}, \tag{S113}$$

see eq. (S44).

The lateral osmotic pressure in the condensed phase is derived as

$$\frac{\Pi_{\rm sol}(\phi_{\rm b}^{\rm c})b^3}{k_{\rm B}T} \approx -\log\delta\phi_{\rm b}^{\rm c} - 1 - \chi_0 - \chi_{\rm m} \tag{S114}$$

by substituting eqs. (S110) - (S108) into eq. (S45). The lateral osmotic pressure in the swollen phase is derived as

$$\frac{\Pi_{\rm sol}(\phi_{\rm b}^{\rm s})b^3}{k_{\rm B}T} \approx \frac{1}{2}\left((1+\alpha_{\rm cb}^{\rm s}(0))^2 - 2\chi_0\right)(\phi_{\rm b}^{\rm s})^2 \tag{S115}$$

in the good solvent condition, $(1+\alpha_{\rm cb}^{\rm s}(0))^2 - 2\chi_0 > 0$, and

$$\frac{\Pi_{\rm sol}(\phi_{\rm b}^{\rm s})b^3}{k_{\rm B}T} \approx \frac{1}{3}(\phi_{\rm b}^{\rm s})^3 \tag{S116}$$

in the $\theta$-solvent condition, $(1+\alpha_{\rm cb}^{\rm s}(0))^2 - 2\chi_0 \approx 0$, see eq. (S100). For both cases of the solvents, $\Pi_{\rm sol}(\phi_{\rm b}^{\rm s}) \approx 0$. Eq. (S111) leads to the form of the DNA volume fraction

$$\phi_{\rm b}^{\rm c} = 1 - {\rm e}^{-\chi_{\rm m}-\chi_0-1} \tag{S117}$$

at the binodal line.

By using eqs. (S108), (S110), and (S117), the free energy per DNA unit in the condensed phase is derived as

$$\frac{f_{\rm b}(\alpha_{\rm cb}^{\rm c}, \alpha_{\rm hb}^{\rm c}, \phi_{\rm b}^{\rm c}) b^3}{\phi_{\rm b}^{\rm c} k_{\rm B} T} = -\chi_0 - \chi_{\rm m} + \epsilon_{\rm h} - \mu_{\rm h}. \tag{S118}$$

In the good solvent condition, the free energy per DNA unit in the swollen phase has the form

$$\frac{f_{\rm b}(\alpha_{\rm cb}^{\rm s}, \alpha_{\rm hb}^{\rm s}, \phi_{\rm b}^{\rm s}) b^3}{\phi_{\rm b}^{\rm s} k_{\rm B} T} = \frac{3}{4} \left( (1 + \alpha_{\rm c}(0))^2 - 2\chi_0 \right) \phi_{\rm b}^{\rm s} - 1 + \bar{f}_{\rm c} \tag{S119}$$

with

$$\bar{f}_{\rm c} = \alpha_{\rm cb}^{\rm s} \log \alpha_{\rm cb}^{\rm s} + (1 - \alpha_{\rm cb}^{\rm s}) \log(1 - \alpha_{\rm cb}^{\rm s}) + (\epsilon_{\rm c} - \mu_{\rm c} - 1)\alpha_{\rm c}^{\rm s} \tag{S120}$$

Eq. (S119) is derived by substituting eq. (S113) into eq. (S101), see also eq. (S44). The term $\bar{f}_{\rm c}$ is expanded in a series of $\phi_{\rm b}$ as

$$\bar{f}_{\rm c} = \bar{f}_{\rm c0} + \bar{s}_{\rm c0} \phi_{\rm b} \tag{S121}$$

with

$$\bar{f}_{\rm c0} = \alpha_{\rm cb}^{\rm s}(0) \log \alpha_{\rm cb}^{\rm s}(0) + (\epsilon_{\rm c} - \mu_{\rm c} - 2)\alpha_{\rm cb}^{\rm s}(0) \tag{S122}$$

$$\bar{s}_{\rm c0} = -\alpha_{\rm cb}^{\rm s}(0) \log \alpha_{\rm cb}^{\rm s}(0) - (\epsilon_{\rm c} - \mu_{\rm c} - 1)\alpha_{\rm cb}^{\rm s}(0), \tag{S123}$$

where we also used $(1 - \alpha_{\rm cb}^{\rm s}) \log(1 - \alpha_{\rm cb}^{\rm s}) \approx -\alpha_{\rm cb}^{\rm s}$. The DNA volume fraction in the swollen phase is derived as

$$\phi_{\rm b}^{\rm s} = \frac{4}{5} \frac{1 - \bar{f}_{\rm c0} - \chi_0 - \chi_{\rm m} + \epsilon_{\rm h} - \mu_{\rm h}}{(1 + \alpha_{\rm cb}^{\rm s}(0))^2 - 2\chi_0 + \frac{4}{5}\bar{s}_{\rm c0}} \tag{S124}$$

by using eq. (S112).

In the $\theta$-solvent condition, the free energy per DNA unit has the form

$$\frac{f_{\rm b}(\alpha_{\rm cb}^{\rm s}, \alpha_{\rm hb}^{\rm s}, \phi_{\rm b}^{\rm s}) b^3}{\phi_{\rm b}^{\rm s} k_{\rm B} T} = \frac{1}{2}(\phi_{\rm b}^{\rm s})^2 - \frac{1}{3}\alpha_{\rm cb}^{\rm s}(0)(\phi_{\rm b}^{\rm s})^2 + \bar{f}_{\rm c0} - 1. \tag{S125}$$

The DNA volume fraction in the swollen phase is derived as

$$\phi_{\rm b}^{\rm s} = \frac{3}{5}\left(-\bar{s}_{\rm c0} + \sqrt{\bar{s}_{\rm c0}^2 + \frac{10}{3}\left(1 - \bar{f}_{\rm c0} - \chi_0 - \chi_{\rm m} + \epsilon_{\rm h} - \mu_{\rm h}\right)}\right) \tag{S126}$$

by using eq. (S112).

For both cases of the solvents, the occupancy $\alpha_{\rm c0}$ of condensin complexes must satisfy the condition

$$-\frac{\alpha_{\rm c0}}{1+n_{\rm h0}}\log\frac{\alpha_{\rm c0}}{1+n_{\rm h0}} - (\epsilon_{\rm c} - \mu_{\rm c} - 2)\frac{\alpha_{\rm c0}}{1+n_{\rm h0}} > \chi_0 + \chi_{\rm m} - (\epsilon_{\rm h} - \mu_{\rm h} + 1) \tag{S127}$$

for the DNA volume fraction $\phi_{\rm b}^{\rm s}$ at the binodal to be a positive value. We assumed $n_{\rm b} \approx 1$ to derive eq. (S127).

#### S5.3 Swollen phase in central region

In the steady state, the occupancy of condensin complexes has the form

$$\alpha_{\rm cg} = \alpha_{\rm c0}(1 - \alpha_{\rm hg})(1 - \phi_{\rm g}), \tag{S128}$$

see eq. (28) in the main article. The subscripts "cg" and "hg" indicate the quantities of condensin complexes and linker histones in the central region, respectively. Substituting eq. (S128) into eq. (S36) leads to the form

$$\log\alpha_{\rm hg} - \log(1 - \alpha_{\rm hg}) - \log\left(1 - \alpha_{\rm c0}(1 - \phi_{\rm g})\right) - 2\chi_{\rm m}\phi_{\rm g}\alpha_{\rm hg} + \epsilon_{\rm h} - \mu_{\rm h} = 0. \tag{S129}$$

By using $\alpha_{\mathrm{hg}} \ll 1$ in the swollen state, eq. (S129) has an approximate form

$$\log \alpha_{\mathrm{hg}} - \log\left(1 - \alpha_{\mathrm{c0}}(1-\phi_{\mathrm{g}})\right) + \epsilon_{\mathrm{h}} - \mu_{\mathrm{h}} = 0, \tag{S130}$$

where we expanded eq. (S129) with respect to the power series of $\alpha_{\mathrm{hg}}$ and omitted higher order terms. By solving eq. (S130) with respect to $\alpha_{\mathrm{hg}}$, the occupancy of linker histones is derived as

$$\alpha_{\mathrm{hg}} = \left(1 - \alpha_{\mathrm{c0}}(1-\phi_{\mathrm{g}})\right) \mathrm{e}^{-(\epsilon_{\mathrm{h}}-\mu_{\mathrm{h}})} \tag{S131}$$

By using $\alpha_{\mathrm{hg}} \ll 1$, the condensin occupancy $\alpha_{\mathrm{cg}}$ has an approximate form

$$\alpha_{\mathrm{cg}} = \alpha_{\mathrm{c0}}(1-\phi_{\mathrm{g}})(1 - \mathrm{e}^{-(\epsilon_{\mathrm{h}}-\mu_{\mathrm{h}})}). \tag{S132}$$

For cases in which the DNA tension is relatively small, $\zeta \ll 1$, the fraction $n_{\mathrm{e}}$ of DNA units in the elastically effective chains has an asymptotic form

$$\frac{n_{\mathrm{e}}}{1-n_{\mathrm{b}}} = \frac{1}{1 + 2\alpha_{\mathrm{cg}} v_0 \tau_{\mathrm{ex}} T(t)} + 2\alpha_{\mathrm{cg}} v_0 \tau_{\mathrm{ex}} T(t) \zeta, \tag{S133}$$

which is derived by expanding eq. (S83) in a power series of $\zeta$ and by omitting the higher order terms. In this case, the elastic stress has an asymptotic form

$$\Pi_{\mathrm{ela}} = -G_0 \frac{1 + 2\alpha_{\mathrm{cg}} v_0 \tau_{\mathrm{ex}} T(t)}{(1-n_{\mathrm{b}})^{4/3}} \left(\frac{\phi_{\mathrm{g}}}{\phi_0}\right)^{1/3}. \tag{S134}$$

Eq. (S134) implies that the loop extrusion in the central region enhances the shear modulus of the DNA network by the factor $1 + 2\alpha_{\mathrm{cg}} v_0 \tau_{\mathrm{ex}} T(t)$.

For cases in which the DNA tension is large, $\zeta \approx 1$, the fraction $n_{\mathrm{e}}$ has an asymptotic form

$$\frac{n_{\rm e}}{1-n_{\rm b}} = \sqrt{\zeta}\left[1+\frac{1}{2\alpha_{\rm cg}v_0\tau_{\rm ex}T(t)}\frac{1-\sqrt{\zeta}}{\sqrt{\zeta}}\right], \tag{S135}$$

which is derived by expanding eq. (S83) in a power series of $1/(2\alpha_{\rm c}v_0\tau_{\rm ex}T(t))$ and by omitting the higher order terms. In this case, the elastic stress has an asymptotic form

$$\Pi_{\rm ela} = -\sqrt{\frac{2}{3}\frac{G_0 f_{\rm s}}{\phi_0^{1/3}b^2}}\left(\frac{\phi_{\rm g}}{1-n_{\rm b}}\right)^{2/3}. \tag{S136}$$

#### S5.4 Osmotic pressure in central region

##### S5.4.1 Swollen phase: Good solvent

In the good solvent condition for both of the peripheral and central regions, the DNA volume fraction in the peripheral region is derived as

$$\phi_{\rm b} = \left(\frac{6\eta_{\rm m}^{4/3}}{(1+\alpha_{\rm cb}(0))^2-2\chi_0}\right)^{1/3}\left(\frac{\phi_{\rm g}}{1-n_{\rm b}}\right)^{4/9} \tag{S137}$$

by substituting eq. (S27) and $\sigma_0 b^2 = \eta_{\rm m}^{2/3}$ into eq. (S102). By using eq. (S137), the lateral osmotic pressure $\Pi_{\rm sol}(\phi_{\rm g})$ and the ratio $h/r_{\rm in}$ are derived as

$$\frac{\Pi_{\rm sol}(\phi_{\rm b})b^3}{k_{\rm B}T} = \frac{1}{2}(6\eta_{\rm m}^{4/3})^{2/3}\left((1+\alpha_{\rm cb}(0))^2-2\chi_0\right)^{1/3}\left(\frac{\phi_{\rm g}}{1-n_{\rm b}}\right)^{8/9} \tag{S138}$$

and

$$\frac{2h}{r_{\rm in}} = \frac{2}{3}n_{\rm b}\left(\frac{(1+\alpha_{\rm cb}(0))^2-2\chi_0}{6\eta_{\rm m}^{4/3}}\right)^{1/3}\left(\frac{\phi_{\rm g}}{1-n_{\rm b}}\right)^{5/9}, \tag{S139}$$

respectively, see eqs. (S50) and (S100). The osmotic pressure contribution $\Delta\Pi_{\rm g}$ by the the peripheral region is derived by using eqs. (S49) and (S51). The osmotic pressure in the central region has the form

$$\frac{\Pi_{\rm in} b^3}{k_{\rm B}T} = \frac{\Pi_{\rm ela} b^3}{k_{\rm B}T} + \frac{1}{2}\left((1+\alpha_{\rm cg}(0))^2 - 2\chi_0\right)\phi_{\rm g}^2 + \frac{1}{3}(6\eta_{\rm m}^{4/3})^{1/3} n_{\rm b}\left((1+\alpha_{\rm cb}(0))^2 - 2\chi_0\right)^{2/3}\left(\frac{\phi_{\rm g}}{1-n_{\rm b}}\right)^{13/9} \quad \text{(S140)}$$

for $2h/r_{\rm in} < 1$ and

$$\frac{\Pi_{\rm in} b^3}{k_{\rm B}T} = \frac{\Pi_{\rm ela} b^3}{k_{\rm B}T} + \frac{1}{2}\left((1+\alpha_{\rm cg}(0))^2 - 2\chi_0\right)\phi_{\rm g}^2 + \frac{1}{2}(6\eta_{\rm m}^{4/3})^{2/3}\left((1+\alpha_{\rm cb}(0))^2 - 2\chi_0\right)^{1/3}\left(\frac{\phi_{\rm g}}{1-n_{\rm b}}\right)^{8/9} \quad \text{(S141)}$$

for $2h/r_{\rm in} > 1$, see eq. (S52). The elastic stress $\Pi_{\rm ela}$ has the forms of eqs. (S134) and (S136), depending on the DNA tension.

###### S5.4.2 Swollen phase: $\theta$-solvent

In the $\theta$-solvent condition for both of the central and peripheral regions, the DNA volume fraction in the peripheral region is derived as

$$\phi_{\rm b} = (9\eta_{\rm m})^{4/3}\left(\frac{\phi_{\rm g}}{1-n_{\rm b}}\right)^{1/3} \quad \text{(S142)}$$

by substituting eq. (S27) into eq. (S103). The lateral osmotic pressure $\Pi_{\rm sol}$ and the ratio $h/r_{\rm in}$ have the forms

$$\frac{\Pi_{\rm sol}(\phi_{\rm b}) b^3}{k_{\rm B}T} = \frac{1}{3}(9\eta_{\rm m}^{4/3})^{3/4}\frac{\phi_{\rm g}}{1-n_{\rm b}} \quad \text{(S143)}$$

$$\frac{2h}{r_{\rm in}} = \frac{2}{3}\frac{n_{\rm b}}{(9\eta_{\rm m}^{4/3})^{1/4}}\left(\frac{\phi_{\rm g}}{1-n_{\rm b}}\right)^{2/3}, \quad \text{(S144)}$$

respectively, see eqs. (S50) and (S100). The osmotic pressure in the central region has the form

$$\frac{\Pi_{\rm in} b^3}{k_{\rm B}T} = \frac{\Pi_{\rm ela} b^3}{k_{\rm B}T} + \frac{1}{3}\phi_{\rm g}^3 + \frac{2}{9} n_{\rm b} (9\eta_{\rm m}^{4/3})^{1/2} \left( \frac{\phi_{\rm g}}{1 - n_{\rm b}} \right)^{5/3} \tag{S145}$$

for $2h/r_{\rm in} < 1$ and

$$\frac{\Pi_{\rm in} b^3}{k_{\rm B}T} = \frac{\Pi_{\rm ela} b^3}{k_{\rm B}T} + \frac{1}{3}\phi_{\rm g}^3 + \frac{1}{3} (9\eta_{\rm m}^{4/3})^{3/4} \left( \frac{\phi_{\rm g}}{1 - n_{\rm b}} \right) \tag{S146}$$

for $2h/r_{\rm in} > 1$.

##### S5.4.3 Condensed phase

In the condensed phase, we write the DNA volume fraction and the occupancy of linker histones in the forms

$$\phi_{\rm g} = 1 - \delta\phi_{\rm g} \tag{S147}$$

$$\alpha_{\rm hg} = 1 - \delta\alpha_{\rm hg} \tag{S148}$$

with $\delta\phi_{\rm g} \ll 1$ and $\delta\alpha_{\rm hg} \ll 1$. The occupancy of condensin complexes is the second order in $\delta\phi_{\rm g}$ and $\delta\alpha_{\rm hg}$ and thus is negligible

$$\alpha_{\rm cg} = \alpha_{\rm c0} \delta\phi_{\rm g} \delta\alpha_{\rm hg} \approx 0, \tag{S149}$$

see eq. (S128). By using eqs. (S147) - (S149), eq. (S129) is rewritten as

$$-\log \delta\alpha_{\rm hg} - 2\chi_{\rm m} + \epsilon_{\rm h} - \mu_{\rm h} = 0. \tag{S150}$$

The occupancy of linker histones is derived as

$$\alpha_{\rm hg} = 1 - {\rm e}^{-2\chi_{\rm m}+\epsilon_{\rm h}-\mu_{\rm h}} \tag{S151}$$

by solving eq. (S150) with respect to $\delta\alpha_{\rm hg}$ and by using eq. (S148).

The osmotic pressure $\Pi_{\rm sol}(\phi_{\rm g})$ has an asymptotic form

$$\frac{\Pi_{\rm sol}(\phi_{\rm g})b^3}{k_{\rm B}T} = -\log\delta\phi_{\rm g} - 1 - \chi_0 - \chi_{\rm m}, \tag{S152}$$

which is derived by substituting eqs. (S147) - (S149) into eq. (S45), by expanding it in a power series of $\delta\phi_{\rm g}$ and $\delta\alpha_{\rm hg}$, and by omitting the higher order terms. By using eq. (S149), the fraction $n_{\rm e}$ of DNA units in the elastically effective chains has the form

$$n_{\rm e} = 1 - n_{\rm b}. \tag{S153}$$

By using eqs. (S147), (S149) and (S153), the elastic stress is derived as

$$\Pi_{\rm ela} = -\frac{G_0\phi_0^{-1/3}}{(1-n_{\rm b})^{4/3}}. \tag{S154}$$

For cases that most of DNA units in the peripheral brush region have been already reeled into the central region, the osmotic pressure contribution $\Delta\Pi_{\rm g}$ is negligible. In such cases, the osmotic pressure in the central region has an approximate form

$$\frac{\Pi_{\rm in}b^3}{k_{\rm B}T} = -\frac{G_0 b^3}{\phi_0^{1/3}k_{\rm B}T}\frac{1}{(1-n_{\rm b})^{4/3}} - \log\delta\phi_{\rm g} - 1 - \chi_0 - \chi_{\rm m}. \tag{S155}$$

#### S5.5 DNA volume fraction for fast relaxation limit

In the fast relaxation limit, the DNA tension is relaxed rapidly and does not decelerate the loop extrusion, $\zeta \ll 1$. The DNA volume fraction is determined by the balance of osmotic pressure, $\Pi_{\rm in} = 0$.

##### S5.5.1 Swollen phase: Good solvent

In the good solvent condition for both of the central and peripheral regions, $(1+\alpha_{\rm c})^2 - 2\chi_0 > 0$, the pressure balance equations are written as

$$-\Lambda s^{-5/3}\left(\frac{\phi_{\rm g}}{s\phi_{\rm g}^*}\right)^{1/3} + \left(\frac{\phi_{\rm g}}{s\phi_{\rm g}^*}\right)^{2} + \left(\frac{\phi_{\rm g}}{s\phi_{\rm g}^*}\right)^{13/9} = 0 \tag{S156}$$

for $2h/r_{\rm in} < 1$ and

$$-\Lambda\left(\frac{\phi_{\rm g}}{\phi_{\rm g}^*}\right)^{1/3} + \left(\frac{\phi_{\rm g}}{\phi_{\rm g}^*}\right)^{2} + \left(\frac{\phi_{\rm g}}{\phi_{\rm g}^*}\right)^{8/9} = 0 \tag{S157}$$

for $2h/r_{\rm in} > 1$ by using

$$\phi_{\rm g}^* = \frac{(6\eta_{\rm m}^{4/3})^{3/5}}{(1-n_{\rm b})^{4/5}}\frac{((1+\alpha_{\rm cb}(0))^2 - 2\chi_0)^{3/10}}{((1+\alpha_{\rm cg}(0))^2 - 2\chi_0)^{9/10}} \tag{S158}$$

$$\Lambda = \frac{1}{3}\frac{G_0 b^3(1+2\alpha_{\rm cg}v_0\tau_{\rm ex}T(t))}{\phi_0^{1/3}k_{\rm B}T\eta_{\rm m}^{4/3}}\left(\frac{(1+\alpha_{\rm cg}(0))^2 - 2\chi_0}{(1+\alpha_{\rm cb}(0))^2 - 2\chi_0}\right)^{1/2} \tag{S159}$$

$$s = \left(\frac{2}{3}\frac{n_{\rm b}}{1-n_{\rm b}}\left(\frac{(1+\alpha_{\rm cb}(0))^2 - 2\chi_0}{(1+\alpha_{\rm cg}(0))^2 - 2\chi_0}\right)^{1/2}\right)^{9/5}, \tag{S160}$$

see eqs. (S140) and (S141). The ratio $2h/r_{\rm in}$ is also rewritten as

$$\frac{2h}{r_{\rm in}} = s^{5/9}\left(\frac{\phi_{\rm g}}{\phi_{\rm g}^*}\right)^{5/9} \tag{S161}$$

Eqs. (S156) and (S157) have three types of solutions depending on the two parameters, $\Lambda$ and $s$, see fig. S6**a**:

i). For the case of $s < \Lambda^{3/5}$ or $\Lambda > 1$, the DNA volume fraction in the central region has an asymptotic form

$$\begin{aligned}\phi_{\mathrm{g}} &= \Lambda^{3/5}\phi_{\mathrm{g}}^{*}\\ &= \left(\frac{2G_0b^3(1+2\alpha_{\mathrm{cg}}v_0\tau_{\mathrm{ex}}T(t))}{\phi_0^{1/3}k_{\mathrm{B}}T((1+\alpha_{\mathrm{cb}}(0))^2-2\chi_0)}\right)^{3/5}\frac{1}{(1-n_{\mathrm{b}})^{4/5}}.\end{aligned} \tag{S162}$$

In this case, the osmotic pressure $\Pi_{\mathrm{sol}}(\phi_{\mathrm{g}})$ in the central region is larger than the osmotic pressure contribution $\Delta\Pi_{\mathrm{g}}$ of the peripheral region and is balanced by the elastic stress. The DNA volume fraction in the peripheral region is derived as

$$\phi_{\mathrm{b}} = \left(\frac{2G_0b^3(1+2\alpha_{\mathrm{cg}}v_0\tau_{\mathrm{ex}}T(t))}{\phi_0^{1/3}k_{\mathrm{B}}T((1+\alpha_{\mathrm{cb}}(0))^2-2\chi_0)}\right)^{4/15}\frac{6^{1/3}\eta_{\mathrm{m}}^{4/9}}{((1+\alpha_{\mathrm{cb}}(0))^2-2\chi_0)^{1/3}}\frac{1}{(1-n_{\mathrm{b}})^{4/5}} \tag{S163}$$

by substituting eq. (S162) into eq. (S137). The ratio of the DNA volume fractions thus has the form

$$\frac{\phi_{\mathrm{g}}}{\phi_{\mathrm{b}}} = \left(\frac{G_0b^3(1+2\alpha_{\mathrm{cg}}v_0\tau_{\mathrm{ex}}T(t))}{3\eta_{\mathrm{m}}^{4/3}\phi_0^{1/3}k_{\mathrm{B}}T}\right)^{1/3}\left(\frac{(1+\alpha_{\mathrm{cb}}(0))^2-2\chi_0}{(1+\alpha_{\mathrm{cg}}(0))^2-2\chi_0}\right)^{1/3}. \tag{S164}$$

ii) For the case of $\Lambda^{3/5} < s < \Lambda^{-9/5}$ and $\Lambda < 1$, the DNA volume fraction in the central region has an asymptotic form

$$\begin{aligned}\phi_{\mathrm{g}} &= s^{-1/2}\Lambda^{9/10}\phi_{\mathrm{g}}^{*}\\ &= \left(\frac{3G_0b^3(1+2\alpha_{\mathrm{cg}}v_0\tau_{\mathrm{ex}}T(t))}{6^{1/3}\eta_{\mathrm{m}}^{4/9}\phi_0^{1/3}k_{\mathrm{B}}T((1+\alpha_{\mathrm{cb}}(0))^2-2\chi_0)^{2/3}}\right)^{9/10}\frac{(1-n_{\mathrm{b}})^{1/10}}{n_{\mathrm{b}}^{9/10}}.\end{aligned} \tag{S165}$$

In this case, the osmotic pressure contribution $\Delta\Pi_{\mathrm{g}}$ in the peripheral region is larger than the osmotic pressure $\Pi_{\mathrm{sol}}(\phi_{\mathrm{g}})$ in the central region and is balanced by the elastic stress $\Pi_{\mathrm{ela}}$. The ratio $2h/r_{\mathrm{in}}$ is smaller than unity. The DNA volume fraction in the peripheral region has the form

$$\phi_{\rm b} \;=\; \left(\frac{3G_0b^3(1+2\alpha_{\rm cg}v_0\tau_{\rm ex}T(t))}{6^{1/3}\eta_{\rm m}^{4/9}\phi_0^{1/3}k_{\rm B}T}\right)^{2/5}\frac{6^{1/3}\eta_{\rm m}^{4/9}}{((1+\alpha_{\rm cb}(0))^2-2\chi_0)^{3/5}}\frac{1}{n_{\rm b}^{2/5}(1-n_{\rm b})^{2/5}}. \tag{S166}$$

The ratio of the DNA volume fractions has the form

$$\frac{\phi_{\rm g}}{\phi_{\rm b}} = \left(\frac{G_0b^3(1+2\alpha_{\rm cg}v_0\tau_{\rm ex}T(t))}{2\eta_{\rm m}^{4/3}\phi_0^{1/3}k_{\rm B}T}\right)^{1/2}\left(\frac{1-n_{\rm b}}{n_{\rm b}}\right)^{1/2}. \tag{S167}$$

iii) For the case of $s > \Lambda^{-9/5}$ and $\Lambda < 1$, the DNA volume fraction in the central region has an asymptotic form

$$\begin{aligned}\phi_{\rm g} &= \Lambda^{9/5}\phi_{\rm g}^{*}\\ &= \left(\frac{2G_0b^3(1+2\alpha_{\rm cg}v_0\tau_{\rm ex}T(t))}{6^{2/3}\eta_{\rm m}^{8/9}\phi_0^{1/3}k_{\rm B}T((1+\alpha_{\rm cb}(0))^2-2\chi_0)}\right)^{9/5}\frac{1}{(1-n_{\rm b})^{4/5}}.\end{aligned} \tag{S168}$$

In this case, the osmotic pressure contribution $\Delta\Pi_{\rm g}$ in the peripheral region is larger than the osmotic pressure $\Pi_{\rm sol}(\phi_{\rm g})$ in the central region and is balanced by the elastic stress $\Pi_{\rm ela}$. The ratio $2h/r_{\rm in}$ is larger than unity. The DNA volume fraction in the peripheral region has the form

$$\phi_{\rm b} \;=\; \left(\frac{2G_0b^3(1+2\alpha_{\rm cg}v_0\tau_{\rm ex}T(t))}{6^{2/3}\eta_{\rm m}^{8/9}\phi_0^{1/3}k_{\rm B}T((1+\alpha_{\rm cb}(0))^2-2\chi_0)}\right)^{4/5}\frac{6^{1/3}\eta_{\rm m}^{4/9}}{((1+\alpha_{\rm cb}(0))^2-2\chi_0)^{1/3}}\frac{1}{(1-n_{\rm b})^{4/5}}. \tag{S169}$$

The ratio of the DNA volume fractions has the form

$$\frac{\phi_{\rm g}}{\phi_{\rm b}} = \frac{G_0b^3(1+2\alpha_{\rm cg}v_0\tau_{\rm ex}T(t))}{3\eta_{\rm m}^{4/3}\phi_0^{1/3}k_{\rm B}T((1+\alpha_{\rm cb}(0))^2-2\chi_0)^{2/3}}. \tag{S170}$$

##### S5.5.2 Swollen phase: $\theta$-solvent

In the $\theta$-solvent condition for both of the central and peripheral regions, $(1+\alpha_{\rm c})^2 - 2\chi_0 \approx 0$, the pressure balance equations are written as

$$-\Lambda_\theta s_\theta^{-8/3} \left(\frac{\phi_{\rm g}}{s_\theta \phi_\theta^*}\right)^{1/3} + \left(\frac{\phi_{\rm g}}{s_\theta \phi_\theta^*}\right)^{3} + \left(\frac{\phi_{\rm g}}{s_\theta \phi_\theta^*}\right)^{5/3} = 0 \tag{S171}$$

for $2h/r_{\rm in} < 1$ and

$$-\Lambda_\theta \left(\frac{\phi_{\rm g}}{s_\theta \phi_\theta^*}\right)^{1/3} + \left(\frac{\phi_{\rm g}}{s_\theta \phi_\theta^*}\right)^{3} + \left(\frac{\phi_{\rm g}}{s_\theta \phi_\theta^*}\right) = 0 \tag{S172}$$

for $2h/r_{\rm in} > 1$ by using

$$\phi_\theta^* = \frac{(9\eta_{\rm m}^{4/3})^{3/8}}{(1-n_{\rm b})^{1/2}} \tag{S173}$$

$$\Lambda_\theta = \frac{G_0 b^3 (1 + 2\alpha_{\rm cg} v_0 \tau_{\rm ex} T(t))}{3\eta_{\rm m}^{4/3} \phi_0^{1/3} k_{\rm B} T} \tag{S174}$$

$$s_\theta = \left(\frac{2}{3}\frac{n_{\rm b}}{1-n_{\rm b}}\right)^{3/4}, \tag{S175}$$

see eqs. (S145) and (S146). Eqs. (S171) and (S172) have three types of solutions depending on the two parameters, $\Lambda_\theta$ and $s_\theta$, see fig. S6**b**:

i) For the case of $s_\theta < \Lambda_\theta^{3/8}$ or $\Lambda_\theta > 1$, the DNA volume fraction in the central region has an asymptotic form

$$\begin{aligned}\phi_{\rm g} &= \Lambda_\theta^{3/8} \phi_\theta^* \\ &= \left(\frac{3 G_0 b^3 (1 + 2\alpha_{\rm cg} v_0 \tau_{\rm ex} T(t))}{\phi_0^{1/3} k_{\rm B} T}\right)^{3/8} \frac{1}{(1-n_{\rm b})^{1/2}}.\end{aligned} \tag{S176}$$

In this case, the osmotic pressure $\Pi_{\rm sol}(\phi_{\rm g})$ of the central region is larger than the pressure contribution $\Delta\Pi_{\rm g}$ of the peripheral region and is balanced by the elastic stress

$\Pi_{\rm ela}$. The DNA volume fraction in the peripheral region is derived as

$$\phi_{\rm b} = (9\eta_{\rm m}^{4/3})^{1/4}\left(\frac{3G_0b^3(1+2\alpha_{\rm cg}v_0\tau_{\rm ex}T(t))}{\phi_0^{1/3}k_{\rm B}T}\right)^{1/8}\frac{1}{(1-n_{\rm b})^{1/2}} \tag{S177}$$

by substituting eq. (S176) into eq. (S142). The ratio of the DNA volume fractions thus has the form

$$\frac{\phi_{\rm g}}{\phi_{\rm b}} = \left(\frac{G_0b^3(1+2\alpha_{\rm cg}v_0\tau_{\rm ex}T(t))}{3\eta_{\rm m}^{4/3}\phi_0^{1/3}k_{\rm B}T}\right)^{1/4}. \tag{S178}$$

ii) For the case of $\Lambda_\theta^{3/8} < s_\theta < \Lambda_\theta^{-3/4}$ and $\Lambda_\theta < 1$, the DNA volume fraction in the central region has an asymptotic form

$$\begin{aligned}\phi_{\rm g} &= s^{-1}\Lambda_\theta^{3/4}\phi_\theta^* \\ &= \left(\frac{3G_0b^3(1+2\alpha_{\rm cg}v_0\tau_{\rm ex}T(t))}{2\eta_{\rm m}^{2/3}}\right)^{3/4}\frac{(1-n_{\rm b})^{1/4}}{n_{\rm b}^{3/4}}.\end{aligned} \tag{S179}$$

The DNA volume fraction in the peripheral region has the form

$$\phi_{\rm b} = (9\eta_{\rm m}^{4/3})^{1/4}\left(\frac{3G_0b^3(1+2\alpha_{\rm cg}v_0\tau_{\rm ex}T(t))}{2\eta_{\rm m}^{2/3}}\right)^{1/4}\frac{1}{n_{\rm b}^{1/4}(1-n_{\rm b})^{1/4}}. \tag{S180}$$

The ratio of the DNA volume fractions has the form

$$\frac{\phi_{\rm g}}{\phi_{\rm b}} = \left(\frac{G_0b^3(1+2\alpha_{\rm cg}v_0\tau_{\rm ex}T(t))}{2\eta_{\rm m}^{4/3}\phi_0^{1/3}k_{\rm B}T}\right)^{1/2}\left(\frac{1-n_{\rm b}}{n_{\rm b}}\right)^{1/2}. \tag{S181}$$

iii) For the case of $s_\theta > \Lambda_\theta^{-3/4}$ and $\Lambda_\theta < 1$, the DNA volume fraction in the central region has an asymptotic form

$$\phi_{\rm g} = \Lambda_\theta^{3/2}\phi_\theta^* = \left(\frac{G_0 b^3(1+2\alpha_{\rm cg}v_0\tau_{\rm ex}T(t))}{\sqrt{3}\eta_{\rm m}\phi_0^{1/3}k_{\rm B}T}\right)^{3/2}\frac{1}{(1-n_{\rm b})^{1/2}}. \quad \text{(S182)}$$

The DNA volume fraction in the peripheral region has the form

$$\phi_{\rm b} = (9\eta_{\rm m}^{4/3})^{1/4}\left(\frac{G_0 b^3(1+2\alpha_{\rm cg}v_0\tau_{\rm ex}T(t))}{\sqrt{3}\eta_{\rm m}\phi_0^{1/3}k_{\rm B}T}\right)^{1/2}\frac{1}{(1-n_{\rm b})^{1/2}}. \quad \text{(S183)}$$

The ratio of the DNA volume fractions has the form

$$\frac{\phi_{\rm g}}{\phi_{\rm b}} = \frac{G_0 b^3(1+2\alpha_{\rm cg}v_0\tau_{\rm ex}T(t))}{3\eta_{\rm m}^{4/3}\phi_0^{1/3}k_{\rm B}T}. \quad \text{(S184)}$$

##### S5.5.3 Condensed phase

By solving the pressure balance equation, $\Pi_{\rm in} = 0$, the DNA volume fraction in the central region is derived as

$$\phi_{\rm g} = 1 - {\rm e}^{-\frac{G_0 b^3}{\phi_0^{1/3}k_{\rm B}T}\frac{1}{(1-n_{\rm b})^{4/3}} - 1 - \chi_0 - \chi_{\rm m}}, \quad \text{(S185)}$$

see eqs. (S147) and (S155).

#### S5.6 DNA tension in condensed phase

In the condensed phase the contribution of DNA tension from the central region has an asymptotic form

$$\mu_{\rm ex}^{\rm g} = \frac{3}{2}\frac{G_0 b^3}{\phi_0^{1/3}k_{\rm B}T}\frac{1}{(1-n_{\rm b})^{4/3}} - \log\delta\phi_{\rm g} - 1 - 2\chi_0 - 2\chi_{\rm m} + \epsilon_{\rm h} - \mu_{\rm h}, \quad \text{(S186)}$$

which is derived by substituting eqs. (S147) - (S149) into eq. (S61), by expanding it in a power series of $\delta\phi_{\rm g}$ and $\delta\alpha_{\rm hg}$, and by omitting the higher order terms. For cases in which the central region is rapidly relaxed, $\Pi_{\rm in} = 0$, eq. (S186) is rewritten as

$$\mu_{\rm ex}^{\rm g} = \frac{5}{2}\frac{G_0 b^3}{\phi_0^{1/3} k_{\rm B} T}\frac{1}{(1-n_{\rm b})^{4/3}} - \chi_0 - \chi_{\rm m} + \epsilon_{\rm h} - \mu_{\rm h}, \tag{S187}$$

see eq. (S185). The contribution of the entropic elasticity of DNA, the first term of eq. (S187), is relatively weak because the DNA network is shranken and is lack of elastically ineffective chains in the condensed phase, see eqs. (S147) and (S149). Moreover, after most of the DNA units in the peripheral region are reeled into the central region, the fraction of DNA in the peripheral region is relatively small, $n_{\rm b} \ll 1$. The main contribution to the DNA tension is thus the histone-histone interaction, $-\chi_{\rm m} + \epsilon_{\rm h} - \mu_{\rm h}$. The magnitude of the histone-histone interaction is expected to be in the order of the thermal energy and thus is much smaller than the stall force of condensin complexes.

#### S5.7 Loop extrusion dynamics for fast relaxation limit

##### S5.7.1 Large condensin concentration

The time evolution equation of the number $N_{\rm bl}$ of DNA units in a loop at the peripheral region has the forms of eqs. (S70) and (S71). In the fast relaxation limit, the DNA tension is relaxed rapidly and does not decelerate the loop extrusion, $v_{\rm ex} \approx v_0$. When the occupancy $\alpha_{\rm c0}$ of condensin complexes is large enough, DNA reeling due to condensin unloading, the second term of eq. (S71), is also negligible. The fraction $n_{\rm b}$ of DNA units in the peripheral region thus has an asymptotic form

$$N_{\rm b} \approx 2 n_{\rm l} v_0 t. \tag{S188}$$

##### S5.7.2 Small condensin concentration

The stable and uniform peripheral region is assembled when the occupancy $\alpha_{c0}$ of condensin complexes is relatively small. In such case, DNA reeling by condensin unloading, the second term of eq. (S71), is significant. The peripheral region is at the $\theta$-solvent condition for $\chi_0 \approx 1/2$ because of the lack of condensin occupancy. The time evolution equation of the DNA units in the peripheral region has the form

$$\frac{dN_\mathrm{b}}{dt} = 2v_0 n_\mathrm{l} \left(1 - \frac{1}{\mathrm{e}^{2v_0\tau_\mathrm{ex}\alpha_\mathrm{cb}} - 1}\right), \tag{S189}$$

where we used $v_\mathrm{ex} \approx v_0$ effective in the fast relaxation limit. The steady state condition, $dN_\mathrm{b}/dt = 0$, is rewritten in the form

$$\frac{n_\mathrm{b}}{n_\mathrm{b} + n_\mathrm{h0}} \left(1 - \frac{\phi_\mathrm{b0}}{(1 - n_\mathrm{b})^{1/2}}\right) = \frac{\log 2}{2v_0\tau_\mathrm{ex}\alpha_\mathrm{c0}(1 - \mathrm{e}^{-(\epsilon_\mathrm{h} - \mu_\mathrm{h})})}, \tag{S190}$$

where we used eq. (S95) and the fact that the DNA volume fraction $\phi_\mathrm{b}$ has the form

$$\phi_\mathrm{b} = \frac{\phi_\mathrm{b0}}{(1 - n_\mathrm{b})^{1/2}} \tag{S191}$$

in the $\theta$-solvent condition for both $\Lambda_\theta < 1$ and $\Lambda_\theta > 1$, see eqs. (S176) and (S182).

The left side of eq. (S190) is a non-monotonic function and there is not solution if the right side of this equation is too large. The derivative of the left side of eq. (S190) has the form

$$\frac{\partial}{\partial n_\mathrm{b}} \left[\frac{n_\mathrm{b}}{n_\mathrm{b} + n_\mathrm{h0}} \left(1 - \frac{\phi_\mathrm{b0}}{(1 - n_\mathrm{b})^{1/2}}\right)\right] = \frac{n_\mathrm{h0}}{(n_\mathrm{b} + n_\mathrm{h0})^2} \left(1 - \frac{\phi_\mathrm{b0}}{(1 - n_\mathrm{b})^{1/2}}\right) - \frac{1}{2} \frac{n_\mathrm{b}}{n_\mathrm{b} + n_\mathrm{h0}} \frac{\phi_\mathrm{b0}}{(1 - n_\mathrm{b})^{3/2}}. \tag{S192}$$

For cases in which the maximum of the left side of eq. (S190) is at $n_\mathrm{b} \approx 1$, the DNA fraction

$n_{\rm b}$ at the maximum has an approximate form

$$n_{\rm b}^{\rm th} = 1 - \left(\frac{1+n_{\rm h0}}{2n_{\rm h0}}\right)^{2/3}\phi_{\rm b0}^{2/3}. \tag{S193}$$

Eq. (S193) is thus effective for $\phi_{\rm b0} \ll 1$. Eq. (S190) thus has a solution the case of $\alpha_{\rm c0} \geq \alpha_{\rm c0}^{\rm th}$ with

$$\alpha_{\rm c0}^{\rm th} = \frac{(1+n_{\rm h0})\log 2}{2v_0\tau_{\rm ex}(1-{\rm e}^{-(\epsilon_{\rm h}-\mu_{\rm h})})}\frac{1}{1-\frac{3}{2}\left(\frac{2n_{\rm h0}}{1+n_{\rm h0}}\right)^{1/3}\phi_{\rm b0}^{2/3}}. \tag{S194}$$

Eq. (S194) is derived by substituting eq. (S194) into eq. (S190).

The DNA fraction $n_{\rm b}$ at the steady state has an approximate form

$$n_{\rm b}^{\rm s} = 1 - \frac{\alpha_{\rm c0}^2}{(\alpha_{\rm c0}-\bar{\alpha}_{\rm c0})^2}\phi_{\rm b0}^2 \tag{S195}$$

with

$$\bar{\alpha}_{\rm c0} = \frac{(1+n_{\rm h0})\log 2}{2v_0\tau_{\rm ex}(1-{\rm e}^{-(\epsilon_{\rm h}-\mu_{\rm h})})}. \tag{S196}$$

Eq. (S195) is derived by solving eq. (S190) for $n_{\rm b} \approx 1$.

#### S5.8 Loop extrusion dynamics for slow relaxation limit

In the slow relaxation limit, the volume of the central region is constant except for the volume of DNA extruded to the peripheral region $b^3 N_{\rm b}$. The DNA volume fraction in the central region thus has the forms

$$\begin{aligned}\phi_{\rm g} &= \frac{(1-n_{\rm b})\phi_{\rm g0}}{1-n_{\rm b}\phi_{\rm g0}} \\ &\simeq (1-n_{\rm b})\phi_{\rm g0}, \end{aligned} \tag{S197}$$

where $\phi_{g0}$ is the DNA volume fraction in the central region at the initial condition. In this case, the tension factor $\zeta$ is represented as

$$\zeta = \frac{\zeta_0}{(1-n_b)^2} \tag{S198}$$

with

$$\zeta_0 = \frac{3}{2}\frac{G_0 b^2}{\phi_0^{1/3} f_s}\phi_{g0}^{-2/3}. \tag{S199}$$

In the short time scale, the DNA tension in the central region is not yet significant, $\zeta \ll 1$. In the good solvent condition for both of the central and brush regions, $(1+\alpha_c)^2 - 2\chi_0 > 0$, the contributions of the DNA tension has approximate forms

$$\begin{aligned}\mu_{ex}^{g} &= \frac{3}{2}\frac{G_0 b^2}{\phi_0^{1/3} f_s}\phi_{g0}^{-2/3}\frac{(1+2\alpha_{cg}(0)v_0\tau_{ex}T(t))^2}{(1-n_b)^2} - 1 + \bar{f}(\alpha_{cg},\alpha_{hg}) \\ &\quad +((1+\alpha_{cg}(0))^2 - 2\chi_0)(1-n_b)\phi_{g0} \qquad \text{(S200)} \\ \mu_{ex}^{b} &= \frac{3}{4}(6\eta_m^{4/3})^{1/3}((1+\alpha_{cg}(0))^2-2\chi_0)^{2/3}\phi_{g0}^{4/9} - 1 + \bar{f}(\alpha_{cb},\alpha_{hb}) \qquad \text{(S201)}\end{aligned}$$

with

$$\begin{aligned}\bar{f}(\alpha_c,\alpha_h) &= \alpha_c\log\alpha_c + \alpha_h\log\alpha_h + (1-\alpha_c-\alpha_h)\log(1-\alpha_c-\alpha_h) \\ &\quad +(\epsilon_c-\mu_c-1)\alpha_c + (\epsilon_h-\mu_h)\alpha_h. \qquad \text{(S202)}\end{aligned}$$

Eqs. (S200) and (S201) are derived by expanding eqs. (S61) and (S62) in a power series of $\phi_g$ and by omitting the higher order terms. We used eq. (S133) for the DNA fraction in the elastically effective chains. The contributions, $\bar{f}(\alpha_{cg},\alpha_{hg})$ and $\bar{f}(\alpha_{cb},\alpha_{hb})$, of the binding free energy are in the order of unity and are much smaller than the energy scale $f_s b$ of the stall force of condensin complexes. The last term of eq. (S200) and the first term of eq. (S201) is small for $\phi_{g0} < 1$. By neglecting these small contributions, the time evolution equation of the fraction $n_{\rm b}$ of DNA units in the peripheral region is written as

$$\frac{dn_{\rm b}}{dt} = \frac{n_{\rm h0}}{\tau_{\rm ex}}\left(1 - \zeta_0\frac{(1+2\alpha_{\rm cg}(0)v_0\tau_{\rm ex}T(t))^2}{(1-n_{\rm b})^2}\right). \tag{S203}$$

Eq. (S203) is effective for cases in which the first term of this equation is larger than the second term and is negligible for $\zeta_0 \ll 1$ and $t < \tau_{\rm ex}$. The DNA fraction $n_{\rm b}$ in the peripheral region has an asymptotic form

$$n_{\rm b}(t) = n_{\rm h0}\frac{t}{\tau_{\rm ex}}. \tag{S204}$$

In the long time scale, the DNA tension is significant, $\zeta \approx 1$. In this case, the contribution of the DNA tension from the central region have an asymptotic form

$$\mu_{\rm ex}^{\rm g} = \frac{f_{\rm s}b}{k_{\rm B}T}\frac{1}{\left(1+\frac{1}{4\alpha_{\rm cg}v_0\tau_{\rm ex}}\frac{1-\sqrt{\zeta}}{\sqrt{\zeta}}\right)^2}, \tag{S205}$$

where it is derived by substituting eq. (S135) into eq. (S200) and by omitting the contributions in the order of unity. The contribution $\mu_{\rm ex}^{\rm b}$ of the DNA tension from the peripheral region is negligible because it is also in the order of unity. The time evolution equation of the DNA fraction $n_{\rm b}$ has an asympotic form

$$\begin{aligned}\frac{dn_{\rm b}}{dt} &= \frac{n_{\rm h0}}{\tau_{\rm ex}}\left(1 - \frac{1}{\left(1+\frac{1-n_{\rm b}-\zeta_0^{1/2}}{4\alpha_{\rm cg}(0)v_0\tau_{\rm ex}\zeta_0^{1/2}}\right)^2}\right)\\ &\simeq \frac{n_{\rm h0}}{\tau_{\rm ex}}\frac{1-n_{\rm b}-\zeta_0^{1/2}}{2\alpha_{\rm cg}(0)v_0\tau_{\rm ex}\zeta_0^{1/2}}\end{aligned} \tag{S206}$$

Eq. (S206) is derived by substituting eq. (S205) and $\mu_{\rm ex}^{\rm b} \approx 0$ into eq. (S71), where the second term of eq. (S71) is negligible for large condensin occupancy $\alpha_{\rm c0}$. The last form of eq. (S206) is derived by expanding the second term of eq. (S206) for $2\alpha_{\rm cg}v_0\tau_{\rm ex}\zeta_0^{1/2} \gg 1$.

By solving eq. (S206), the DNA fraction $n_\mathrm{b}$ is derived as

$$n_\mathrm{b}(t) = 1 - \zeta_0^{1/2} - (1 - \zeta_0^{1/2} - n_\mathrm{b}(t_0))\mathrm{e}^{-(t-t_0)/\tau_\mathrm{s}}. \tag{S207}$$

with

$$\frac{\tau_\mathrm{s}}{\tau_\mathrm{ex}} = \frac{2\alpha_\mathrm{cg}(0)v_0\tau_\mathrm{ex}\zeta_0^{1/2}}{n_\mathrm{h0}} = \frac{\alpha_\mathrm{cg}(0)N}{n_\mathrm{l}}\left(\frac{3}{2}\frac{G_0 b^2}{\phi_0^{1/3} f_\mathrm{s}}\right)^{1/2}\phi_\mathrm{g0}^{-1/3}. \tag{S208}$$

The short and long time regimes crossover at a time $t_0$. Eq. (S208) implies that the time scale $\tau_\mathrm{s}$ to reach the steady state increases linearly with the swelling ratio of the entangled DNA strands in the initial state. The DNA fraction $n_\mathrm{b}$ at the steady state has the form

$$n_\mathrm{b}^\mathrm{s} = 1 - \zeta_0^{1/2} = 1 - \left(\frac{3}{2}\frac{G_0 b^2}{\phi_0^{1/3} f_\mathrm{s}}\right)^{1/2}\phi_\mathrm{g0}^{-1/3}, \tag{S209}$$

see eq. (S207). The DNA volume fraction in the central region at the steady state has the form

$$\phi_\mathrm{g}^\mathrm{s} = \left(\frac{3}{2}\frac{G_0 b^2}{\phi_0^{1/3} f_\mathrm{s}}\right)^{1/2}\phi_\mathrm{g0}^{2/3}, \tag{S210}$$

where it is derived by substituting eq. (S209) into eq. (S197).

#### S5.9 Relaxation dynamics for slow relaxation limit

In the slow relaxation limit, the loop extrusion dynamics reaches the steady state, $\frac{dn_\mathrm{b}}{dt} = 0$, before the relaxation starts. In this state, the loop extrusion is stalled by the DNA tension. The fraction $n_\mathrm{b}$ of DNA units in the peripheral region is derived as

$$n_\mathrm{b} = 1 - \left(\frac{3}{2}\frac{G_0 b^2}{\phi_0^{1/3} f_\mathrm{s}}\right)^{3/4}\phi_\mathrm{g}^{-1/2} \tag{S211}$$

by using $\zeta \approx 1$. The elastic stress due to the DNA tension has the form

$$\frac{\Pi_{\rm ela} b^3}{k_{\rm B}T} = -\frac{2}{3}\frac{f_{\rm s} b}{k_{\rm B}T}\phi_{\rm g}, \tag{S212}$$

which is derived by substituting eq. (S211) into eq. (S136). The elastic stress dominates the other contributions of the osmotic pressure.

In the good solvent condition, the DNA volume fraction at the peripheral region is derived as

$$\phi_{\rm b} = \left(\frac{4\eta_{\rm m}^{4/3}}{(1+\alpha_{\rm cb}(0))^2 - 2\chi_0}\frac{\phi_0^{1/3} f_{\rm s}}{G_0 b^2}\right)^{1/3} \phi_{\rm g}^{2/3} \tag{S213}$$

by substituting eq. (S211) into eq. (S137). The thickness $h$ of the peripheral region thus has the form

$$\begin{aligned} \frac{h}{r_0} &= \frac{1}{3}\frac{n_{\rm b}}{\phi_{\rm b}}\left(\frac{\phi_{\rm g}}{1-n_{\rm b}}\right)^{2/3} \\ &\simeq \frac{1}{3}\left(\frac{2}{3}\frac{\phi_0^{1/3} f_{\rm s}}{G_0 b^2}\right)^{1/6}\left(\frac{(1+\alpha_{\rm cb}(0)^2 - 2\chi_0}{6\eta_{\rm m}^{4/3}}\right)^{1/3} \phi_{\rm g}^{1/3}. \end{aligned} \tag{S214}$$

Similarly, the surface density $\sigma$ of DNA loops in the peripheral region is derived as

$$\sigma b^2 = \sqrt{\frac{2}{3}\frac{\phi_0^{1/3} f_{\rm s}}{G_0 b^2}\eta_{\rm m}^{4/3}\phi_{\rm g}} \tag{S215}$$

by substituting eq. (S211) into eq. (S27).

The radius $r_{\rm in}$ of the central region is derived as

$$\begin{aligned} \frac{r_{\rm in}}{r_0} &= \left(\frac{1-n_{\rm b}}{\phi_{\rm g}}\right)^{1/3} \\ &= \left(\frac{3}{2}\frac{G_0 b^2}{\phi_0^{1/3} f_{\rm s}}\right)^{1/4} \phi_{\rm g}^{-1/2}, \end{aligned} \tag{S216}$$

where the last form is derived by using eq. (S211). By taking the time derivative to both sides of eq. (S216) leads to the form

$$\frac{dr_{\rm in}}{dt} = -\frac{1}{2}\left(\frac{3}{2}\frac{G_0 b^2}{\phi_0^{1/3} f_{\rm s}}\right)^{1/4} \phi_{\rm g}^{-3/2} r_0 \frac{d\phi_{\rm g}}{dt}, \tag{S217}$$

which corresponds to the left side of eq. (S86). The right side of eq. (S86) has the form

$$-\kappa(\phi_{\rm b})\frac{p_{\rm in}}{h} = -\frac{2r_0}{\tau_{\rm r}}\frac{f_{\rm s} b}{\eta_{\rm m}^{4/3} k_{\rm B} T}\left(\frac{3}{2}\frac{G_0 b^2}{\phi_0^{1/3} f_{\rm s}}\right)^{2/3}\left(\frac{6\eta_{\rm m}^{4/3}}{(1+\alpha_{\rm cb}(0))^2 - 2\chi_0}\right)^{1/3} \phi_{\rm g}^{-1/2}(1-\phi_{\rm b})^2 \tag{S218}$$

by using eqs. (S87), (S93), (S212), (S214), and (S215) with $p_{\rm in} = -\Pi_{\rm g}$. By equating eqs. (S217) and (S218), the time evolution equation of the DNA volume fraction is derived as

$$\tau_{\rm r}\frac{d\phi_{\rm g}}{dt} = 4\left(\frac{3}{2}\frac{G_0 b^2}{\phi_0^{1/3} f_{\rm s}}\right)^{5/12}\left(\frac{6\eta_{\rm m}^{4/3}}{(1+\alpha_{\rm cb}(0))^2 - 2\chi_0}\right)^{1/3}\frac{f_{\rm s} b}{\eta_{\rm m}^{4/3} k_{\rm B} T}\phi_{\rm g}^{7/6}(1-\phi_{\rm b})^2. \tag{S219}$$

Eq. (S219) is rewritten as

$$\frac{d\phi_{\rm b}}{dt} = \frac{\phi_{\rm b}^{5/4}(1-\phi_{\rm b})^2}{\tilde{\tau}_{\rm r}} \tag{S220}$$

with

$$\tilde{\tau}_{\rm r} = \frac{3}{8}\tau_{\rm r}\left(\frac{2}{3}\frac{\phi_0^{1/3} f_{\rm s}}{G_0 b^2}\right)^{1/2}\left(\frac{(1+\alpha_{\rm cb}(0))^2 - 2\chi_0}{6\eta_{\rm m}^{4/3}}\right)^{1/4}\frac{\eta_{\rm m}^{4/3} k_{\rm B} T}{f_{\rm s} b} \tag{S221}$$

by using eq. (S213). For the case of $\phi_{\rm b} < 1$, the DNA volume fraction $\phi_{\rm b}$ has an asymptotic form

$$\phi_{\rm b}(t) = \frac{\phi_{\rm b}(t_0)}{\left(1 - \frac{3}{4}\phi_{\rm b}^{1/4}(t_0)\frac{t-t_0}{\tilde{\tau}_{\rm r}}\right)^4}. \tag{S222}$$

The DNA volume fraction $\phi_{\mathrm{b}}(t)$ in the peripheral region increases with time during the relaxation process.

#### S5.10 Relaxation time per unit extrusion step

In the good solvent condition, the derivative of the osmotic pressure with respect to $n_{\mathrm{b}}$ is calculated as

$$\frac{\partial}{\partial n_{\mathrm{b}}}\left(\frac{\Pi_{\mathrm{in}} b^3}{k_{\mathrm{B}}T}\right)\Big|_{V_{\mathrm{in}}} = -\frac{G_0 b^3}{\phi_0^{1/3} k_{\mathrm{B}}T}\frac{1+2\alpha_{\mathrm{cg}} v_0 \tau_{\mathrm{ex}} T(t)}{(1-n_{\mathrm{b}})^2}\left(\frac{\phi_{\mathrm{g}}}{1-n_{\mathrm{b}}}\right)^{1/3} -((1+\alpha_{\mathrm{cg}}(0))^2-2\chi_0)(1-n_{\mathrm{b}})\left(\frac{\phi_{\mathrm{g}}}{1-n_{\mathrm{b}}}\right)^2, \qquad \text{(S223)}$$

where $|_{V_{\mathrm{in}}}$ indicates the condition of fixed volume $V_{\mathrm{in}}$, see eqs. (S140) and (S141). We used the fact that the changes of the occupancy of condensin complexes during a unit extrusion step is small, $\frac{\partial \alpha_{\mathrm{cb}}(0)}{\partial n_{\mathrm{b}}} \approx 0$. It is useful to notice the fact that $\phi_{\mathrm{g}}/(1-n_{\mathrm{b}})$ is a constant if the volume $V_{\mathrm{in}}$ of the central region is fixed in the derivation of eq. (S223). The derivative of the radius $r_{\mathrm{in}}$ is calculated as

$$\frac{\partial}{\partial n_{\mathrm{b}}}\left(\frac{1-n_{\mathrm{b}}}{\phi_{\mathrm{g}}}\right)^{1/3} = -\frac{r_{\mathrm{in}}}{3r_0}\frac{\partial}{\partial n_{\mathrm{b}}}\left[\log\left(\frac{\phi_{\mathrm{g}}}{1-n_{\mathrm{b}}}\right)\right]. \qquad \text{(S224)}$$

It is useful to use the relationship

$$\sigma b^2 = \frac{2n_{\mathrm{l}}}{4\pi r_{\mathrm{in}}^2} = \frac{r_0^2 \eta_{\mathrm{m}}^{2/3}}{r_{\mathrm{in}}^2}. \qquad \text{(S225)}$$

The relaxation time $\Delta\tau_{\mathrm{r}}$ per unit extrusion step is calculated by substituting the three types of solutions derived in sec. S5.5.1 into eq. (S94):

i) For the case of $s < \Lambda^{3/5}$ or $\Lambda > 1$, the derivatives of the osmotic pressure $\Pi_{\mathrm{in}}$ and the radius $r_{\rm in}$ are calculated as

$$\frac{\partial}{\partial n_{\rm b}}\left(\frac{\Pi_{\rm in}b^3}{k_{\rm B}T}\right)\bigg|_{V_{\rm in}} = -\frac{3}{2}\left(\frac{2G_0b^3}{\phi_0^{1/3}k_{\rm B}T}\right)^{6/5}\frac{(1+2\alpha_{\rm cg}(0)v_0\tau_{\rm ex}T(t))^{6/5}}{((1+\alpha_{\rm cg}(0))^2-2\chi_0)^{1/5}}\frac{1}{(1-n_{\rm b})^{13/5}}, \tag{S226}$$

$$\frac{\partial}{\partial n_{\rm b}}\left(\frac{1-n_{\rm b}}{\phi_{\rm g}}\right)^{1/3} = -\frac{3}{5}\frac{r_{\rm in}}{r_0(1-n_{\rm b})}, \tag{S227}$$

by substituting eq. (S162) into eqs. (S223) and (S224), respectively. The ratio $h/r_{\rm in}$ is calculated as

$$\frac{h}{r_{\rm in}} = \frac{1}{3}\left(\frac{G_0b^3(1+2\alpha_{\rm cg}v_0\tau_{\rm ex}T(t))}{3\eta_{\rm m}^{4/3}\phi_0^{1/3}k_{\rm B}T}\right)^{1/3}\left(\frac{(1+\alpha_{\rm cb}(0))^2-2\chi_0}{(1+\alpha_{\rm cg}(0))^2-2\chi_0}\right)^{1/3}\frac{n_{\rm b}}{1-n_{\rm b}}, \tag{S228}$$

by substituting eq. (S164) into eq. (S50). By substituting eqs. (S226) - (S228) into (S94), the relaxation time $\Delta\tau_{\rm r}$ per unit extrusion step is derived as

$$\frac{\Delta\tau_{\rm r}}{\tau_{\rm r}} = \frac{2}{15\cdot 6^{1/3}}\frac{n_{\rm b}(1-n_{\rm b})^{3/5}}{(1-\phi_{\rm b})^2}\frac{\eta_{\rm m}^{2/9}(\phi_0^{1/3}k_{\rm B}T)^{13/15}}{(2G_0b^3(1+2\alpha_{\rm cg}v_0\tau_{\rm ex}T(t)))^{13/15}}\frac{((1+\alpha_{\rm cb}(0))^2-2\chi_0)^{1/3}}{((1+\alpha_{\rm cg}(0))^2-2\chi_0)^{2/15}}. \tag{S229}$$

ii) For the case of $\Lambda^{3/5} < s < \Lambda^{-9/5}$ and $\Lambda < 1$, the derivatives of the osmotic pressure $\Pi_{\rm in}$ and the radius $r_{\rm in}$ is calculated as

$$\frac{\partial}{\partial n_{\rm b}}\left(\frac{\Pi_{\rm in}b^3}{k_{\rm B}T}\right)\bigg|_{V_{\rm in}} = -\frac{1}{3}\left(\frac{3G_0b^3}{\phi_0^{1/3}k_{\rm B}T}\right)^{13/10}\frac{(1+2\alpha_{\rm cg}(0)v_0\tau_{\rm ex}T(t))^{13/10}}{(6\eta_{\rm m})^{1/10}((1+\alpha_{\rm cb}(0)^2-2\chi_0))^{1/5}}\frac{1}{n_{\rm b}^{3/10}(1-n_{\rm b})^{23/10}} \tag{S230}$$

$$\frac{\partial}{\partial n_{\rm b}}\left(\frac{1-n_{\rm b}}{\phi_{\rm g}}\right)^{1/3} = -\frac{9}{10}\frac{r_{\rm in}}{r_0}\frac{2n_{\rm b}-1}{n_{\rm b}(1-n_{\rm b})}, \tag{S231}$$

by substituting eq. (S165) into eqs. (S223) and (S224), respectively. In this regime, the second term of eq. (S223) is negligible. The ratio $h/r_{\rm in}$ is calculated as

$$\frac{h}{r_{\rm in}} = \frac{1}{3}\left(\frac{G_0 b^3(1+2\alpha_{\rm cg}v_0\tau_{\rm ex}T(t))}{2\eta_{\rm m}^{4/3}\phi_0^{1/3}k_{\rm B}T}\right)^{1/2}\left(\frac{n_{\rm b}}{1-n_{\rm b}}\right)^{1/2} \tag{S232}$$

by substituting eq. (S167) into eq. (S50). By substituting eqs. (S230) - (S232) into (S94), the relaxation time per unit extrusion step thus is derived as

$$\frac{\Delta\tau_{\rm r}}{\tau_{\rm r}} = \frac{9}{10\cdot 6^{2/5}}\left(\frac{\phi_0^{1/3}k_{\rm B}T}{3G_0b^3}\right)^{4/5}\frac{\eta_{\rm m}^{2/15}((1+\alpha_{\rm cb}(0))^2-2\chi_0)^{1/5}}{(1+2\alpha_{\rm cg}(0)v_0\tau_{\rm ex}T(t))^{4/5}}\frac{(1-n_{\rm b})^{4/5}(2n_{\rm b}-1)}{n_{\rm b}^{1/5}(1-\phi_{\rm b})^2}. \tag{S233}$$

iii) For the case of $s > \Lambda^{-9/5}$ and $\Lambda < 1$, the derivatives of the osmotic pressure $\Pi_{\rm in}$ and the radius $r_{\rm in}$ is calculated as

$$\frac{\partial}{\partial n_{\rm b}}\left(\frac{\Pi_{\rm in}b^3}{k_{\rm B}T}\right)\bigg|_{V_{\rm in}} = -\frac{1}{2}\left(\frac{2G_0b^3}{\phi_0^{1/3}k_{\rm B}T}\right)^{8/5}\frac{(1+2\alpha_{\rm cg}(0)v_0\tau_{\rm ex}T(t))^{8/5}}{(6\eta_{\rm m}^{4/3})^{2/5}((1+\alpha_{\rm cb}(0))^2-2\chi_0)^{3/5}}\frac{1}{(1-n_{\rm b})^{13/5}} \tag{S234}$$

$$\frac{\partial}{\partial n_{\rm b}}\left(\frac{1-n_{\rm b}}{\phi_{\rm g}}\right)^{1/3} = -\frac{3}{5}\frac{r_{\rm in}}{r_0}\frac{1}{1-n_{\rm b}}, \tag{S235}$$

by substituting eq. (S168) into eqs. (S223) and (S224), respectively. In this regime, the second term of eq. (S223) is negligible. The ratio $h/r_{\rm in}$ is calculated as

$$\frac{h}{r_{\rm in}} = \frac{1}{9}\frac{G_0b^3(1+2\alpha_{\rm cg}v_0\tau_{\rm ex}T(t))}{3\eta_{\rm m}^{4/3}\phi_0^{1/3}k_{\rm B}T((1+\alpha_{\rm cb}(0))^2-2\chi_0)^{2/3}}\frac{n_{\rm b}}{1-n_{\rm b}} \tag{S236}$$

by substituting eq. (S170) into eq. (S50). By substituting eqs. (S234) - (S236) into (S94), the relaxation time per unit extrusion step thus is derived as

$$\frac{\Delta\tau_{\rm r}}{\tau_{\rm r}} = \frac{6^{2/5}}{15}\frac{n_{\rm b}(1-n_{\rm b})^{3/5}}{(1-\phi_{\rm b})^2}\left(\frac{\phi_0^{1/3}k_{\rm B}T}{2G_0b^3}\right)^{3/5}\frac{\eta_{\rm m}^{-2/15}}{(1+2\alpha_{\rm cg}(0)v_0\tau_{\rm ex}T(t))^{3/5}}\frac{1}{((1+\alpha_{\rm cb}(0))^2-2\chi_0)^{1/15}}. \tag{S237}$$

### S6 Supplementary Discussion

#### S6.1 Sparkler is not assembled at low condensin concentrations

Experimentally, entangled DNA strands in Xenopus egg extract can be observed by changing the concentration $\phi_{\rm c0}$ of condensin complexes. In our theory, it corresponds to study the dependence of the dynamics of entangled DNA strands on condensin occupancy $\alpha_{\rm c0}$ $(= k_{\rm on}\tau_{\rm ex}\phi_{\rm c0})$ . Our theory predicts that the peripheral brush region is not stable if the condensin occupancy $\alpha_{\rm c0}$ is too small, see the purple line in Fig. S7**a**. In a window of the condensin occupancy $\alpha_{\rm c0}$, the entangled DNA strands form a uniform swollen brush at the peripheral region in the steady state, see the brown line in Fig. S7**a**. However, this state may not be the true steady state because the branching of DNA loops due to the simultaneous loading of condensin complexes, which is not taken into account in our theory, becomes significant in a long time scale. The symmetry breaking happens with larger condensin occupancy $\alpha_{\rm c0}$, see Fig. 3a in the main article. These results are consistent with the steady state analysis, $dN_{\rm bl}/dt = 0$ and $dr_{\rm in}/dt = 0$, see Fig. S7**a** and **b**. The size of the central gel region decreases with increasing the condensin occupancy $\alpha_{\rm c0}$ to relax the DNA tension developed by the loop extrusion process, see the magenta line in Fig. S7**c**. These predictions may be experimentally accessible.

#### S6.2 Relative DNA volume fraction between the central and peripheral regions reflects the entanglement of DNA strands

The shear modulus $G_0$ of the central gel region reflects the entanglement of DNA strands. Although it is, in principle, experimentally accessible by measuring the rheology of entangled DNA strands, the estimate of its value is difficult because the DNA strands are not entangled in an equilibrium solution. We thus study the dependence of the dynamics of entangled DNA strands on the shear modulus $G_0$.

For the case of the fast relaxation, the radius $r_{\rm in}$ of the central gel region is determined by the balance of the osmotic pressure $\Pi_{\rm sol}(\phi_{\rm g}) + \Delta\Pi_{\rm g}$ and the elastic force $\Pi_{\rm ela}$, which is proportional to the shear modulus $G_0$. Both of the DNA volume fractions in the central $\phi_{\rm g}$ and peripheral $\phi_{\rm b}$ regions decreases with decreasing the shear modulus $G_0$, see Figs. 3 in the main article and Figs. S7**a** and **b**. In a good solvent, the ratio $\phi_{\rm g}/\phi_{\rm b}$ is represented as eq. (S164) for cases in which the osmotic pressure $\Pi_{\rm sol}(\phi_{\rm g})$ of the central gel region is larger than the osmotic pressure contribution $\Delta\Pi_{\rm g}$ of the peripheral brush region. Eq. (S164) suggests that the ratio $\phi_{\rm g}/\phi_{\rm b}$ decreases with decreasing the shear modulus $G_0$. In typical experiments, the DNA concentration in the central region is smaller than that in the peripheral region before most part of the DNA in the peripheral region vanishes (Shintomi and Hirano 2021). This implies that the shear modulus $G_0$ of the mouse sperm DNA used in the experiments was relatively small.

Interestingly, the DNA volume fraction $\phi_{\rm b}$ in the central gel region is a non-monotonic function of time for cases in which the shear modulus $G_0$ is very small, see the light green line in Fig. S7**b**. In such cases, the ratio $\phi_{\rm g}/\phi_{\rm b}$ is represented as eq. (S167) for $2h/r_{\rm in} < 1$ and eq. (S170) for $2h/r_{\rm in} > 1$. These results may be accessible by experimentally measure the ratio of DNA concentration between the central and peripheral region as a function of time.

The swollen state of the central gel region becomes unstable at a value $\phi_{\rm g}^{\rm sp1}$ of DNA volume fraction and shows the volume phase transition to the condensed state, see Fig. 3c in the main article. A window of the fraction $s$ of domains of swollen DNA loops in the peripheral region is not allowed at the phase transition because of the volume phase transition, see Fig. 4**b** in the main article. The DNA volume fraction $\phi_{\rm g}^{\rm sp1}$ at the stability limit increases with decreasing the shear modulus $G_0$. When the shear modulus $G_0$ is small enough, the fraction $s$ of swollen domains continuously decreases from unity to zero as the DNA volume fraction $\phi_{\rm b}$ at the phase transition increases, see Fig. S7**c**. In these cases, DNA in the central gel region is condensed only after the phase separation of the peripheral region happens, see Fig. 5**b** in the main article.

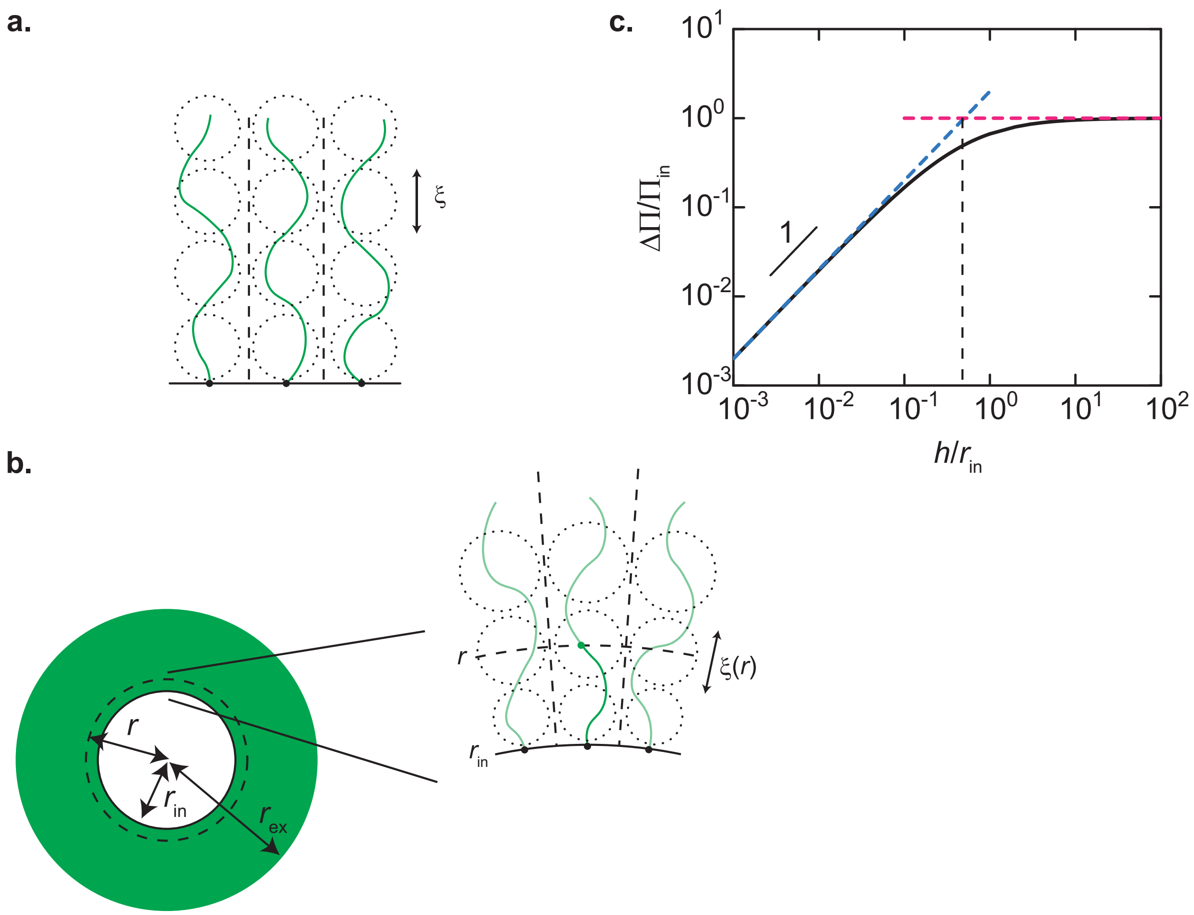


Figure S1: **Scaling theory of polymer brush. a.** A polymer brush on a planer surface. Grafted polymers are shown by green lines. The correlation length $\xi$ of polymers is equal to the distance between the grafting points (black dots). The dotted circles are blobs of size $\xi$. **b.** A polymer brush on a spherical surface of radius $r_{\rm in}$. The position in the brush is represented by the distance $r$ from the center of the sphere (dotted line). **c.** The pressure $\Delta\Pi$ applied to the surface by the brush shown as function of the brush height $h$. The solid line is the osmotic pressure predicted by the Daoud-Cotton theory, eq. (S10). The cyan and maganta broken lines are derived by using the asymptotic forms, eqs. (S12) and (S13), respectively.

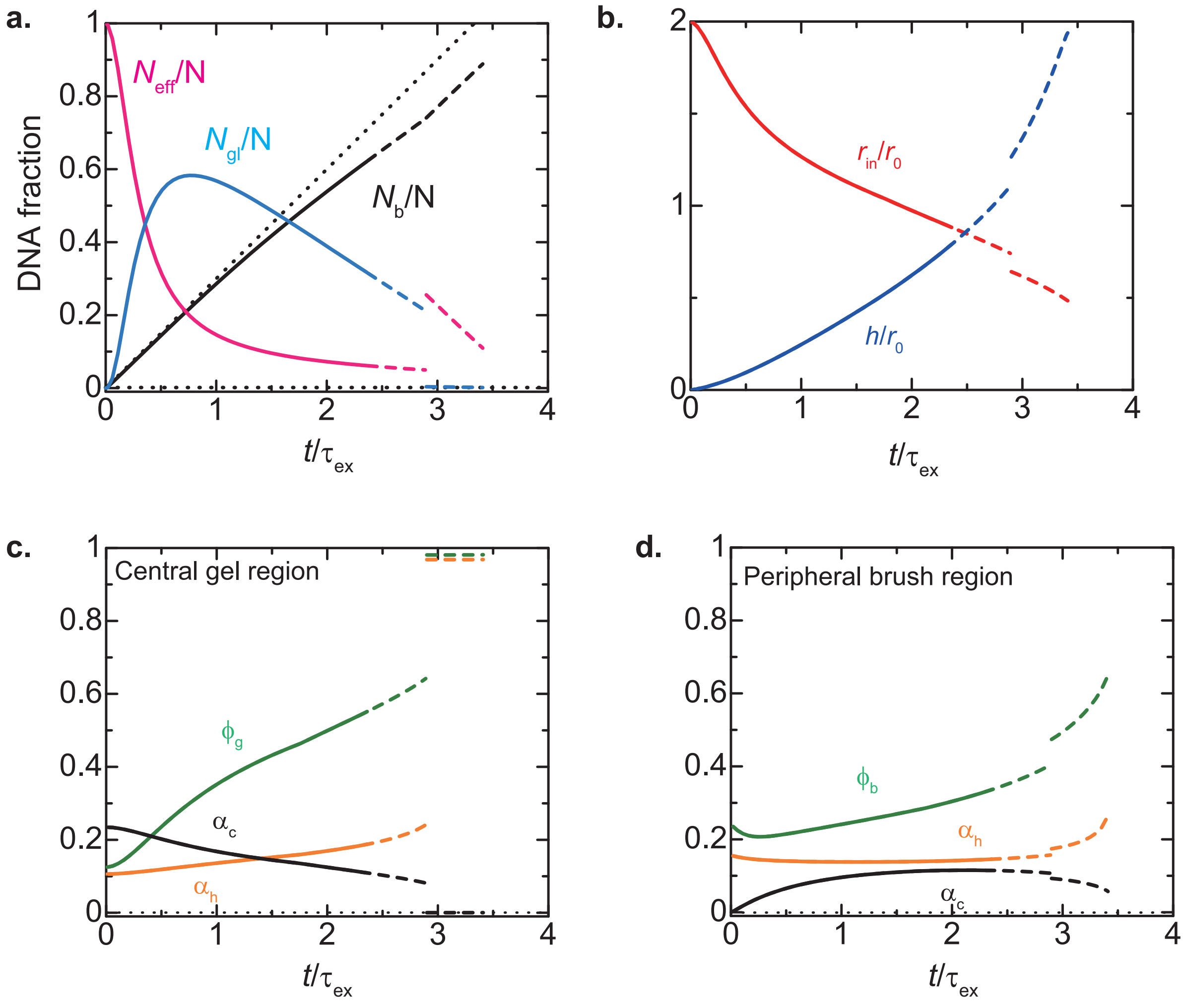


Figure S2: **Loop extrusion dynamics with fast relaxation:** The structural parameters of entangled DNA strands are shown as a function of time $t$ for cases in which the relaxation is faster than loop extrusion. **a.** The fractions of DNA in the peripheral brush region $N_b/N$ (black solid and broken lines) as well as the elastically effective chains $N_{eff}/N$ (magenta lines) and elastically ineffective chains $N_{gl}/N$ (cyan lines) in the central gel region. **b.** The radius $r_{in}$ of the central gel region (red line) and the thickness $h$ of the peripheral brush region. **c.** The DNA volume fraction $\phi_g$ (green line), the condensin occupancy (black), and the linker histone occupancy (orange) in the central gel region. **d.** The DNA volume fraction $\phi_g$ (green line), the condensin occupancy (black), and the linker histone occupancy (orange) in the peripheral brush region. The values of parameters used for this calculation are summarized in Table 1 in the main article.

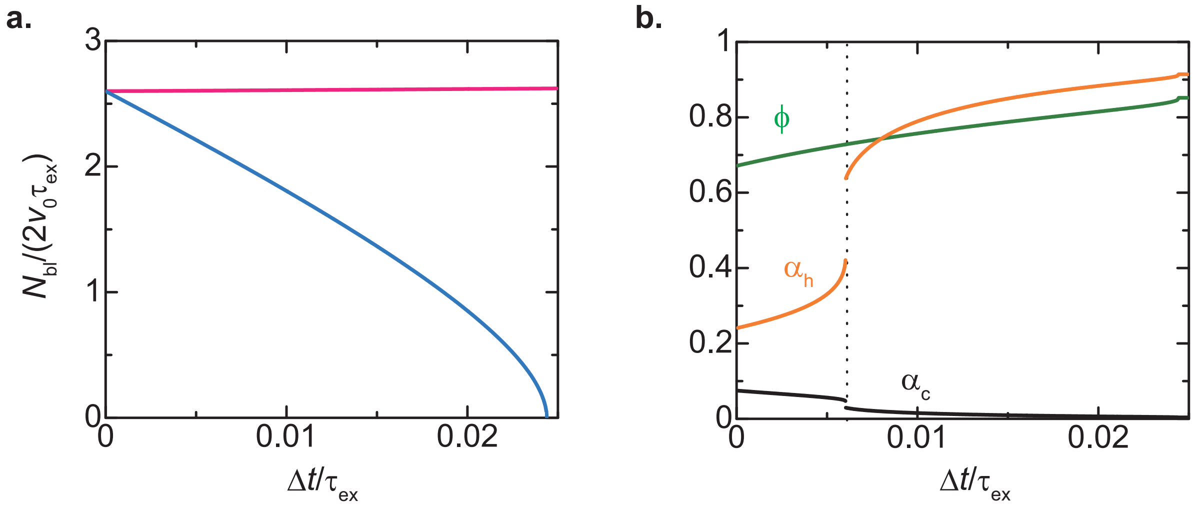


Figure S3: **Binding transition: a.** The numbers $N_{\mathrm{bl}}$ of DNA units in a swollen loop rich in condensin complexes (magenta) and a condensed loop rich in linker histones (cyan) are shown as a function of time $\Delta t$ after the phase separation. **b.** The DNA volume fraction $\phi$ (green), the linker histone occupancy $\alpha_{\mathrm{h}}$ (orange), the condensin occupancy $\alpha_{\mathrm{c}}$ (black) are shown as a function of time $\Delta t$. We used $\epsilon_{\mathrm{h}} - \mu_{\mathrm{h}} = 1.95$, which is larger than the critical value $\epsilon_{\mathrm{h}}^{\mathrm{c}} - \mu_{\mathrm{h}}^{\mathrm{c}} (= 2 + \log(1 - \alpha_{\mathrm{c0}}(1 - 2/\chi_{\mathrm{m}})))$. The values of other parameters are summarized in Table 1.

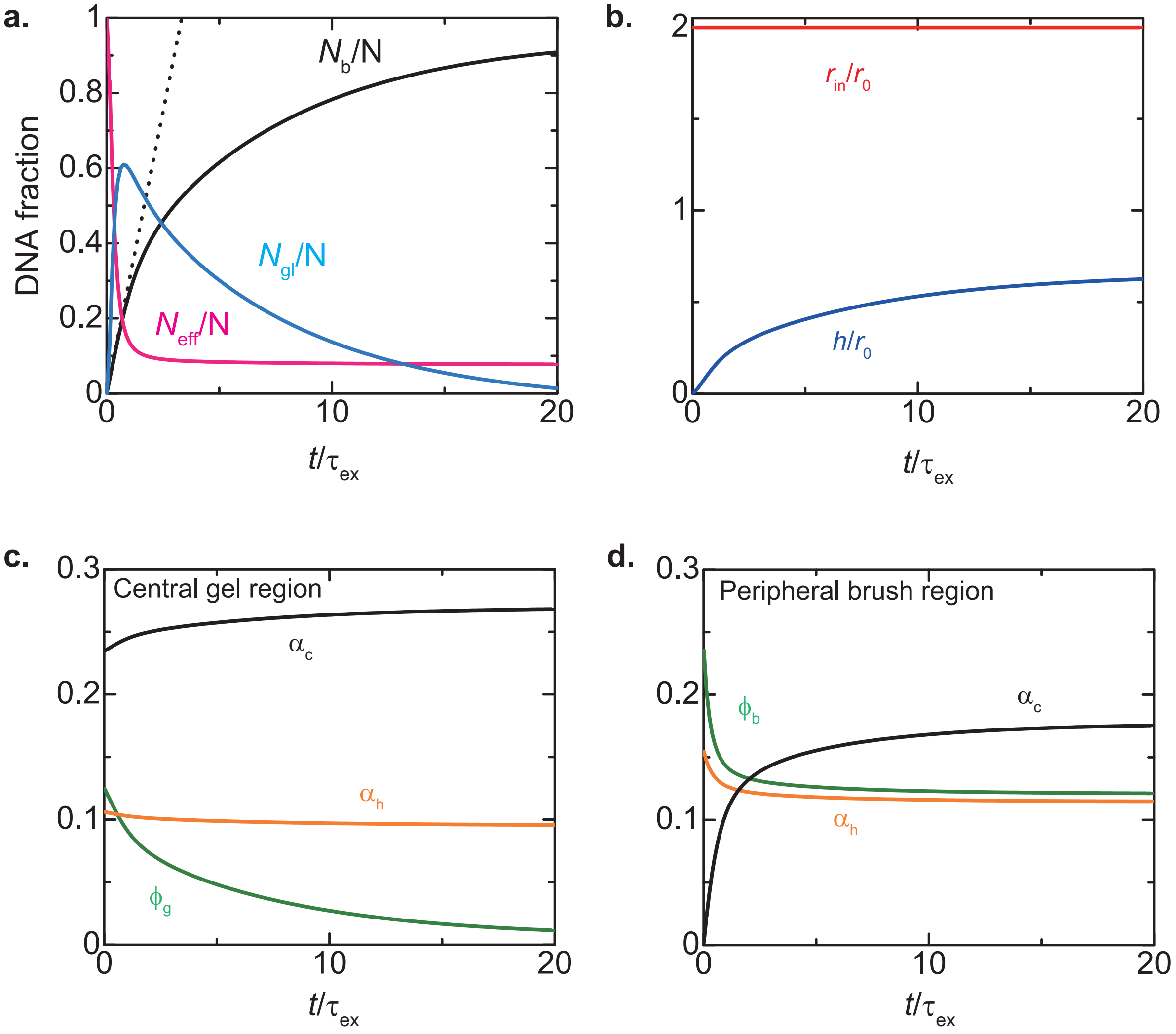


Figure S4: **Loop extrusion dynamics with slow relaxation:** The structural parameters of entangled DNA strands are shown as a function of time $t$ for cases in which the relaxation is slower than loop extrusion until the DNA tension stops the loop extrusion. **a.** The fractions of DNA in the peripheral brush region $N_b/N$ (black solid and broken lines) as well as the elastically effective chains $N_{eff}/N$ (magenta lines) and elastically ineffective chains $N_{gl}/N$ (cyan lines) in the central gel region. **b.** The radius $r_{in}$ of the central gel region (red line) and the thickness $h$ of the peripheral brush region. **c.** The DNA volume fraction $\phi_g$ (green line), the condensin occupancy (black), and the linker histone occupancy (orange) in the central gel region. **d.** The DNA volume fraction $\phi_g$ (green line), the condensin occupancy (black), and the linker histone occupancy (orange) in the peripheral brush region. The values of parameters used for this calculation are summarized in Table 1 in the main article.

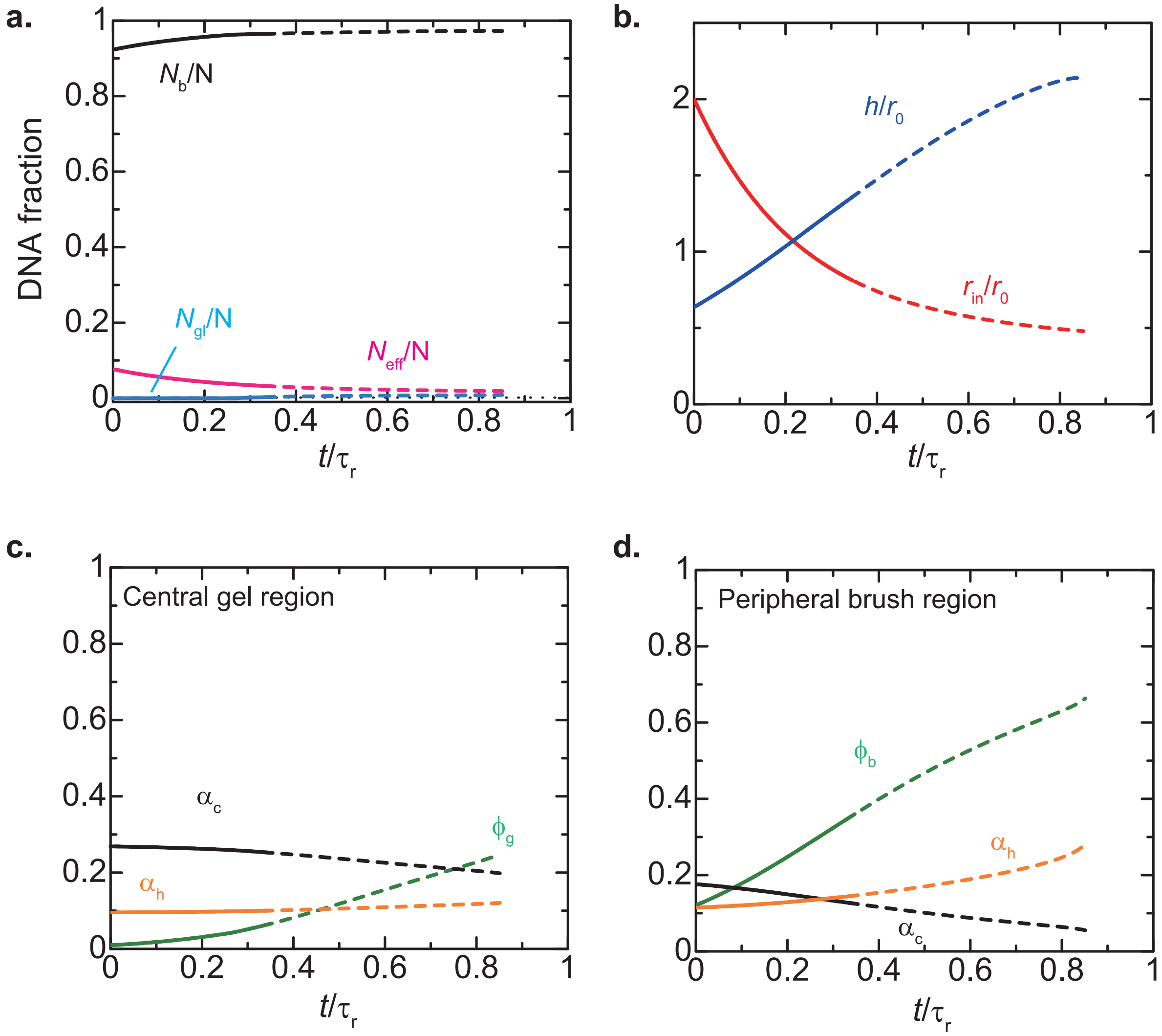


Figure S5: **Relaxation dynamics with slow relaxation:** The structural parameters of entangled DNA strands are shown as a function of time $t$ for cases in which the relaxation is slower than loop extrusion after the DNA tension stops the loop extrusion. **a.** The fractions of DNA in the peripheral brush region $N_{\mathrm{b}}/N$ (black solid and broken lines) as well as the elastically effective chains $N_{\mathrm{eff}}/N$ (magenta lines) and elastically ineffective chains $N_{\mathrm{gl}}/N$ (cyan lines) in the central gel region. **b.** The radius $r_{\mathrm{in}}$ of the central gel region (red line) and the thickness $h$ of the peripheral brush region. **c.** The DNA volume fraction $\phi_{\mathrm{g}}$ (green line), the condensin occupancy (black), and the linker histone occupancy (orange) in the central gel region. **d.** The DNA volume fraction $\phi_{\mathrm{g}}$ (green line), the condensin occupancy (black), and the linker histone occupancy (orange) in the peripheral brush region. The values of parameters used for this calculation are summarized in Table 1 in the main article.

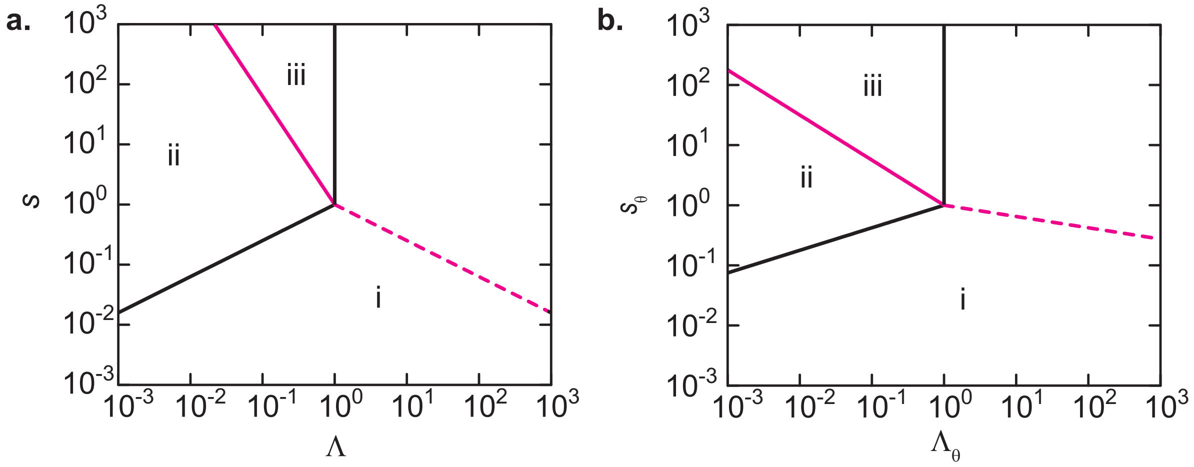


Figure S6: **Regime diagram for fast relaxation limit:** The ranges of the parameters $\Lambda$ and $s$ for three types of solutions, i, ii, and iii, of the pressure balance equation are shown for the good (**a**) and $\theta$ (**b**) solvent conditions. The black lines show the condition of $\Pi_{\mathrm{sol}}(\phi_{\mathrm{g}}) = \Delta\Pi_{\mathrm{g}}$ and the magenta lines show the condition of $2h/r_{\mathrm{in}} = 1$. The solution does not change at the magenta broken line because the contribution of the peripheral region is small.

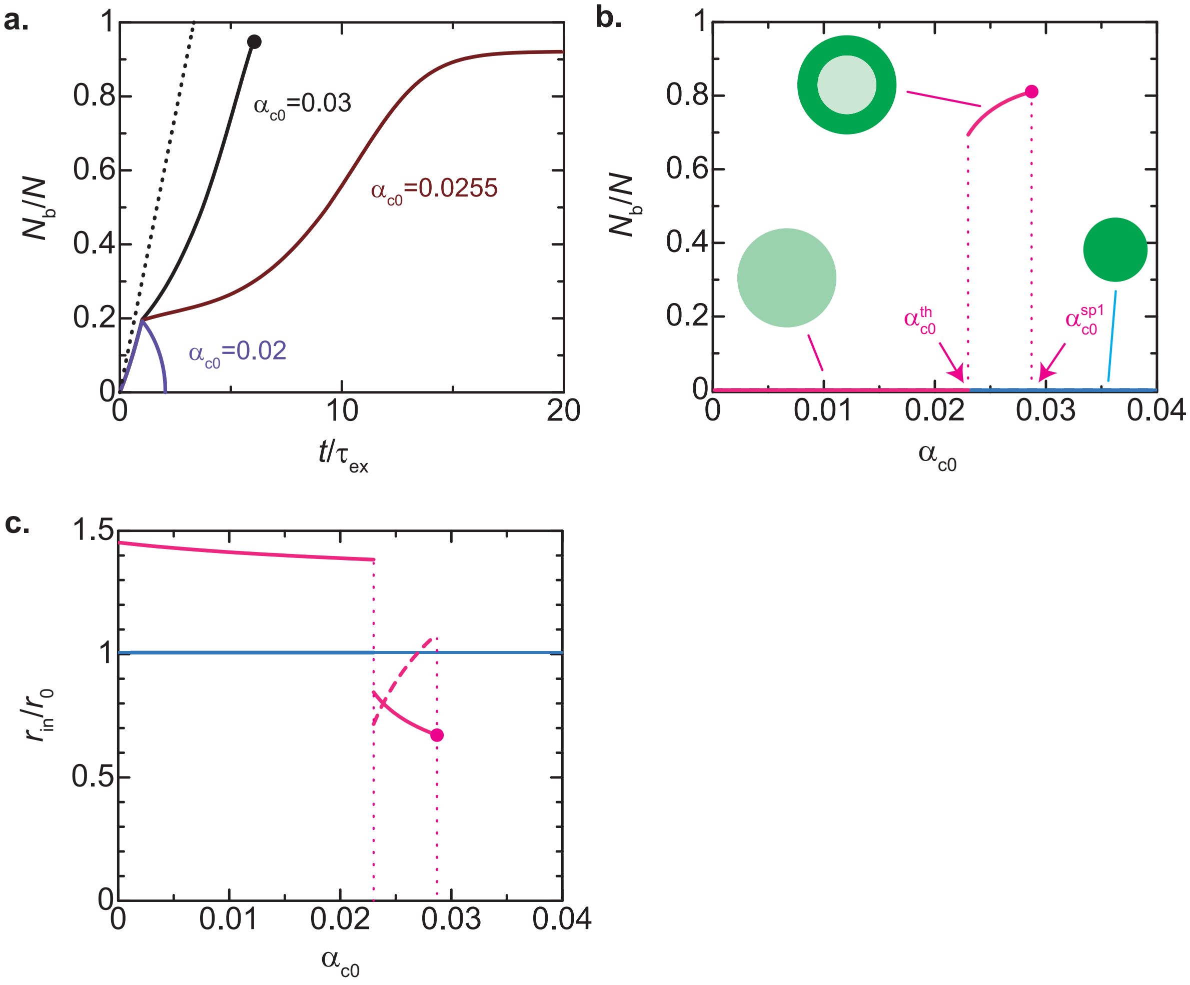


Figure S7: **Dependence of loop extrusion dynamics on condensin occupancy $\alpha_{c0}$.** **a.** The fraction $N_b/N$ of DNA units in the peripheral brush region is shown as a function of time for $\alpha_{c0} = 0.03$ (black), 0.0255 (brown), and 0.02 (purple). The dotted line represents the case of ideal loop extrusion without deceleration by DNA tension and condensin unloading. The fraction $N_b/N$ of DNA units in the peripheral brush region (**b**) and the radius of the central gel region (**c**) are shown as a function of the condensin occupancy $\alpha_{c0}$ as derived by the steady state analysis, $dN_{bl}/dt = 0$ and $dr_{in}/dt = 0$. The broken line is the thickness of the peripheral brush region. The magenta and cyan lines are the solutions for the swollen and condensed states, respectively. The values of parameters used for these calculations are summarized in Table 1 in the main article, except for the condensin occupancy $\alpha_{c0}$.

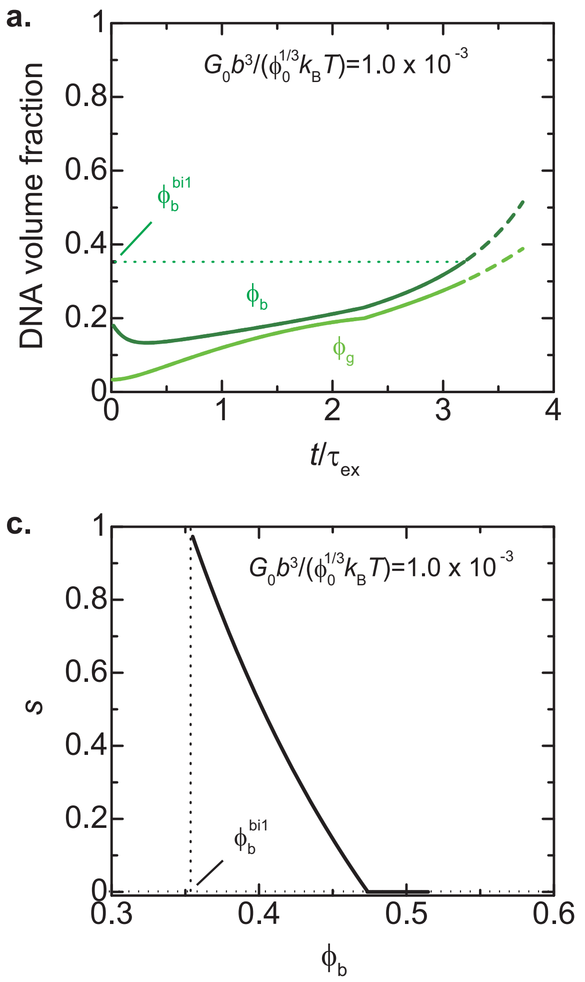


Figure S8: **Dependence of loop extrusion dynamics on the shear modulus $G_0$.** The DNA volume fractions in the central (light green) and peripheral (deep green) regions are shown as a function of time t for $G_0b^3/(\phi_0^{1/3}k_BT) = 1.0x10^{-3}$ (**a**) and $1.0x10^{-4}$ (**b**). $\phi_b^{bi1}$ is the DNA volume fraction at the binodal line. The peripheral brush region is a uniformly swollen state for $\phi_b < \phi_b^{bi1}$ (solid line) and can show phase separation at a DNA volume fraction $\phi_b > \phi_b^{bi1}$ (broken line). **c.** The fraction $s$ of the domains of swollen loops rich in condensin complexes is shown as a function of the DNA volume fraction $\phi_b$ at which the phase separation occurs. The values of parameters used for these calculations are summarized in Table 1 in the main article, except for $G_0b^3/(\phi_0^{1/3}k_BT)$.
